# Genomic estimation of quantitative genetic parameters in wild admixed populations

**DOI:** 10.1101/2021.09.10.459723

**Authors:** Kenneth Aase, Henrik Jensen, Stefanie Muff

## Abstract

1. Heritable genetic variation among free-living animals or plants is essential for populations to respond to selection and adapt. It is therefore important to be able to estimate additive genetic variance *V*_A_, which can be obtained using a generalized linear mixed model known as the *animal model*. An underlying assumption of the standard animal model is that the study population is genetically unstructured, which is often unrealistic. In fact, admixture might be the norm rather than the exception in the wild, like in geographically structured populations, in the presence of (im)migration, or in re-introduction and conservation contexts. Unfortunately, animal model estimators may be biased in such cases. So-called *genetic group* animal models that account for genetically differentiated subpopulations have recently become popular, but methodology is currently only available for cases where relatedness among individuals can be estimated from pedigrees.
2. To ensure that genetic group animal models with heterogeneous *V*_A_ remain applicable to populations with genomic data but no pedigrees, there is a clear need to generalize these models to the case when exclusively genomic data is available. We therefore introduce such methodology for wild admixed systems by extending methods that were recently suggested in the context of plant breeding. Our extension relaxes the limiting assumptions that currently restrict their use to artificial breeding setups.
3. We illustrate the usefulness of the extended genomic genetic groups animal model on a wild admixed population of house sparrows resident in an island system in Northern Norway, where genome-wide data on more than 180 000 single nucleotide polymorphisms (SNPs) is available to derive genomic relatedness. We compare our estimates of quantitative genetic parameters to those derived from a corresponding pedigree-based genetic groups animal model. The satisfactory agreement indicates that the new method works as expected.
4. Our extension of the very popular animal model ensures that the upcoming challenges with increasing availability of genomic data for quantitative genetic studies of wild admixed populations can be handled. To make the method widely available to the scientific community, we offer guidance in the form of a tutorial including step-by-step instructions to facilitate implementation.

## Introduction

A major goal in quantitative genetics is to disentangle environmental and genetic contributions to a phenotype within a study population (Falconer & Mackay, 1996; Lynch & Walsh, 1998; Charmantier *et al*., 2014). The partitioning of the phenotypic variance in a population into additive genetic and environmental components is of particular interest, as the additive genetic variance (*V*_A_) is a crucial determinant of the rate by which phenotypes may respond to selection across generations. Simply speaking, the larger the *V*_A_ of a focal trait, the faster a population is able to respond to a given strength of selection, leading to a higher rate of adaptive evolution (Walsh & Lynch, 2018). The magnitude of *V*_A_ can thus be a determinant for how quickly and well wild populations may adapt to changing environmental conditions. Insight into such mechanisms is particularly important in conservation and wildlife management (Frankham *et al*., 2010; Carlson *et al*., 2014), and also highly relevant in breeding programs (Nyquist & Baker, 1991).

One well-established statistical tool to estimate genetic parameters like *V*_A_ is the linear mixed effects model known as the *animal model* (Henderson, 1984; Kruuk, 2004; Wilson *et al*., 2010). The animal model relies on knowledge about genetic relatedness between every pair of individuals (stored in a genetic relatedness matrix), which should reflect how similar they are at causal loci within their genomes (Weir *et al*., 2006; Speed & Balding, 2015). We denote the additive genetic impact on an individual’s phenotype as its *genetic value* (or breeding value). Importantly, the *V*_A_ estimated by the animal model is relative to the so-called *base population*, while the definition of the base population itself depends on how one chooses to measure genetic relatedness (Legarra,2016). The basic animal model relies on the assumption that there is a single, homogeneous base population without genetic substructures, and thus the model may produce biased estimates of *V*_A_ when base populations consist of individuals with systematically different genetic parameters (Wolak & Reid, 2017).

Genetic substructures are present in populations where breeding occurs between individuals from genetically divergent populations (*e.g*., due to historical isolation). The resulting gene flow is known as *admixture* (Tang *et al*., 2005), and is frequently encountered in the contexts of crossbreeding (Toosi *et al*., 2010) and hybridization (Grabenstein & Taylor, 2018). Instances of admixture in wild systems can arise in various ways, both naturally and human-induced (Lenormand, 2002). Populations can, for example, receive immigrants from distant populations (*e.g*., Wolak & Reid, 2017), and metapopulations are subject to ongoing admixture between subpopulations through dispersal (*e.g*., Saatoglu *et al*., 2021). In conservation contexts, reintroduction schemes may involve translocating individuals from elsewhere to reinforce endangered populations (*e.g*., Ranke *et al*., 2020). Admixture is indeed expected to be relevant in most wild populations, because some level of dispersal and gene flow occurs into all populations except the few that are completely isolated (Bowler & Benton, 2005; Ronce, 2007). Consequently, the aforementioned homogeneity assumption of the basic animal model might frequently be violated in wild systems, potentially producing biased estimates. Thus, to accurately estimate the genetic parameters of wild populations, admixture should be taken into account.

Fortunately, animal models have been extended to account for admixture by partitioning the base population into *genetic groups* (Quaas, 1988; Wolak & Reid, 2017), where *group-specific* mean genetic values, and optionally *group-specific* additive genetic variances, are allowed. If the genetic groups only differ in the mean breeding value, but not in their *V*_A_, then allowing for group-specific mean genetic values is sufficient to correct the aforementioned biases. However, in some applications it is more realistic to also allow for group-specific additive genetic variances, which comes at the cost of higher demands on both computation time and data (Muff *et al*., 2019).

In genetic group animal models (or simply genetic group models) we will differentiate between *purebred* individuals whose genomes belong to a single genetic group, and *admixed* individuals whose genomes are a mix of contributions from two or more genetic groups. As an example, when a local population receives immigrants, genetic group models may assign known natives as purebred in one genetic group, and known immigrants as purebred in a second genetic group, thereby explicitly incorporating the base population’s genetic structure into the model (Wolak & Reid, 2017). The admixed set of individuals then contains all descendants from native/immigrant matings, and we track the respective admixture proportions in each individual in later generations, for example by following the pedigree. Some genetic group animal models also include segregation variance, an additional source of variance that emerges under admixture. These variances are mostly relevant in artificial breeding scenarios, or when the number of loci impacting the phenotype is very small (Slatkin & Lande, 1994). Fortunately, segregation variances are negligible under the assumption of the infinitesimal model (Bulmer, 1971), which is standard for complex traits in humans, animal and plant breeding, as well as natural populations (Hill & Kirkpatrick, 2010; Hill, 2012).

Genetic relatedness estimates used for basic animal models have traditionally been derived from pedigrees. Similarly, pedigree-based genetic group extensions of the animal model that account for admixture are well-established in the plant and animal breeding literature (Schaeffer, 1991; Lo *et al*., 1993; García-Cortés & Toro, 2006), and have recently found their way into applications for wild animal systems (Wolak & Reid, 2017; Muff *et al*., 2019; Reid *et al*., 2021). Pedigrees can produce estimates of relatedness and global ancestry (the overall proportion of a genome belonging to some genetic group) that are true on expectation (given a correct pedigree), by tracing all matings and applying the usual Mendelian rules of inheritance (Wright, 1922). Pedigree-based animal models have favorable computational properties due to the sparseness of the relatedness matrices and their inverses (Henderson, 1984), but these methods also have some inherent weaknesses. Realized genetic relatedness and global ancestry, for example, often differ greatly from the expected value derived from pedigrees (Hill & Weir, 2011), and pedigrees are often error-prone or incomplete (Keller *et al*., 2001; Ponzi *et al*., 2019).

Animal models that use single nucleotide polymorphisms (SNPs) to derive relatedness (Stanton-Geddes *et al*., 2013; Speed & Balding, 2015; Wang *et al*., 2017) have become increasingly popular due to improved genotyping technologies and decreased costs (Andrews *et al*., 2018). Provided that the number of genotyped loci is high enough, such *genomic* animal models, which rely on *genomic* relatedness matrices (GRMs), generally provide more accurate estimates of quantitative genetic parameters than pedigree-based animal models (Bérénos *et al*., 2014; Gienapp *et al*., 2017). On the other hand, the genomic approach is more computationally challenging compared to the pedigree-based version, as GRMs are dense.

A current drawback of genomic animal models is that it is still unclear how to handle genetic groups, in particular in the case of wild admixed populations with heterogeneous *V*_A_. Recently, Rio *et al*. (2020b) developed a heterogeneous *V*_A_ genetic group animal model with a genomic framework (denoted MAGBLUP-RI) for use in plant breeding. MAGBLUP-RI relies on knowledge about *local* ancestry, that is, the ancestry of each individual allele, indicating which genetic group that allele is descended from (Gravel, 2012). Through the use of local ancestry information, the model explicitly incorporates the fact that an admixed individual’s genome is a mosaic of ancestries from different genetic groups. Local ancestry can be inferred from genotype data (Geza *et al*., 2019; Schubert *et al*., 2020), and thus MAGBLUP-RI is a genetic group model that solely relies on genomic data. The main drawback of the MAGBLUP-RI method is that it assumes homozygosity at all loci. While this restriction might be justified in a plant breeding setup, it precludes use of the model on wild study systems, where heterozygous SNPs are common. Moreover, Rio *et al*. (2020b) allow for only two genetic groups, which is likely to be insufficient in fragmented systems in the wild.

In this paper (based on a master’s thesis by Aase, 2021) we overcome existing limitations of previous methodology and generalize *genomic* genetic group animal models such that they can be readily used to analyze wild admixed populations of diploid individuals with group-specific *V*_A_. To this end, we have extended MAGBLUP-RI so that it handles SNP data with heterozygous (rather than only homozygous) loci and any number of genetic groups. Additionally, we assume that segregation variances are negligible, although we also provide full model derivations for segregation terms in Supporting Information S1. As a proof of concept, we have applied the extended genomic genetic group animal model to a metapopulation of house sparrows (*Passer domesticus*), and compared the results to a pedigree-based, but otherwise equivalent, model proposed by Muff *et al*. (2019). All the steps of the implementation are detailed in a tutorial (Supporting Information S2).

## Materials and Methods

### The animal model and genetic group models

The simplest version of the animal model for a continuous phenotype *y_i_* of individual *i* in a population of size *N* is given by the mixed model

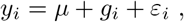

where the intercept *μ* is a fixed effect, and the genetic (or breeding) value *g_i_* and the independent residual term 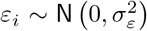 of individual *i* are random effects (Kruuk, 2004; Wilson *et al*., 2010). Additional fixed and random effects can be included to account for other (non-genetic) sources of covariance between observations, such as sex, time of measurement, or environmental effects (Kruuk & Hadfield, 2007). Fitting the model involves estimating the values of fixed effects and the parameters of the probability distributions of random effects. We give the vector of genetic values ***g*** = (*g*_1_,…, *g_N_*)^⊤^ the distribution 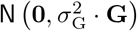, where the random effect parameter 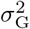 (to be estimated) is the *V*_A_ and **G** is a matrix containing known estimates of all pairwise genetic relatednesses between individuals. The mean genetic value (of zero) and the additive genetic variance 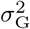 are assumed to be homogeneous within the base population. The entries of **G** can be estimated from a pedigree or from genomic data.

In the case where **G** is a GRM, that is, when its entries are derived from genomic (SNP) data, a widely used estimate was given by VanRaden (2008), where the genomic relatedness *G_ij_* between individuals *i* and *j* are

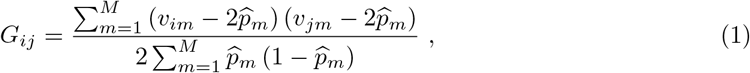

with *M* being the number of genotyped loci, *v_im_* and *v_jm_* being the number of copies of the alternate allele (usually the minor allele) at the *m*^th^ locus in diploid individuals *i* and *j* respectively (known from genotyping), and 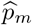 being the estimated allele frequency of the alternate allele at locus *m*. A relatedness measure is thus obtained by comparing the alleles of two individuals at every SNP, while weighting by the allele frequencies at each locus. Sharing rare alleles thus contributes more to relatedness estimates than sharing common alleles.

When the assumption that the genetic values in the entire base population are identically distributed is violated, the animal model should be extended accordingly. In the genetic groups animal model, admixture is accounted for by assuming each genetic group *r* = 1,…, *R* has a group-specific mean genetic value *γ_r_* (rather than mean 0) and (if desired) a group-specific *V*_A_ (Wolak & Reid, 2017; Muff *et al*., 2019). To this end, we partition ***g*** into group-specific genetic value vectors 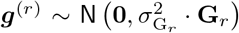, where 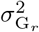 is the group-specific *V*_A_ and **G**_*r*_ is a group-specific relatedness matrix. Let the *total genetic value U_i_* be the sum of group-specific genetic values whose probability distributions depend on the group *r*. If a purebred individual in group *r* has mean genetic value *γ_r_*, denoted as the *genetic group effect* of group *r*, then 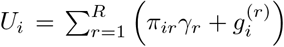, where *π_ir_* is the global ancestry of individual *i* with respect to group *r*. For identifiability reasons we add the constraint that one of the groups must have *γ_r_* = 0, and we label this group as the “reference group.” Thus, genetic group effects *γ_r_* can be estimated in the animal model by including estimates of global ancestries *π_ir_* as fixed effect covariates, while group-specific additive genetic variances 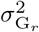 can be found by including the stochastic part of *U_i_*, namely 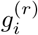, as random effects. The estimators for the global ancestries *π_ir_* and group-specific genetic relatedness matrices **G**_*r*_ depend on the genetic group method at hand (*e.g*., García-Cortés & Toro, 2006; Wolak & Reid, 2017; Muff *et al*., 2019; Rio *et al*., 2020b).

### Genomic genetic groups model in an inbred system

Rio *et al*. (2020b) previously included admixture in an animal model based on genomic data (which they label MAGBLUP-RI). They let the total genetic value *U_i_* be the sum of numeric contributions to the phenotype from each genotyped locus (*i.e*., a sum of allele effects). The contribution from each locus depends on its genotype and, importantly, on its *local ancestry*. All loci are assumed to be homozygous, and thus each has only two possible genotypes (both chromosomes have either the reference or alternate allele), as is the case in highly inbred plant breeding systems. In other words, the organisms under study are treated as *de facto* haploid. Further, only two genetic groups are assumed to exist. While the contribution of a locus given its local ancestry and genotype is considered deterministic, the genotype and local ancestry of each locus are themselves considered random variables, and therefore *U_i_* will also be a random variable.

Rio *et al*. (2020b) use the statistical properties of the total genetic values *U_i_* to derive the distribution of the group-specific genetic values 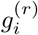. To derive the entries of the group-specific GRMs, the covariances between total genetic values *U_i_* must be considered. A central assumption here is that genotypes on different loci do not correlate, so no LD is present. In addition to the expected total genetic value of a purebred individual in group *r*, *γ_r_*, Rio *et al*. (2020b) define the parameters 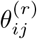 and 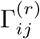. We can interpret 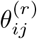 as the proportion of the alleles of individuals *i* and *j* that have the same local ancestry, averaged across all loci, that is, the overlap of *r*-descended regions in the two genomes. 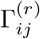 can be thought of as a genetic relatedness conditional on shared group membership in group *r*. The expected value and covariance for the total genetic values (when ignoring segregation variances) has then been shown to be

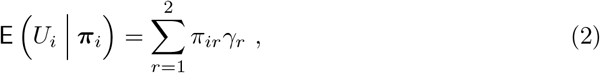

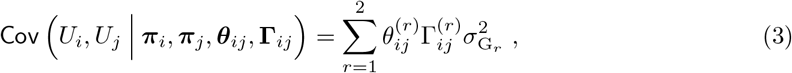

where 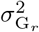 is the group-specific *V*_A_ for group *r* (Rio *et al*., 2020b). Thus, as in other genetic group models, the expected total genetic value of individual *i* is a weighted average of the mean expected value in the genetic groups, where the weights *π_ir_* are *i*’s global ancestry proportions in the groups *r* = 1, 2 (equation (2)). Furthermore, using equation (3) and estimators for 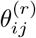 and 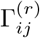, the entries of the group-specific GRMs were estimated by Rio *et al*. (2020b) as

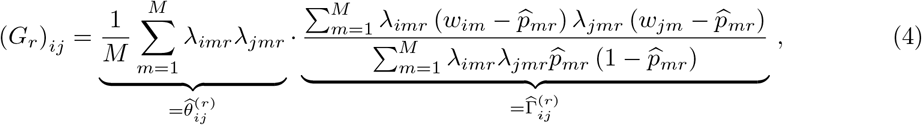

where *w_im_* = 1 indicates the presence of two copies of the alternate allele on individual *i*’s *m*^th^ locus, *w_im_* = 0 indicates two reference alleles, *λ_imr_* = 1 if these alleles are descended from group *r* (and 0 otherwise), and 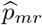 is the estimated alternate allele frequency on locus *m* within group *r*. Note that the estimator 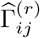 in expression (4) is a version of the VanRaden (2008) GRM in equation (1), modified so that genotypes only contribute to the relatedness estimate if they share the same local ancestry (*i.e*., when *λ_imr_* = *λ_jmr_* = 1). Like this, admixture is explicitly incorporated into the relatedness estimator. This modified GRM is scaled by the estimated genome-overlap 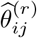 so that the impact on the group-specific additive genetic variance from a pair of individuals only comes from the proportion of their genes that originate from the same group. To model group-specific *V*_A_ with MAGBLUP-RI one can thus include the random effects 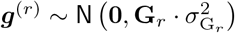 for each group *r*, where the entries of **G**_*r*_ are estimated in equation (4).

### An extension to wild systems

In order to make the genomic genetic group model applicable to wild systems, we need to extend MAGBLUP-RI such that it allows for heterozygosity, as well as for an arbitrary number of genetic groups *R* ≥ 2. These extension will allow us to model admixture in most wild study systems.

To allow for heterozygosity, we consider the contributions (allele effects) of the two alleles at each locus *separately*. We split the genotype indicator *w_im_* into two allele indicators 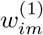 and 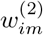, where the local ancestries are indicated by the binary variables 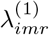 and 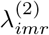, respectively (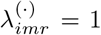 indicates local ancestry from group *r*). When we write 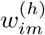, *h* ∈ {1, 2} thus denotes which of the two copies of the chromosomes the allele is located on (we only consider diploid organisms). We further assume the contributions to the total genetic value *U_i_* from a given locus to be equally weighted between its two present alleles. In other words, we assume a lack of dominance effects within loci with respect to their contributions to the phenotype, as the contribution from a heterozygous locus will be the mean of the effects of the possible homozygotes at the locus. As in MAGBLUP-RI, the allele effects are still assumed to depend on the locus, on the allele variant (reference or alternate), and on the local ancestry of the allele (*i.e*., the effect of the allele depends on which group the allele was inherited from). We also retain the assumption that allele variant indicators on different loci (*e.g*., *m* and *m*’ ≠ m) are uncorrelated (which results in a lack of LD). While the two allele indicators on the same locus are given identical probability distributions, we also assume that they are uncorrelated.

Similarly to Rio *et al*. (2020b) we consider the expected value and covariance between total genetic values *U_i_* of different individuals, but under an updated definition of *U_i_* where we model single allele effects and an arbitrary number of groups. We only present the results and assume segregation variances can be neglected, but the full model derivation (including segregation variances) is shown in Supporting Information S1. Our derivation shows that the statistical properties of *U_i_* found by Rio *et al*. (2020b) are retained under the new definition of *U_i_*. In other words, we show that equations (2) and (3) still hold in the extended model, if we sum from *r* = 1 up to *R* rather than 2 in both equations. Thus, our estimate of the GRM entries remains 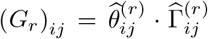. However, the parameters 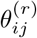 and 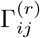 have somewhat expanded meanings in the extended model, as they now refer to overlap of *allele* ancestry and relatedness from comparing alleles *separately* (rather than genotypes), respectively. We therefore update their respective estimators to fit our single-allele paradigm.

Let us first generalize the parameter estimators from Rio *et al*. (2020b) to account for single alleles. Let the global ancestry proportion *π_ir_* for individual *i* in group *r* be estimated by the observed group membership proportion

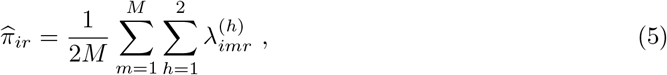

and the group allele-frequency *p_mr_* at locus *m* in group *r* by the observed group allele-frequency

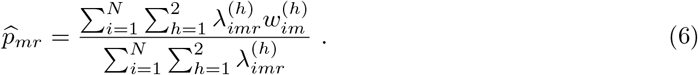

Since the base population induced by using GRMs corresponds to the population from which the allele frequency is derived (Hayes *et al*., 2009; Legarra, 2016), the base population of a genetic group will, due to equation (6), consist of its purebred individuals along with the parts of the genomes of admixed individuals which have local ancestry from that group. Admixed individuals are thus partially members of multiple base populations.

When dealing with local ancestry indicators 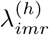 and allele indicators 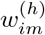 in different individuals, the assignment of a chromosome copy to be *h* = 1 or *h* = 2 is arbitrary and will not correspond between different individuals. These designations thus cannot be differentiated when comparing different individuals, so both alleles at locus *m* in one individual must be compared to both alleles on locus *m* in another individual. Thus, we estimate the genome overlap coefficient 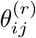 by

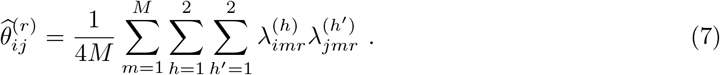

To arrive at the estimate for the group-conditional relatedness 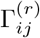, we again compare both alleles at locus *m* in individual *i* with both alleles at locus m in individual *j*:

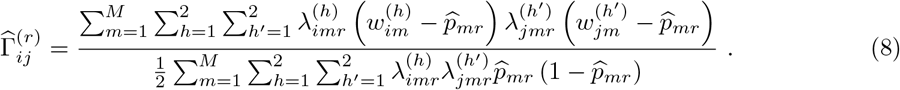

Our GRM with entries 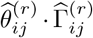 is a generalization of the well-known GRM **G** in equation (1), and also incorporates the use of local ancestry from the estimator in equation (4). It is easy to see that in the one-group case (*R* = 1) the GRM entries 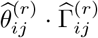 simplify to the entries of **G** (shown in Supporting Information S1).

### Application to a house sparrow metapopulation

We applied the extended genomic genetic groups animal model to a metapopulation of house sparrows living on islands in the Helgeland region in Northern Norway. This metapopulation has been subject to a long-term study since 1993, and the data used here include in total 4625 phenotypic records of wing length, body mass and tarsus length from 1932 sparrows measured between 1993 and 2016 on eight islands that are known to have inter-island dispersal (Baalsrud *et al*., 2014; Muff *et al*., 2019; Saatoglu *et al*., 2021). Large-scale genotyping of house sparrows from the study system has resulted in high-quality genotypes at 181 354 SNPs in 3032 individuals (including the 1932 phenotyped individuals, Niskanen *et al*., 2020). Across the 3032 × 181 354 genotypes in the genomic data set, roughly one third are heterozygous, which justifies the need for our extension of the genomic genetic groups model.

Subpopulations of sparrows located on the eight different islands vary in environmental conditions such as habitat, buffering against bad weather, and population density, and can be broadly defined as belonging to either the set of inner or outer islands (Baalsrud *et al*., 2014; Muff *et al*., 2019; Niskanen *et al*., 2020; Saatoglu *et al*., 2021). To account for possible genetic differences between the subpopulations originating from these two sets of islands, we partitioned the study population into genetic groups, namely an `inner` genetic group (encoded as 1), and an `outer` genetic group (encoded as 2). Sparrows from other islands in the study system (that were not systematically SNP-genotyped) are also present in the data set due to dispersal, and we place these sparrows into a third genetic group `other` (encoded as 3).

Throughout, we will be comparing the results from the genomic genetic group animal model to results from an otherwise equivalent pedigree-based model (Muff *et al*., 2019). The two models are identical, except that the pedigree-based model uses pedigree-derived global ancestries denoted *q_ir_* (see Wolak & Reid, 2017), rather than 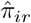, and pedigree-derived group-specific genetic relatedness matrices (see Muff *et al*., 2019) instead of the group-specific GRMs **G**_*r*_.

In a natural study system like ours, every individual is probably admixed at least to some extent due to dispersal in the past. To perform the local ancestry inference, however, we had to assign some individuals as purebred in the genetic groups of interest. We did this by classifying any individual that has both parents missing in the pedigree as a purebred in one of the groups for the genomic model, as determined by information about their natal island (Saatoglu *et al*., 2021, Table 1 shows the respective sizes of the purebred and admixed subpopulations). Thus, the starting condition of the genomic genetic groups model is as similar as possible to the corresponding pedigree-based genetic groups model (Muff *et al*., 2019) and hence allows a valid comparison between pedigree-based and genomic genetic group models. Conversely, individuals with at least one known parent in the pedigree were considered admixed in the genomic model. A further description of the empirical data from the house sparrow study system can be found in Supporting Information S3.

**Table 1:** The numbers of genotyped and phenotyped individuals in the purebred reference populations (Inner, Outer, Other), the admixed population (Admixed), and in total (Total). Note that all phenotyped individuals were genotyped.

|  | Inner | Outer | Other | Admixed | Total |
| --- | --- | --- | --- | --- | --- |
| Genotyped | 224 | 97 | 85 | 2626 | 3032 |
| Phenotyped | 197 | 73 | 41 | 1620 | 1932 |

As a prerequisite for our chosen local ancestry inference method (Dias-Alves *et al*., 2018), we performed gametic phasing of the SNP data, which involves identifying which of the two alleles at each locus was inherited from which parent. In terms of our mathematical notation, this step determines which alleles within an individual should be designated which values of *h* ∈ {1, 2} together. After using `PLINK 1.9` (Chang *et al*., 2015) to convert the genomic data to the appropriate input format, we used `Beagle 5.1` (Browning *et al*., 2018) with default settings to perform the gametic phasing. The phasing was done separately on each of the purebred populations and the admixed population since they are assumed to be genetically distinct. In addition to performing the gametic phasing, `Beagle` imputed any missing genotypes in the genomic data. The local ancestry 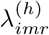 of every considered allele in the admixed population was then inferred via the command-line version of the Python package `Loter` (Dias-Alves *et al*., 2018).

With values for the allele variant indicators 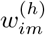 and local ancestry indicators 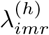 we estimated global ancestries **π**_r_, group-specific allele frequencies 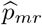 and group-specific GRMs **G**_*r*_ for *r* ∈ {1, 2, 3} using equations (5), (6), (7) and (8). Despite only 1932 sparrows having phenotype data, all 3032 genotyped sparrows were used in setting up the group-specific relatedness matrices **G**_*r*_ to improve the accuracy of the relatedness estimates. We used the R package `BGData` (Grueneberg & de los Campos, 2019) to manage the large genomic data sets, and to calculate the group-specific GRMs. Finally, a value of 10^-12^ was added to the diagonals of the **G**_*r*_ matrices to ensure positive-definiteness (since purebreds induce zeros on the diagonals).

Given the global ancestries **π**_*r*_ and group-specific GRMs **G**_*r*_, we can formulate the full genomic genetic groups model

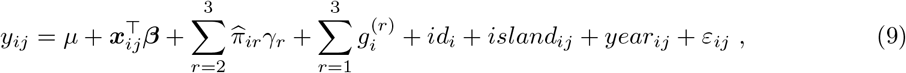

where *y_ij_* is the *j*th phenotypic measurement for individual *i*, *μ* is an intercept, ***x**_ij_* is a vector storing the fixed covariates sex (0 male, 1 female), age (in years since birth), month (May through August as a continuous covariate) and inbreeding coefficient *F*_GRM_ (computed by Niskanen *et al*., 2020), ***β*** is a vector of the fixed effects, 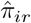 is *i*’s estimated global ancestry proportion in group *r*, and *γ_r_* is the fixed genetic group effects in groups *r* ∈ {1, 2, 3}. We set *γ*_1_ =0 for identifiability reasons, thus the genetic group effects *γ*_2_ and *γ*_3_ denote the deviations in the respective group’s mean total additive genetic effect from `inner` (Wolak & Reid, 2017). The random effects include group-specific genetic values 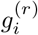, which are entries in the random vector 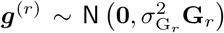, the permanent environmental effects 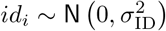, the effect of island of measurement 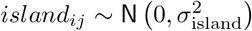, the effect of year of measurement 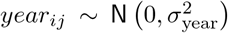, and the residual term 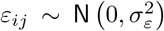. Inclusion of the island of measurement as a common environmental random effect is especially critical in this model, as it ensures the model can distinguish between environmental island effects and the genetic group effects. We implemented all genetic group animal models in a Bayesian framework with the `R-INLA` package (Rue *et al*., 2009, see tutorial in Supporting Information S2).

## Results

### Global ancestry proportions

As an initial plausibility check of the local ancestry inference, we first compared the estimated genomics-based global ancestry proportions 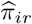 to their pedigree-derived counterparts *q_ir_*. Recall that *π_ir_* was not estimated for purebred individuals, but was instead assumed to be 0 or 1 for use in a reference panel. We therefore only compare 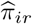 and *q_ir_* for admixed individuals that were also phenotyped. For most individuals in this subpopulation, the global ancestry proportions estimated with the two methods were relatively similar (Fig. 1), with correlations of 0.82, 0.83 and 0.60 for `inner`, `outer` and `other`, respectively. Both the correlations and the scatter plots thus indicate that the methods are in relatively good agreement.

**Figure 1:**
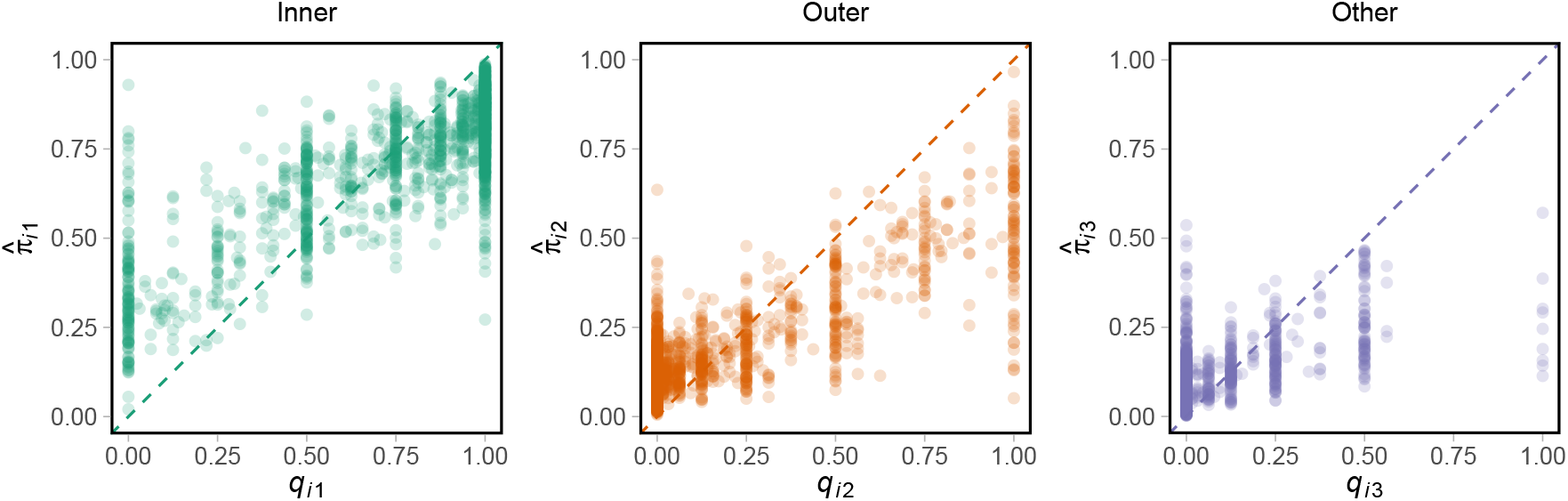
Scatter plots for global ancestry proportion derived from the pedigree (*x*-axes, *q_ij_*) and genomic local ancestry inference (*y*-axes, 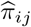). Each point refers to one individual. The plots only contain points for phenotyped admixed individuals (*N* = 1620). Points are partially transparent to show density patterns in areas with overlapping points. Diagonals are shown as dashed lines.

We observe that individuals with global ancestries equal to 0 or 1, or fractions like 0.5, 0.25 and 0.75, are common in the pedigree-based model, but not in the genomic model. These individuals are the offspring of two purebreds of the same group (and thus also considered purebred by the pedigree model), “hybrid” offspring of purebreds from different groups, and offspring of hybrids back-crossed with purebred individuals, respectively. The accumulation of these values reflects that the pedigree-based model operates with *expected* inheritance of ancestry (which is, *e.g*., exactly 0.5 in a hybrid), whereas the genomic model considers *realized* ancestries. Another noteworthy difference is that more points lie above the diagonal than below for the `inner` global ancestry proportions (when *q*_*i*1_ is below ca. 0.75), reflecting that the genomic method tends to assign larger `inner`-ancestries than the pedigree-based method, while the opposite is true for `outer` (more points lie under the diagonal, for *q*_*i*2_ > 0.25). Both methods rarely assign large (> 0.5) admixture proportions to the `other` group, but the pedigree-based method considers more individuals purebred in this group. Moreover, the agreement between pedigree-derived and genomics-derived global ancestries is lowest for the `other` group, which is not surprising given that the group is a heterogeneous collection of individuals from all remaining islands. Generally, the methods seem to disagree the most for individuals that are considered purebred by just the pedigree (*q_ir_* ∈ {0, 1}), and mostly agree regarding individuals with intermediate global ancestry proportions.

### Comparison of pedigree-based and genomic results

We report posterior means and 95% highest posterior density credible intervals (HPD CIs) for all parameters in the models, and also posterior modes for the random effect variances since they potentially have skewed posterior distributions (Tables 2 and 3). Additionally, the full posterior distributions for the genetic group means *γ_r_* and variances 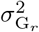 are displayed graphically for each of the three groups *r* ∈ {1, 2, 3} (Figs. 2 and 3, respectively). Since `inner` serves as the baseline for the mean genetic value, *γ*_1_ was fixed to zero and therefore has no posterior distribution.

**Figure 2:**
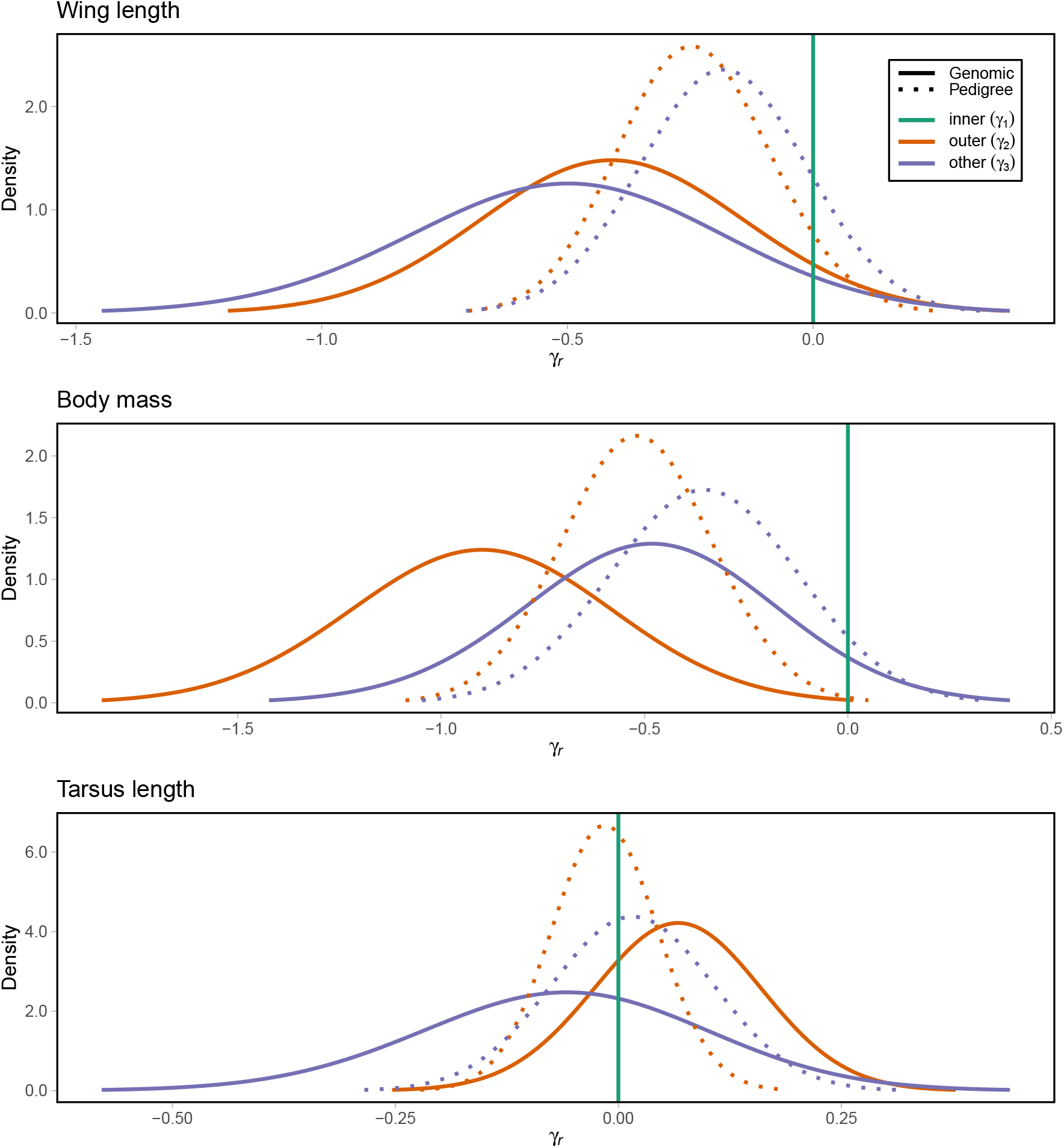
The estimated posterior distribution of genetic group effects *γ_r_* (*i.e*., mean genetic value) in the models for wing length (top), body mass (middle) and tarsus length (bottom). Posterior effects for the different genetic groups are shown in different colors. Posteriors from genomic models have solid lines, whereas the pedigree-based model posteriors are shown with dotted lines. Since `inner` is assumed to be the baseline mean, *γ*_1_ = 0 is shown as a vertical line.

**Figure 3:**
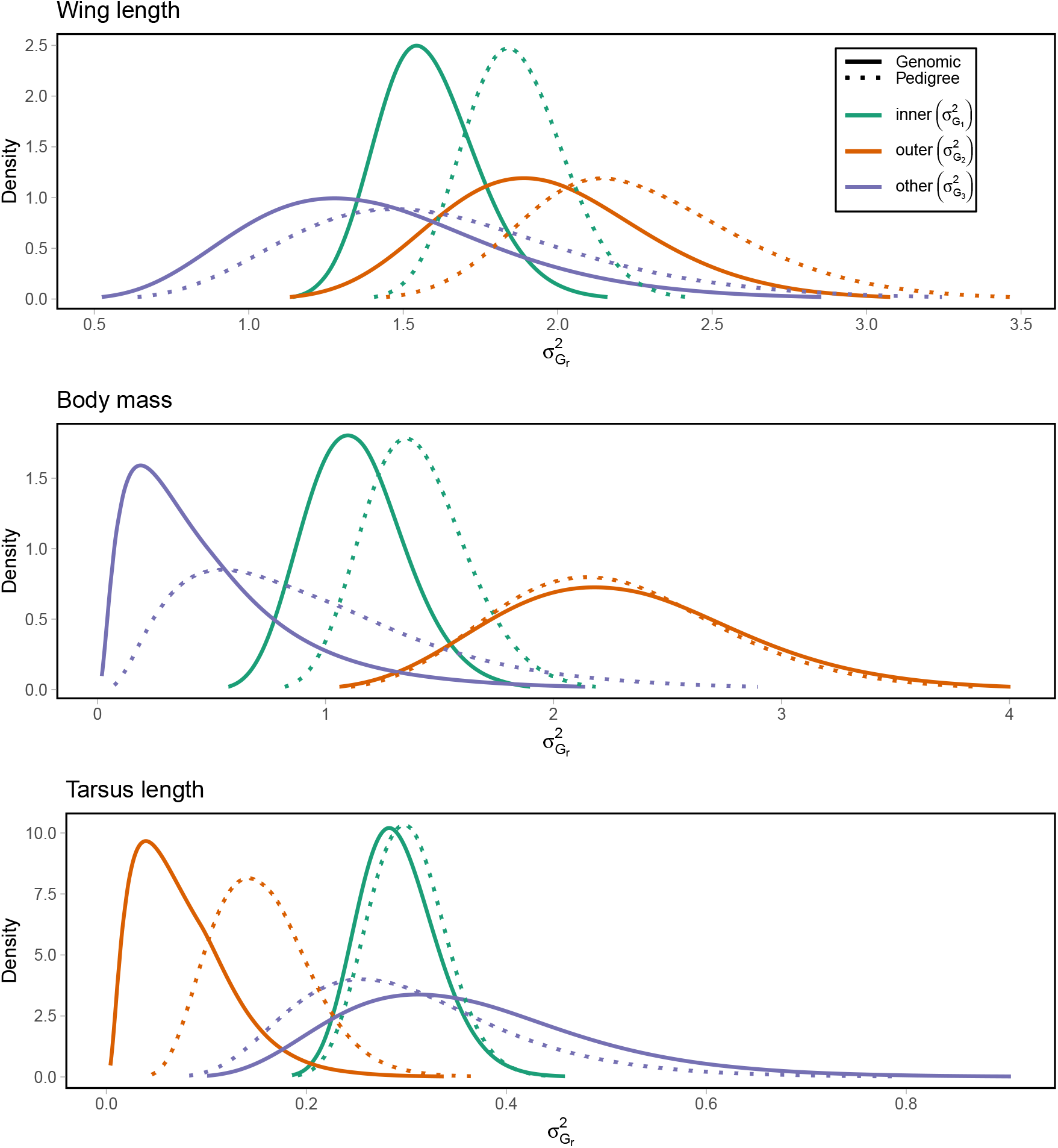
The estimated posterior distribution of group-specific additive genetic variances 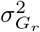 in the models for wing length (top), body mass (middle) and tarsus length (bottom). Posterior variances for the different genetic groups are shown in different colors. Posteriors from genomic models have solid lines, whereas the pedigree-based model posteriors are shown with dotted lines.

**Table 2:** Posterior statistics for the fixed effects of the genetic group animal models, as derived from pedigree-based and genomic genetic group models for the three investigated phenotypic traits. For each parameter, the posterior mean is reported in the first row, and the 95% HPD CI in the second row. The parameters denote the effects of being female compared to male (*β*_sex_), inbreeding (*β*_*F*_GRM__), month (*β*_month_) and age (*β*_age_). The genetic group effects *γ*_2_ and *γ*_3_ of `outer` and `other`, respectively, denote the effect of being purebred in these groups relative to `inner`.

| Parameter | Wing length |  | Body mass |  | Tarsus length |  |
| --- | --- | --- | --- | --- | --- | --- |
|  | Pedigree | Genomic | Pedigree | Genomic | Pedigree | Genomic |
| $\beta_{\text{sex}}$ | -2.75<br>(-2.89, -2.62) | -2.76<br>(-2.89, -2.63) | 0.47<br>(0.29, 0.64) | 0.47<br>(0.29, 0.65) | -0.09<br>(-0.16, -0.02) | -0.08<br>(-0.15, -0.01) |
| $\beta_{F_{\text{GRM}}}$ | -2.14<br>(-3.68, -0.60) | -1.77<br>(-3.32, -0.23) | -0.98<br>(-3.06, 1.09) | -0.93<br>(-3.02, 1.15) | -0.65<br>(-1.46, 0.16) | -0.67<br>(-1.48, 0.15) |
| $\beta_{\text{month}}$ | -0.19<br>(-0.23, -0.15) | -0.19<br>(-0.23, -0.15) | -0.30<br>(-0.36, -0.24) | -0.30<br>(-0.36, -0.24) | 0.03<br>(0.02, 0.03) | 0.03<br>(0.02, 0.04) |
| $\beta_{\text{age}}$ | 0.46<br>(0.43, 0.50) | 0.46<br>(0.43, 0.50) | 0.09<br>(0.03, 0.15) | 0.09<br>(0.03, 0.15) | -0.00<br>(-0.01, 0.00) | -0.00<br>(-0.01, 0.00) |
| $\gamma_2$ | -0.24<br>(-0.53, 0.07) | -0.41<br>(-0.93, 0.13) | -0.52<br>(-0.88, -0.15) | -0.90<br>(-1.54, -0.27) | -0.02<br>(-0.13, 0.10) | 0.07<br>(-0.12, 0.25) |
| $\gamma_3$ | -0.18<br>(-0.51, 0.15) | -0.51<br>(-1.14, 0.12) | -0.35<br>(-0.81, 0.10) | -0.49<br>(-1.12, 0.12) | 0.02<br>(-0.16, 0.19) | -0.06<br>(-0.39, 0.25) |

**Table 3:** Posterior statistics for the random effect variances of the genetic group animal models, as derived from pedigree-based and genomic genetic group models for the three investigated phenotypic traits. For each parameter, the posterior mode and mean (formatted mode;mean) are reported in the first row, and the 95% HPD CI in the second row. The parameters denote variance explained by year of measurement 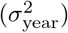, island of measurement 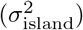, permanent environmental effects 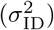, group-specific *V*_A_ (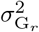, *r* = 1 for `inner`, *r* = 2 for `outer`, *r* = 3 for `other`) and residual environmental effects 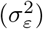.

| Parameter | Wing length |  | Body mass |  | Tarsus length |  |
| --- | --- | --- | --- | --- | --- | --- |
|  | Pedigree | Genomic | Pedigree | Genomic | Pedigree | Genomic |
| $\sigma_{\text{year}}^2$ | 0.03;0.04<br>(0.01, 0.09) | 0.05;0.06<br>(0.02, 0.14) | 0.05;0.06<br>(0.01, 0.16) | 0.04;0.05<br>(0.01, 0.14) | 0.01;0.01<br>(0.00, 0.04) | 0.01;0.01<br>(0.00, 0.03) |
| $\sigma_{\text{island}}^2$ | 0.09;0.12<br>(0.02, 0.35) | 0.07;0.08<br>(0.02, 0.21) | 0.11;0.14<br>(0.03, 0.45) | 0.10;0.14<br>(0.02, 0.47) | 0.00;0.01<br>(0.00, 0.03) | 0.00;0.01<br>(0.00, 0.02) |
| $\sigma_{\text{ID}}^2$ | 0.34;0.35<br>(0.21, 0.52) | 0.43;0.44<br>(0.30, 0.61) | 1.03;1.05<br>(0.75, 1.43) | 1.09;1.11<br>(0.83, 1.48) | 0.35;0.36<br>(0.31, 0.41) | 0.35;0.35<br>(0.31, 0.40) |
| $\sigma_{G_1}^2$ | 1.85;1.86<br>(1.57, 2.20) | 1.57;1.58<br>(1.29, 1.93) | 1.38;1.40<br>(1.00, 1.90) | 1.12;1.14<br>(0.75, 1.61) | 0.30;0.30<br>(0.23, 0.38) | 0.29;0.29<br>(0.22, 0.38) |
| $\sigma_{G_2}^2$ | 2.23;2.27<br>(1.68, 3.08) | 1.94;1.97<br>(1.37, 2.71) | 2.23;2.28<br>(1.41, 3.41) | 2.27;2.32<br>(1.37, 3.55) | 0.15;0.16<br>(0.08, 0.27) | 0.07;0.08<br>(0.01, 0.20) |
| $\sigma_{G_3}^2$ | 1.59;1.65<br>(0.89, 2.79) | 1.38;1.43<br>(0.75, 2.42) | 0.83;0.96<br>(0.20, 2.45) | 0.40;0.53<br>(0.06, 1.71) | 0.29;0.31<br>(0.14, 0.57) | 0.35;0.36<br>(0.17, 0.67) |
| $\sigma_{\varepsilon}^2$ | 0.98;0.98<br>(0.93, 1.04) | 0.98;0.98<br>(0.93, 1.03) | 2.88;2.89<br>(2.73, 3.05) | 2.90;2.90<br>(2.74, 3.06) | 0.02;0.02<br>(0.02, 0.02) | 0.02;0.02<br>(0.02, 0.02) |

Generally, the agreement between the pedigree-based and the genomic models regarding fixed effects is quite good. The results indicate that sparrows descended from the `inner` islands tend to have genes for longer wings and body mass, as visible by the group-differences in the genetic group means *γ_r_* (Table 2 and Fig. 2). These estimated genetic group effects are somewhat more pronounced in the genomic models than the pedigree-based models, but the model disagreements are small compared to the uncertainties in the estimates. For tarsus length, the impact of genetic group ancestry on the mean genetic value is small to non-existent. Interestingly, this implies that genetically, sparrows with `outer` ancestry have longer tarsi relative to their body size (*i.e*., wing length and body mass) compared to sparrows with `inner` ancestry. Among the remaining fixed effects parameters, we see that for wing length the pedigree-based model finds slightly stronger negative effects of inbreeding than the genomic model, but again the differences are small relative to the uncertainty (Table 2). The pedigree-based and genomic models are in close agreement regarding all remaining fixed effects.

For the group-specific additive genetic variances the two model paradigms produce posteriors with very similar shapes, and with similar posterior modes (Table 3 and Fig. 3). Either the posteriors overlap almost completely, or the genomic model gives a posterior shifted to lower values than the pedigree-based model – the latter is a common and expected pattern when comparing these types of models (discussed further in the next section). In all models we find among-group differences in *V*_A_ for each phenotypic trait. For wing length and body mass, the *V*_A_ in `outer` tends to be higher than in `inner`, which again tends to be higher than in `other` (Fig. 3). When it comes to tarsus length, the `outer` group probably has a smaller *V*_A_ than `inner` and `other`. Posteriors for the *V*_A_ are usually narrowest in `inner`, which is expected because sample sizes are largest in this group. All these differences between the groups (including differences in the *γ_r_* estimates) indicate that the use of genetic group models is justified, or even necessary.

## Discussion

In this article we have extended the genetic group animal model such that it can be used with genomic data to estimate group-specific *V*_A_ for wild admixed populations. In order to derive the group-specific GRMs, we incorporated local ancestry information and considered single alleles to allow heterozygous genotypes. This is a generalization of previous methodology that was developed within a plant breeding setup, where it was assumed that only homozygote genotypes were present (Rio *et al*., 2020b). Furthermore, we allow for any number of genetic groups, rather than only two. As a proof of concept, we applied the extended method to a metapopulation of house sparrows, and show that the results are in line with the results from a corresponding, but pedigree-based, genetic groups model.

In our illustrative example there was relatively good agreement between the genomic model and the pedigree-based model results, though we often observe stronger genetic group effects and smaller group-specific *V*_A_ in the genomic model. The disagreement on genetic group effects *γ_r_* might be explained in part by the discrepancies in global ancestry proportions (Fig. 1). Notably, the genomic vs. pedigree differences in *V*_A_ (*i.e*., smaller *V*_A_ in the genomic model) correspond to differences observed in otherwise equivalent genetic group animal models with homogeneous additive genetic variances (Fig. S1, Supporting Information S4.1). This result suggests that the genomic genetic group method itself is not introducing a downward bias in *V*_A_, but rather the inherent differences in using pedigree-based vs. genomic methods (see *e.g*., Powell *et al*., 2010). The pattern of *V*_A_ being smaller when estimated from genomic data than when estimated from pedigree data is often observed and has been extensively discussed in the literature (Legarra, 2016; Yang *et al*., 2017; Evans *et al*., 2018; Gervais *et al*., 2019).

Our results regarding the genetic group structure of the house sparrow system otherwise mirror what has been found previously regarding wing length and body mass (Muff *et al*., 2019), namely group-differences in mean genetic value and *V*_A_, which respectively suggest `outer`-descended birds have shorter wings and lower body mass, and lower *V*_A_ in these phenotypes compared to `inner`. Additionally, we find a lower *V*_A_ in tarsus length among `inner`-descended birds. Note that we have no guarantee against further genetic substructures within the genetic groups (*e.g*., `inner` is made up of five different islands), which could bias the results for group-specific *V*_A_ (Wolak & Reid, 2017). To sum up, our genetic group model results indicate that `outer` and `inner` island sparrows may be genetically divergent in some of their phenotypes, possibly caused by local adaption to different habitats, and that the adaptive potential (*i.e*., *V*_A_) of these possibly locally adapted phenotypic traits could depend on their group ancestry.

Genetic group models are not only attractive because they eliminate the source of bias in estimation of *V*_A_ caused by genetic substructures in the base population, but also because they help identify differences in *V*_A_ for admixed and/or genetically differentiated populations. Since higher *V*_A_ implies higher potential for rapid evolutionary change due to selection, such as adaptation to any environmental changes, the identification of differences is important especially in the light of the current rapid change of environments due to anthropogenic effects (Wood & Brodie III, 2016; Gienapp *et al*., 2017). However, the estimation of group-specific *V*_A_ obviously increases computational requirements and imposes higher demands on the data. The model is dramatically simplified if the assumption of differences in the group-specific 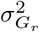 for *r* = 1,…, *R* is dropped, while still allowing for differences in the mean genetic (i.e. breeding) values *γ_r_*. Thus, only a single, homogeneous *V*_A_ without admixture-induced bias is estimated (Wolak & Reid, 2017). Such a simplified model will only need to estimate the group-specific global ancestry proportions *π_ir_* for each individual *i*, as given in equation (5). In this simplified case, however, one does not need to make the computationally heavy detour via the inference of local ancestries of each allele, as we did here, but could directly estimate *π_ir_* using faster methods (such as Raj *et al*., 2014).

A main difference in the methodology discussed here compared to Rio *et al*. (2020b) is that we assume segregation variances to be negligible. As previously mentioned, segregation variances are expected to be negligible under the infinitesimal model, which is commonly assumed to hold for most complex traits, in particular in natural populations (Hill & Kirkpatrick, 2010; Hill, 2012). The inclusion of segregation variances would therefore unnecessarily increase the demands on computational power and data quality, and potentially prohibit the use of the model. To illustrate that omitting segregation terms is indeed not critically affecting the parameter estimates, we also fitted modified models which do include segregation variances to confirm that segregation variances are indeed negligible (Fig. S2, Tables S1-S2, Supporting Information S4.2).

Another interesting assumption of the genomic model is that it allows allele effects to depend on the local ancestry of the allele, not merely the variant. Such group-specific allele effects can, for example, result from the groups having different strengths of LD between SNPs and quantitative trait loci (QTL, Rio *et al*., 2020a). Among-group differences in strength and extent of LD can be caused by differences in effective population sizes leading to different levels of genetic drift, and by differences in local selection pressures on the trait in question. While we do not explicitly model LD between loci, we thus implicitly account for differences in LD across groups through the group-specific allele effects. Hagen *et al*. (2020) indeed found that levels of LD in house sparrows were generally higher and remained higher over longer distances along the chromosomes on islands in our study system with smaller effective population sizes (generally `outer` islands, which recently went through a strong population bottleneck, see Baalsrud *et al*., 2014). Group-specific allele effects can also be caused by QTL having epistatic interactions with loci whose allele frequencies differ between groups (Rio *et al*., 2020a). Loci with group-differences in allele frequencies are common in our study system (Fig. S3, Supporting Information S4.3), as there is evidence for genetic differentiation between the islands (Niskanen *et al*., 2020; Saatoglu *et al*., 2021). Allowing for group-specific allele effects may thus be justified, but further investigations would be required to confirm their presence in the study system.

An important prerequisite when using genetic group models is that there is at least some level of admixture between genetic groups. If the groups were isolated without any admixture, we might otherwise not be able to disentangle evolutionary processes from environmental effects (Wood & Brodie III, 2016; Hoffmann *et al*., 2017). Conversely, having purebred individuals present in the data set is not a requirement in general. In fact, in a wild system with dispersal such as the one we consider, it is more realistic to assume all individuals as admixed, if we look far enough back in time. However, the decision to include purebreds must sometimes be made out of necessity. Here, for example, our choice to consider individuals with both parents missing in the pedigree as purebred increased the comparability of our results to those derived from the pedigree-based genetic groups animal model. Additionally, the purebreds were needed to be used as reference panels in the local ancestry inference.

In a more general setup, it might not always be obvious how the “purebred” reference populations of the different genetic groups should be defined or selected, in particular when pedigree data is missing. Since individuals labeled as “purebreds” will usually not actually be purebred in a strict sense, but are rather individuals with a mix of ancestries from various populations, the purebred (or reference) population can in principle be defined in any desired way that is relevant for the study at hand. As an example, sub-populations present (or rather, individuals captured) in certain areas the first year of a multi-year study can be used as reference panels in local ancestry estimation, which resembles what we did in the sparrow example. This approach requires previous knowledge about the system – for example, we knew from previous studies that the different island-group populations are somewhat genetically differentiated (Muff *et al*., 2019; Niskanen *et al*., 2020; Saatoglu *et al*., 2021). Alternatively, population structure estimation tools can detect the identity of genetic groups, global ancestry proportions and/or the number of groups (*e.g*., Raj *et al*., 2014; Kuismin *et al*., 2017), or even local ancestries directly (*e.g*., Utsunomiya *et al*., 2020). These methods do not generally rely on reference panels. When using a tool where only global ancestry proportions in a given group are detected, individuals with global ancestry higher than some threshold (*e.g*., 0.99) can be assigned as purebred in that group and serve as reference panels to find the local ancestries of the remaining (admixed) individuals (Geza *et al*., 2019; Schubert *et al*., 2020). In short, use of the genomic genetic group model requires either previous knowledge about the population structure, or investigating it with additional tools.

In summary, we believe that the genomic genetic groups animal model is a useful new tool that allows researchers to estimate and compare group-specific additive genetic values and variances in wild admixed populations. Thanks to the generalization from pedigree-based to genomic animal models, researchers are no longer restricted to using long-term data collected over many generations.

To generate the data necessary to fit a genetic group animal model, it is now sufficient to sample phenotype and genotype information from individuals in an admixed natural population at a given time. In a conservation management perspective, the genomic genetic group approach thus opens up possibilities for researchers to estimate *V*_A_ within different populations to find the ones with greatest need of management to ensure sufficient adaptive potential, and also to examine how *V*_A_ in a population is affected by admixture (*e.g*., immigration), to actually quantify how management actions and/or natural dispersal will potentially aid in evolutionary rescue (Whiteley *et al*., 2015).

## Acknowledgments

We thank the many researchers, students, and fieldworkers who helped in collecting the empirical data on house sparrows, and laboratory technicians for assistance with laboratory analyses. This study was supported by grants from the Norwegian Research Council (projects 274930 and 302619) and its Centre of Excellence funding scheme (project 223257). The research done on house sparrows was carried out in accordance with permits from the Norwegian Food Safety Authority and the Bird Ringing Centre at Stavanger Museum, Norway. Genotyping on the custom house sparrow Affymetrix Axiom 200K SNP array was carried out at CIGENE, Norwegian University of Life Sciences, Norway. The computations were performed on resources provided by the NTNU IDUN/EPIC computing cluster (Själander *et al*., 2019).

## Conflict of Interest statement

The authors have no conflicts of interest to declare.

## Authors’ contributions

SM, KA and HJ conceived the idea for the study. KA developed and implemented the model, analyzed the data and wrote the manuscript. SM helped develop and implement the model. HJ provided the sparrow data. SM and HJ provided methodological support, and participated in writing and editing the manuscript. All authors read and approved the final manuscript.

## Data availability

For the pedigree, phenotype and genotype data use the Dryad repository from Niskanen *et al*. (2020): https://doi.org/10.5061/dryad.m0cfxpp10.

For the natal island information use the Dryad repository from Saatoglu *et al*. (2021): https://doi.org/10.5061/dryad.xgxd254h1.

## Notes

### Competing Interest Statement

The authors have declared no competing interest.

