## Supporting Information for "Genomic estimation of quantitative genetic parameters in wild admixed populations"

### S1: Mathematical derivations

This supplement contains the mathematical justification for the genomic genetic group animal model, as well as details of the derivations.

#### Notation and definitions

In this section we define all mathematical notation and definitions used in the derivations. Refer back to this section if necessary.

##### Symbols

Indices for genetic groups: $r$, $r'$, $r''$, $r^*$

Number of groups: $R$

Set containing numbers $1, \ldots, R$: $\mathcal{R}$

Indices for individuals: $i, j$

Number of individuals: $N$

Indices for loci : $m, m'$

Number of loci: $M$

Indices for chromosome copy: $h, h'$

##### Allele effects $\beta_{mr}^{\mathbf{ref}}$ and $\beta_{mr}^{\mathbf{alt}}$

The allele effects are defined as the numerical contributions to the phenotype from a homozygous locus $m$ originating from group $r$, with two reference ($\beta_{mr}^{\text{ref}}$) or alternate ($\beta_{mr}^{\text{alt}}$) alleles.

##### Global ancestry $\pi_{ir}$

The global ancestry $\pi_{ir}$ is defined as the proportion of individual $i$'s genome that is descended from genetic group $r$. Note that $\sum_{r=1}^{R} \pi_{ir} = 1$.

##### Group-specific allele frequency $p_{mr}$

The group-specific allele frequency $p_{mr}$ is the allele frequency of the alternate allele at locus $m$, given that the allele is descended from group $r$.

##### Local ancestry indicator $\Lambda_{imr}^{(h)}$

The local ancestry indicator $\Lambda_{imr}^{(h)}$ is a random variable indicating whether the allele in individual $i$, on locus $m$, chromosome copy $h$ is descended from group $r$. Thus, $\Lambda_{imr}^{(h)} = 1$ if this allele is descended from group $r$, and $\Lambda_{imr}^{(h)} = 0$ otherwise. The random vector of local ancestry indicators $\boldsymbol{\Lambda}_{im}^{(h)} = \left[\Lambda_{im1}^{(h)}, \Lambda_{im2}^{(h)}, \ldots, \Lambda_{imR}^{(h)}\right]$ has a categorical distribution, that is, a multinomial distribution with only one trial. In other words, an allele can only be descended from one group[1], and the probability of being descended from a group depends on the global ancestry of the individual. As a result, $\mathsf{E}\left(\Lambda_{imr}^{(h)}\right) = \mathsf{P}\left(\Lambda_{imr}^{(h)} = 1\right) = \pi_{ir}$ and $\sum_{r=1}^{R} \Lambda_{imr}^{(h)} = 1$.

##### Allele variant indicator $W_{imr}^{(h)}$

The allele variant indicator $W_{imr}^{(h)}$ is a random variable indicating whether the alternate allele is present in individual $i$'s locus $m$, chromosome copy $h$. Thus, $W_{im}^{(h)} = 1$ if the alternate allele is present, and $W_{im}^{(h)} = 0$ otherwise. The probability of this occurring will depend on the alternate allele frequency, which we have allowed to differ between genetic groups. Thus, the distribution of $W_{im}^{(h)}$ is given conditionally on the local ancestry of the allele:
$W_{im}^{(h)} \,\Big|\, \left(\Lambda_{imr}^{(h)} = 1\right) \sim \text{Bernoulli}(p_{mr})$.

##### Group-specific relatedness $\Gamma_{ij}^{(r)}$

The group-specific relatedness $\Gamma_{ij}^{(r)}$ denotes the within-group genetic similarity of two individuals $i \neq j$, that is, the relatedness between the two individuals when only considering alleles descended from the same genetic group. We define this as the conditional correlation
$\Gamma_{ij}^{(r)} = \mathsf{Corr}\left(W_{im}^{(h)},\ W_{jm}^{(h)} \,\Big|\, \Lambda_{imr}^{(h)} = 1,\ \Lambda_{jmr}^{(h)} = 1\right)$.

##### Ancestry overlap $\theta_{ij}^{(rr')}$

The ancestry overlap parameter $\theta_{ij}^{(rr')}$ denotes the proportion of alleles where $i$'s allele belongs to group $r$ and $j$'s allele belongs to group $r'$, which can be written mathematically as $\mathsf{E}\left(\Lambda_{imr}^{(h)}\ \Lambda_{jmr'}^{(h)}\right)$. When $r = r'$ we will write $\theta_{ij}^{(rr)} = \theta_{ij}^{(r)}$ for simplicity. Note that $\theta_{ij}^{(rr')} \neq \theta_{ij}^{(r'r)}$ in general.

[1] In assuming that an allele can only be descended from one of the reference populations, the mathematical model implicitly assumes that at some point in the past there existed reference populations which were completely isolated from each other. More precisely, we assume that in the full true pedigree of the species, none of the purebred individuals are descended from a purebred from a different reference population. However, as the reviewer points out, in practice it is artificial to assume that the reference populations are (or have ever been) completely isolated. As we say in the Discussion, it would have been more realistic to assume that the purebred individuals were also admixed, especially in a wild system with dispersal such as ours. Conversely, assuming the presence of segregated reference populations can be more realistic in breeding programs, since the populations can be deliberately created (as in the dataset used by Rio *et al.*, 2020).

**Covariance between local ancestries $\Delta_{ij}^{(rr')}$**

Defined as $\Delta_{ij}^{(rr')} = \mathsf{Cov}\left(\Lambda_{imr}^{(h)}, \Lambda_{jmr'}^{(h)}\right)$. By the using the properties of covariance we can re-write this definition as $\Delta_{ij}^{rr'} = \mathsf{Cov}(\Lambda_{imr}^{(h)}, \Lambda_{jmr'}^{(h)}) = \mathsf{E}(\Lambda_{imr}^{(h)}\Lambda_{jmr'}^{(h)}) - \mathsf{E}(\Lambda_{imr}^{(h)})\mathsf{E}(\Lambda_{jmr'}^{(h)}) = \theta_{ij}^{(rr')} - \pi_{ir}\pi_{jr'}$ . As we do for $\theta$, we will write $\Delta_{ij}^{(rr)} = \Delta_{ij}^{(r)}$ for simplicity.

**Mean-centered local ancestry indicator $\widetilde{\Lambda}_{imr}^{(h)}$**

We define a mean-centered local ancestry indicator $\widetilde{\Lambda}_{imr}^{(h)} = \Lambda_{imr}^{(h)} - \pi_{ir}$, so that $\mathsf{E}\left(\widetilde{\Lambda}_{imr}^{(h)}\right) = 0$.

**Mean-centered group-conditional allele variant indicator $\widetilde{W}_{imr}^{(h)}$**

Defined as $\widetilde{W}_{imr}^{(h)} = \Lambda_{imr}^{(h)}\left(W_{imr}^{(h)} - p_{mr}\right)$. Note that we can show that $\mathsf{E}\left(\widetilde{W}_{imr}^{(h)}\right) = 0$ by The Law of Total Expectation.

**Total genetic value $U_i$**

The total genetic value $U_i$ is the sum of the allele contributions from all loci to individual $i$'s phenotype. We model the contribution from a given locus $m$ to $U_i$ as the mean of the two allele effects on that locus. We assume a lack of dominance effects, so the contribution from heterozygous loci will thus be the mean of the two possible homozygous contributions. For example, when one of the alleles at locus $m$ is the reference allele descended from group $r$, and the other allele is the alternate allele descended from group $r'$, then locus $m$ contributes $\frac{1}{2}\left(\beta_{mr}^{\text{ref}} + \beta_{mr'}^{\text{alt}}\right)$ to the total genetic value $U_i$. Or, if the locus is homozygous with, say, two reference alleles descended from the same genetic group, then the contribution will be $\frac{1}{2}\left(\beta_{mr}^{\text{ref}} + \beta_{mr}^{\text{ref}}\right) = \beta_{mr}^{\text{ref}}$. Thus, we define the total genetic value as $U_i = \sum_{m=1}^{M}\sum_{h=1}^{2}\sum_{r=1}^{R}\frac{1}{2}\Lambda_{imr}^{(h)}\left[\beta_{mr}^{\text{ref}} + W_{im}^{(h)}\left(\beta_{mr}^{\text{alt}} - \beta_{mr}^{\text{ref}}\right)\right]$, where we take the mean on each locus as mentioned, and use the indicators $\Lambda_{imr}^{(h)}$ and $W_{im}^{(h)}$ to ensure the right contribution from each allele. By considering the mean of the total genetic value $U_i$ we can find genetic group mean $\gamma_r$ and by considering the covariance between the total genetic values of two individuals $U_i$ and $U_j$ we can find the covariance structure that we want our random effects (*i.e.*, the group-specific genetic values) to capture.

**Genetic group effect $\gamma_r$**

$\gamma_r$ denotes the expected total genetic value if all alleles in an individual belong to group $r$. Inserting $\Lambda_{imr}^{(h)} = 1$ for all $m, h$ into the definition of $U_i$, and taking the mean we see $\gamma_r =$

$\sum_{m=1}^{M}\sum_{h=1}^{2}\frac{1}{2}\left[\beta_{mr}^{\text{ref}}+p_{mr}\left(\beta_{mr}^{\text{alt}}-\beta_{mr}^{\text{ref}}\right)\right]$. This can also be written as $\gamma_r=\sum_{m=1}^{M}\gamma_{mr}$ where $\gamma_{mr}$ is the expected contribution from locus $m$ if its allele belong to group $r$.

##### Group-specific $V_{\mathbf{A}}$ $\sigma^2_{\mathbf{G}_r}$

We define group-specific additive genetic variance as $\sigma^2_{\text{G}_r}=\frac{1}{2}\sum_{m=1}^{M}p_{mr}\left(1-p_{mr}\right)\left(\beta_{mr}^{\text{alt}}-\beta_{mr}^{\text{ref}}\right)^2$.

##### Segregation variance $\sigma^2_{\mathbf{S}_{rr'}}$

We define the segregation variance as $\sigma^2_{\text{S}_{rr'}}=\frac{M}{2M-1}\sum_{m=1}^{M}\left(\gamma_{mr}-\gamma_{mr'}\right)^2-\frac{1}{2M-1}\left(\gamma_r-\gamma_{r'}\right)^2$. Note that $\sigma^2_{\text{S}_{rr}}=0$ and that $\sigma^2_{\text{S}_{rr'}}=\sigma^2_{\text{S}_{r'r}}$. Also note that when $M$ is very large, we can say $\sigma^2_{\text{S}_{rr'}}\approx\frac{1}{2}\sum_{m=1}^{M}\left(\gamma_{mr}-\gamma_{mr'}\right)^2$, which is the definition of segregation variance in the $F_2$ generation given by Lynch & Walsh (1998).

#### Assumptions

### LD

There is a lack of LD between alleles, both within and across groups and individuals, so
$\mathsf{Corr}\left(W_{im}^{(h)},W_{jm'}^{(h')}\,\middle|\,\Lambda_{imr}^{(h)}=1,\Lambda_{jm'r'}^{(h')}=1\right)=0$ for all $i,\ j,\ m,\ m'\neq m,\ r,\ r',\ h,\ h'$, and
$\mathsf{Corr}\left(W_{im}^{(h)},W_{jm}^{(h')}\,\middle|\,\Lambda_{imr}^{(h)}=1,\Lambda_{jmr'}^{(h')}=1\right)=0$ for all $i,\ j,\ m,\ r,\ r',\ h,\ h'\neq h$.

#### Mean genetic value

We calculate the mean genetic value of an individual $i$ as follows.

$$\begin{aligned}\mathsf{E}\left(U_i\right)&=\sum_{r=1}^{R}\sum_{m=1}^{M}\sum_{h=1}^{2}\frac{1}{2}\mathsf{E}\left\{\Lambda_{imr}^{(h)}\left[\beta_{mr}^{\text{ref}}+W_{im}^{(h)}\left(\beta_{mr}^{\text{alt}}-\beta_{mr}^{\text{ref}}\right)\right]\right\}\\&=\sum_{r=1}^{R}\sum_{m=1}^{M}\sum_{h=1}^{2}\frac{1}{2}\left[\pi_{ir}\beta_{mr}^{\text{ref}}+\mathsf{E}\left(W_{im}^{(h)}\Lambda_{imr}^{(h)}\right)\left(\beta_{mr}^{\text{alt}}-\beta_{mr}^{\text{ref}}\right)\right]\ .\end{aligned}$$

We can show that $\mathsf{E}\left(W_{im}^{(h)}\Lambda_{imr}^{(h)}\right)=p_{mr}\pi_{ir}$, and so

$$\begin{aligned}\mathsf{E}\left(U_i\right)&=\sum_{r=1}^{R}\sum_{m=1}^{M}\sum_{h=1}^{2}\frac{1}{2}\left[\pi_{ir}\beta_{mr}^{\text{ref}}+p_{mr}\pi_{ir}\left(\beta_{mr}^{\text{alt}}-\beta_{mr}^{\text{ref}}\right)\right]\\&=\sum_{r=1}^{R}\sum_{m=1}^{M}\sum_{h=1}^{2}\frac{1}{2}\left[\pi_{ir}\left(\beta_{mr}^{\text{ref}}+p_{mr}\left(\beta_{mr}^{\text{alt}}-\beta_{mr}^{\text{ref}}\right)\right)\right]\\&=\sum_{r=1}^{R}\sum_{m=1}^{M}\sum_{h=1}^{2}\frac{1}{2}\pi_{ir}\gamma_{mr}=\sum_{r=1}^{R}\sum_{m=1}^{M}\pi_{ir}\gamma_{mr}\end{aligned}$$

$$= \sum_{r=1}^{R} \pi_{ir}\gamma_r \ ,$$

a sum of genetic group effects, weighted by global ancestry proportions.

#### Derivation of equivalent statement of genetic value

Here we show an alternative but equivalent expression of genetic value found using mean-centered versions of $\Lambda_{imr}^{(h)}$ and $W_{im}^{(h)}$. We start with the original definition of genetic value and first insert the mean centered allele indicator $\widetilde{W}_{im}^{(h)}$;

$$\begin{aligned}
U_i &= \sum_{m=1}^{M}\sum_{r=1}^{R}\sum_{h=1}^{2}\frac{1}{2}\left[\Lambda_{imr}^{(h)}\left(\beta_{mr}^{\text{ref}} + W_{im}^{(h)}\left(\beta_{mr}^{\text{alt}} - \beta_{mr}^{\text{ref}}\right)\right)\right] \\
&= \sum_{m=1}^{M}\sum_{r=1}^{R}\sum_{h=1}^{2}\frac{1}{2}\left[\Lambda_{imr}^{(h)}\left(\beta_{mr}^{\text{ref}} + \left(\frac{\widetilde{W}_{imr}^{(h)}}{\Lambda_{imr}^{(h)}} + p_{mr}\right)\left(\beta_{mr}^{\text{alt}} - \beta_{mr}^{\text{ref}}\right)\right)\right] \\
&= \sum_{m=1}^{M}\sum_{r=1}^{R}\sum_{h=1}^{2}\frac{1}{2}\left[\Lambda_{imr}^{(h)}\beta_{mr}^{\text{ref}} + \left(\widetilde{W}_{imr}^{(h)} + \Lambda_{imr}^{(h)}p_{mr}\right)\left(\beta_{mr}^{\text{alt}} - \beta_{mr}^{\text{ref}}\right)\right] \\
&= \sum_{m=1}^{M}\sum_{r=1}^{R}\sum_{h=1}^{2}\frac{1}{2}\left[\Lambda_{imr}^{(h)}\left(\beta_{mr}^{\text{ref}} + p_{mr}\left(\beta_{mr}^{\text{alt}} - \beta_{mr}^{\text{ref}}\right)\right) + \widetilde{W}_{imr}^{(h)}\left(\beta_{mr}^{\text{alt}} - \beta_{mr}^{\text{ref}}\right)\right] \\
&= \sum_{m=1}^{M}\sum_{r=1}^{R}\sum_{h=1}^{2}\frac{1}{2}\left[\Lambda_{imr}^{(h)}\gamma_{mr} + \widetilde{W}_{imr}^{(h)}\left(\beta_{mr}^{\text{alt}} - \beta_{mr}^{\text{ref}}\right)\right] \ .
\end{aligned}$$

For the first term in the above brackets, we split out the $R$ term in the sum over genetic groups, as follows

$$U_i = \sum_{m=1}^{M}\sum_{h=1}^{2}\frac{1}{2}\left[\sum_{r=1}^{R-1}\left(\widetilde{\Lambda}_{imr}^{(h)} + \pi_{ir}\right)\gamma_{mr} + \Lambda_{imR}^{(h)}\gamma_{mR} + \sum_{r=1}^{R}\widetilde{W}_{imr}^{(h)}\left(\beta_{mr}^{\text{alt}} - \beta_{mr}^{\text{ref}}\right)\right] \ .$$

Now, since $\Lambda_{imR}^{(h)} = 1 - \sum_{r=1}^{R-1}\Lambda_{imR}^{(h)}$ we can write

$$\begin{aligned}
U_i = \sum_{m=1}^{M}\sum_{h=1}^{2}\frac{1}{2}&\left[\sum_{r=1}^{R-1}\left(\widetilde{\Lambda}_{imr}^{(h)} + \pi_{ir}\right)\gamma_{mr} + \left(1 - \sum_{r=1}^{R-1}\Lambda_{imr}^{(h)}\right)\gamma_{mR}\right. \\
&\left. + \sum_{r=1}^{R}\widetilde{W}_{imr}^{(h)}\left(\beta_{mr}^{\text{alt}} - \beta_{mr}^{\text{ref}}\right)\right] \\
= \sum_{m=1}^{M}\sum_{h=1}^{2}\frac{1}{2}&\left[\sum_{r=1}^{R-1}\left(\widetilde{\Lambda}_{imr}^{(h)} + \pi_{ir}\right)\gamma_{mr} + \left(1 - \sum_{r=1}^{R-1}\left(\widetilde{\Lambda}_{imr}^{(h)} + \pi_{ir}\right)\right)\gamma_{mR}\right. \\
&\left. + \sum_{r=1}^{R}\widetilde{W}_{imr}^{(h)}\left(\beta_{mr}^{\text{alt}} - \beta_{mr}^{\text{ref}}\right)\right] \ .
\end{aligned}$$

Note that $1 - \sum_{r=1}^{R-1} \pi_{ir} = \pi_{iR}$, so

$$\begin{aligned} U_i = & \sum_{m=1}^{M} \sum_{h=1}^{2} \frac{1}{2} \left[ \sum_{r=1}^{R-1} \left( \widetilde{\Lambda}_{imr}^{(h)} + \pi_{ir} \right) \gamma_{mr} + \left( \pi_{iR} - \sum_{r=1}^{R-1} \widetilde{\Lambda}_{imr}^{(h)} \right) \gamma_{mR} \right. \\ & \left. + \sum_{r=1}^{R} \widetilde{W}_{imr}^{(h)} \left( \beta_{mr}^{\text{alt}} - \beta_{mr}^{\text{ref}} \right) \right] \\ = & \sum_{m=1}^{M} \sum_{h=1}^{2} \frac{1}{2} \left[ \sum_{r=1}^{R-1} \widetilde{\Lambda}_{imr}^{(h)} \left( \gamma_{mr} - \gamma_{mR} \right) + \sum_{r=1}^{R} \pi_{ir} \gamma_{mr} \right. \\ & \left. + \sum_{r=1}^{R} \widetilde{W}_{imr}^{(h)} \left( \beta_{mr}^{\text{alt}} - \beta_{mr}^{\text{ref}} \right) \right] . \end{aligned}$$

The term $\sum_{m=1}^{M} \sum_{h=1}^{2} \sum_{r=1}^{R} \frac{1}{2} \pi_{ir} \gamma_{mr}$ equals the mean value of $U_i$, so we can split it out and finally write

$$U_i = \mathsf{E}\left(U_i\right) + \sum_{m=1}^{M} \sum_{h=1}^{2} \frac{1}{2} \left[ \sum_{r=1}^{R-1} \widetilde{\Lambda}_{imr}^{(h)} \left( \gamma_{mr} - \gamma_{mR} \right) + \sum_{r=1}^{R} \widetilde{W}_{imr}^{(h)} \left( \beta_{mr}^{\text{alt}} - \beta_{mr}^{\text{ref}} \right) \right] .$$

#### Covariances between centered allele variant indicators

##### Between-locus

We calculate the covariance between centered allele variant indicators of alleles on different loci ($m \neq m'$). This holds regardless of whether $i = j$, $r = r'$ and $h = h'$ or not. First off, because $\widetilde{W}_{imr}^{(h)}$ has mean 0, we can say

$$\mathsf{Cov}\left( \widetilde{W}_{imr}^{(h)}, \widetilde{W}_{jm'r'}^{(h')} \right) = \mathsf{E}\left( \widetilde{W}_{imr}^{(h)} \widetilde{W}_{jm'r'}^{(h')} \right) - \mathsf{E}\left( \widetilde{W}_{imr}^{(h)} \right) \left( \widetilde{W}_{jm'r'}^{(h')} \right) = \mathsf{E}\left( \widetilde{W}_{imr}^{(h)} \widetilde{W}_{jm'r'}^{(h')} \right) .$$

Using the Law of total expectation, and the fact that $\widetilde{W}_{imr}^{(h)} \widetilde{W}_{jm'r'}^{(h')} = 0$ if at least one of $\Lambda_{imr}^{(h)}$ or $\Lambda_{jm'r'}^{(h')}$ are zero, we can write

$$\begin{aligned} & \mathsf{Cov}\left( \widetilde{W}_{imr}^{(h)}, \widetilde{W}_{jm'r'}^{(h')} \right) \\ & = \mathsf{E}\left( \widetilde{W}_{imr}^{(h)} \widetilde{W}_{jm'r'}^{(h')} \,\middle|\, \Lambda_{imr}^{(h)} = 1, \Lambda_{jm'r'}^{(h')} = 1 \right) \mathsf{P}\left( \Lambda_{imr}^{(h)} = 1, \Lambda_{jm'r'}^{(h')} = 1 \right) \\ & + \mathsf{E}\left( \widetilde{W}_{imr}^{(h)} \widetilde{W}_{jm'r'}^{(h')} \,\middle|\, \Lambda_{imr}^{(h)} = 1, \Lambda_{jm'r'}^{(h')} = 0 \right) \mathsf{P}\left( \Lambda_{imr}^{(h)} = 1, \Lambda_{jm'r'}^{(h')} = 0 \right) \\ & + \mathsf{E}\left( \widetilde{W}_{imr}^{(h)} \widetilde{W}_{jm'r'}^{(h')} \,\middle|\, \Lambda_{imr}^{(h)} = 0, \Lambda_{jm'r'}^{(h')} = 1 \right) \mathsf{P}\left( \Lambda_{imr}^{(h)} = 0, \Lambda_{jm'r'}^{(h')} = 1 \right) \\ & + \mathsf{E}\left( \widetilde{W}_{imr}^{(h)} \widetilde{W}_{jm'r'}^{(h')} \,\middle|\, \Lambda_{imr}^{(h)} = 0, \Lambda_{jm'r'}^{(h')} = 0 \right) \mathsf{P}\left( \Lambda_{imr}^{(h)} = 0, \Lambda_{jm'r'}^{(h')} = 0 \right) \\ & = \mathsf{E}\left( \widetilde{W}_{imr}^{(h)} \widetilde{W}_{jm'r'}^{(h')} \,\middle|\, \Lambda_{imr}^{(h)} = 1, \Lambda_{jm'r'}^{(h')} = 1 \right) \mathsf{P}\left( \Lambda_{imr}^{(h)} = 1, \Lambda_{jm'r'}^{(h')} = 1 \right) . \end{aligned}$$

We have assumed the LD to be zero, so

$$\begin{aligned}&\mathsf{Corr}\left(\widetilde{W}_{imr}^{(h)},\widetilde{W}_{jm'r'}^{(h')}\mid\Lambda_{imr}^{(h)}=1,\ \Lambda_{jm'r'}^{(h')}=1\right)\\&=\mathsf{Corr}\left(W_{im}^{(h)},W_{jm'}^{(h')}\mid\Lambda_{imr}^{(h)}=1,\ \Lambda_{jm'r'}^{(h')}=1\right)=0\ ,\end{aligned}$$

leading to

$$\begin{aligned}\mathsf{E}\left(\widetilde{W}_{imr}^{(h)}\widetilde{W}_{jm'r'}^{(h')}\,\middle|\,\Lambda_{imr}^{(h)}=1,\Lambda_{jm'r'}^{(h')}=1\right)&=\mathsf{Cov}\left(\widetilde{W}_{imr}^{(h)},\widetilde{W}_{jm'r'}^{(h')}\,\middle|\,\Lambda_{imr}^{(h)}=1,\Lambda_{jm'r'}^{(h')}=1\right)+0\times 0\\&=0\times\sqrt{p_{mr}\left(1-p_{mr}\right)p_{m'r'}\left(1-p_{m'r'}\right)}=0\end{aligned}$$

which implies

$$\mathsf{Cov}\left(\widetilde{W}_{imr}^{(h)},\widetilde{W}_{jm'r'}^{(h')}\right)=0,\quad\forall i,\ j,\ m,\ m'\neq m,\ r,\ r',\ h,\ h'\ .$$

**Within-locus, between-allele**

We follow the same logic as in the between-locus case, by noting that by assumption

$$\begin{aligned}&\mathsf{Corr}\left(\widetilde{W}_{imr}^{(h)},\widetilde{W}_{jmr'}^{(h')}\,\middle|\,\Lambda_{imr}^{(h)}=1,\ \Lambda_{jmr'}^{(h')}=1\right)\\&=\mathsf{Corr}\left(W_{im}^{(h)},W_{jm}^{(h')}\,\middle|\,\Lambda_{imr}^{(h)}=1,\ \Lambda_{jmr'}^{(h')}=1\right)=0\ .\end{aligned}$$

Thus,

$$\mathsf{Cov}\left(\widetilde{W}_{imr}^{(h)},\widetilde{W}_{jmr'}^{(h')}\right)=0,\quad\forall i,\ j,\ m,\ r,\ r',\ h,\ h'\neq h\ .$$

**Between-group**

We calculate $\mathsf{Cov}\left(\widetilde{W}_{imr}^{(h)},\widetilde{W}_{imr'}^{(h)}\right)$ with $r\neq r'$, using the definition of $\widetilde{W}_{imr}^{(h)}$, as follows

$$\begin{aligned}\mathsf{Cov}\left(\widetilde{W}_{imr}^{(h)},\widetilde{W}_{imr'}^{(h)}\right)&=\mathsf{Cov}\left(\Lambda_{imr}^{(h)}\left(W_{im}^{(h)}-p_{mr}\right),\ \Lambda_{imr'}^{(h')}\left(W_{im}^{(h)}-p_{mr'}\right)\right)\\&=\mathsf{Cov}\left(\Lambda_{imr}^{(h)}W_{im}^{(h)},\ \Lambda_{imr'}^{(h)}W_{im}^{(h)}\right)\\&-\mathsf{Cov}\left(\Lambda_{imr}^{(h)}W_{im}^{(h)},\ \Lambda_{imr'}^{(h)}\right)p_{mr'}\\&-\mathsf{Cov}\left(\Lambda_{imr}^{(h)},\ \Lambda_{imr'}^{(h)}W_{im}^{(h)}\right)p_{mr}\\&+\mathsf{Cov}\left(\Lambda_{imr}^{(h)},\ \Lambda_{imr'}^{(h)}\right)p_{mr}p_{mr'}\end{aligned}$$

$$
\begin{aligned}
&= \mathsf{E}\left(\Lambda^{(h)}_{imr} W^{(h)}_{im} \Lambda^{(h)}_{imr'} W^{(h)}_{im}\right) \\
&- \mathsf{E}\left(\Lambda^{(h)}_{imr} W^{(h)}_{im}\right) \mathsf{E}\left(\Lambda^{(h)}_{imr'} W^{(h)}_{im}\right) \\
&- \left[\mathsf{E}\left(\Lambda^{(h)}_{imr} W^{(h)}_{im} \Lambda^{(h)}_{imr'}\right) - \mathsf{E}\left(\Lambda^{(h)}_{imr} W^{(h)}_{im}\right) \mathsf{E}\left(\Lambda^{(h)}_{imr'}\right)\right] p_{mr'} \\
&- \left[\mathsf{E}\left(\Lambda^{(h)}_{imr} \Lambda^{(h)}_{imr'} W^{(h)}_{im}\right) - \mathsf{E}\left(\Lambda^{(h)}_{imr}\right) \mathsf{E}\left(\Lambda^{(h)}_{imr'} W^{(h)}_{im}\right)\right] p_{mr} \\
&+ \left[\mathsf{E}\left(\Lambda^{(h)}_{imr} \Lambda^{(h)}_{imr'}\right) - \mathsf{E}\left(\Lambda^{(h)}_{imr}\right) \mathsf{E}\left(\Lambda^{(h)}_{imr'}\right)\right] p_{mr} p_{mr'} \ .
\end{aligned}
$$

Since at least one of $\Lambda^{(h)}_{imr}$ or $\Lambda^{(h)}_{imr'}$ equal zero, all the expectations above containing a product between these two will also equal zero, so we have

$$
\begin{aligned}
\mathsf{Cov}\left(\widetilde{W}^{(h)}_{imr}, \widetilde{W}^{(h)}_{imr'}\right) &= -\mathsf{E}\left(\Lambda^{(h)}_{imr} W^{(h)}_{im}\right) \mathsf{E}\left(\Lambda^{(h)}_{imr'} W^{(h)}_{im}\right) \\
&+ \mathsf{E}\left(\Lambda^{(h)}_{imr} W^{(h)}_{im}\right) \mathsf{E}\left(\Lambda^{(h)}_{imr'}\right) p_{mr'} \\
&+ \mathsf{E}\left(\Lambda^{(h)}_{imr}\right) \mathsf{E}\left(\Lambda^{(h)}_{imr'} W^{(h)}_{im}\right) p_{mr} \\
&- \mathsf{E}\left(\Lambda^{(h)}_{imr}\right) \mathsf{E}\left(\Lambda^{(h)}_{imr'}\right) p_{mr} p_{mr'} \\
&= (-1+1+1-1) \pi_{ir} p_{mr} \pi_{ir'} p_{mr'} \\
&= 0 \ , \quad \forall i,\ m,\ r,\ r' \neq r,\ h \ .
\end{aligned}
$$

Since this holds for the case where the alleles have "the most in common" ($i = j,\ h = h'$), we assume that the same also holds for $i \neq j$ and $h \neq h'$, that is, $\mathsf{Cov}\left(\widetilde{W}^{(h)}_{imr}, \widetilde{W}^{(h')}_{jmr'}\right) = 0$ for all $i, j, m, r, r' \neq r, h, h'$.

**Within-locus, within-group, within-allele**

We calculate the covariance between allele variants on the same allele $h$ in the same locus $m$. By the same logic as in the between-locus and between-allele cases, we obtain

$$
\begin{aligned}
&\mathsf{Cov}\left(\widetilde{W}^{(h)}_{imr}, \widetilde{W}^{(h)}_{jmr}\right) \\
&= \mathsf{E}\left(\widetilde{W}^{(h)}_{imr} \widetilde{W}^{(h)}_{jmr} \,\middle|\, \Lambda^{(h)}_{imr} = 1, \Lambda^{(h)}_{jmr} = 1\right) \mathsf{P}\left(\Lambda^{(h)}_{imr} = 1, \Lambda^{(h)}_{jmr} = 1\right) \ .
\end{aligned}
$$

Since $\Lambda^{(h)}_{imr}$ and $\Lambda^{(h)}_{jmr}$ are indicator variables that only take the values of 1 and 0, we can observe that $\mathsf{P}\left(\Lambda^{(h)}_{imr} = 1, \Lambda^{(h)}_{jmr} = 1\right) = \mathsf{P}\left(\Lambda^{(h)}_{imr} = 1, \Lambda^{(h)}_{jmr} = 1\right) = \mathsf{E}\left(\Lambda^{(h)}_{imr} \Lambda^{(h)}_{jmr}\right)$, which is also the definition of $\theta_{ij}^{(r)}$. Further note that

$$\begin{aligned}&\mathsf{Corr}\left(\widetilde{W}_{imr}^{(h)},\widetilde{W}_{jmr}^{(h)}\,\middle|\,\Lambda_{imr}^{(h)}=1,\ \Lambda_{jmr}^{(h)}=1\right)\\&=\mathsf{Corr}\left(W_{im}^{(h)},W_{jm}^{(h)}\,\middle|\,\Lambda_{imr}^{(h)}=1,\ \Lambda_{jmr}^{(h)}=1\right)=\Gamma_{ij}^{(r)}\end{aligned}$$

and that

$$\begin{aligned}&\mathsf{Var}\left(\widetilde{W}_{imr}^{(h)}\,\middle|\,\Lambda_{imr}^{(h)}=1,\ \Lambda_{jmr}^{(h)}=1\right)\\&=\mathsf{Var}\left(W_{im}^{(h)}\,\middle|\,\Lambda_{imr}^{(h)}=1,\ \Lambda_{jmr}^{(h)}=1\right)\\&=p_{mr}\left(1-p_{mr}\right)\ ,\end{aligned}$$

with the same result for the conditional variance of $\widetilde{W}_{jmr}^{(h)}$. With this in hand we can write

$$\begin{aligned}\mathsf{E}\left(\widetilde{W}_{imr}^{(h)}\widetilde{W}_{jmr}^{(h)}\,\middle|\,\Lambda_{imr}^{(h)}=1,\Lambda_{jmr}^{(h)}=1\right)&=\mathsf{Cov}\left(\widetilde{W}_{imr}^{(h)},\widetilde{W}_{jmr}^{(h)}\,\middle|\,\Lambda_{imr}^{(h)}=1,\Lambda_{jmr}^{(h)}=1\right)\\&=\Gamma_{ij}^{(r)}p_{mr}\left(1-p_{mr}\right)\end{aligned}$$

by the definition of correlation. Then finally, we arrive at

$$\begin{aligned}&\mathsf{Cov}\left(\widetilde{W}_{imr}^{(h)},\widetilde{W}_{jmr}^{(h)}\right)\\&=\mathsf{E}\left(\widetilde{W}_{imr}^{(h)}\widetilde{W}_{jmr}^{(h)}\,\middle|\,\Lambda_{imr}^{(h)}=1,\Lambda_{jmr}^{(h)}=1\right)\mathsf{P}\left(\Lambda_{imr}^{(h)}=1,\Lambda_{jmr}^{(h)}=1\right)\\&=\theta_{ij}^{(r)}\Gamma_{ij}^{(r)}p_{mr}\left(1-p_{mr}\right)\quad\forall i,\ j,\ m,\ r,\ h\ .\end{aligned}$$

#### Covariances between local ancestry indicators

We find the covariance between local ancestry indicators. Note that for the within-locus, within-allele case we have

$$\mathsf{Cov}\left(\Lambda_{imr}^{(h)},\Lambda_{jmr'}^{(h)}\right)=\Delta_{ij}^{(rr')}=\theta_{ij}^{(rr')}-\pi_{ir}\pi_{jr'},\quad\forall\ i,\ j,\ m,\ r,\ r',\ h$$

by definition. We now find the covariance between local ancestry indicators on different loci, first within groups and then between groups.

**Between-locus, within-group**

The between-locus, within-group case was already shown in the Supplementary Material 1 of Rio *et al.* (2020) (they do not need the between-group case), but we include it here for completeness, using our own notation. From the definition of $\widetilde{\Lambda}_{imr}^{(h)}$, we can say

$$\begin{aligned}
\mathsf{Cov}\left(\widetilde{\Lambda}_{imr}^{(h)}, \widetilde{\Lambda}_{jm'r}^{(h')}\right) &= \mathsf{Cov}\left(\Lambda_{imr}^{(h)}, \Lambda_{jm'r}^{(h')}\right) \\
&= \mathsf{E}\left(\Lambda_{imr}^{(h)}\Lambda_{jm'r}^{(h')}\right) - \mathsf{E}\left(\Lambda_{imr}^{(h)}\right)\mathsf{E}\left(\Lambda_{jm'r}^{(h')}\right) \\
&= \mathsf{P}\left(\Lambda_{imr}^{(h)} = 1, \Lambda_{jm'r}^{(h')} = 1\right) - \pi_{ir}\pi_{jr} \\
&= \mathsf{P}\left(\Lambda_{imr}^{(h)} = 1 \,\middle|\, \Lambda_{jm'r}^{(h')} = 1\right)\mathsf{P}\left(\Lambda_{jm'r}^{(h')} = 1\right) - \pi_{ir}\pi_{jr} \\
&= \mathsf{P}\left(\Lambda_{imr}^{(h)} = 1 \,\middle|\, \Lambda_{jm'r}^{(h')} = 1\right)\pi_{jr} - \pi_{ir}\pi_{jr} \ . \qquad \text{(S1)}
\end{aligned}$$

We now consider $\mathsf{P}\left(\Lambda_{imr}^{(h)} = 1 \,\middle|\, \Lambda_{jm'r}^{(h')} = 1\right)$. We use the law of total expectation by conditioning on $\Lambda_{im'r}^{(h')}$, so that

$$\begin{aligned}
&\mathsf{P}\left(\Lambda_{imr}^{(h)} = 1 \,\middle|\, \Lambda_{jm'r}^{(h')} = 1\right) \\
=&\mathsf{P}\left(\Lambda_{imr}^{(h)} = 1 \,\middle|\, \Lambda_{im'r}^{(h')} = 1, \Lambda_{jm'r}^{(h')} = 1\right)\mathsf{P}\left(\Lambda_{im'r}^{(h')} = 1 \,\middle|\, \Lambda_{jm'r}^{(h')} = 1\right) \\
+&\mathsf{P}\left(\Lambda_{imr}^{(h)} = 1 \,\middle|\, \Lambda_{im'r}^{(h')} = 0, \Lambda_{jm'r}^{(h')} = 1\right)\mathsf{P}\left(\Lambda_{im'r}^{(h')} = 0 \,\middle|\, \Lambda_{jm'r}^{(h')} = 1\right) \\
=&\mathsf{P}\left(\Lambda_{imr}^{(h)} = 1 \,\middle|\, \Lambda_{im'r}^{(h')} = 1, \Lambda_{jm'r}^{(h')} = 1\right)\frac{\mathsf{P}\left(\Lambda_{im'r}^{(h')} = 1, \Lambda_{jm'r}^{(h')} = 1\right)}{\mathsf{P}\left(\Lambda_{jm'r}^{(h')} = 1\right)} \\
+&\mathsf{P}\left(\Lambda_{imr}^{(h)} = 1 \,\middle|\, \Lambda_{im'r}^{(h')} = 0, \Lambda_{jm'r}^{(h')} = 1\right)\frac{\mathsf{P}\left(\Lambda_{im'r}^{(h')} = 0, \Lambda_{jm'r}^{(h')} = 1\right)}{\mathsf{P}\left(\Lambda_{jm'r}^{(h')} = 1\right)} \\
=&\mathsf{P}\left(\Lambda_{imr}^{(h)} = 1 \,\middle|\, \Lambda_{im'r}^{(h')} = 1, \Lambda_{jm'r}^{(h')} = 1\right)\frac{\theta_{ij}^{(r)}}{\pi_{jr}} \\
+&\mathsf{P}\left(\Lambda_{imr}^{(h)} = 1 \,\middle|\, \Lambda_{im'r}^{(h')} = 0, \Lambda_{jm'r}^{(h')} = 1\right)\frac{\pi_{jr} - \theta_{ij}^{(r)}}{\pi_{jr}} \ . \qquad \text{(S2)}
\end{aligned}$$

Consider that $\mathsf{P}\left(\Lambda_{imr}^{(h)} = 1 \,\middle|\, \Lambda_{im'r}^{(h')} = 1, \Lambda_{jm'r}^{(h')} = 1\right)$ will be the global ancestry proportion of $i$ in group $r$, on a shrunken set of alleles, since 1 out of the $2M$ alleles are already known to be in group $r$. Thus, we can say

$$\mathsf{P}\left(\Lambda_{imr}^{(h)} = 1 \,\middle|\, \Lambda_{im'r}^{(h')} = 1, \Lambda_{jm'r}^{(h')} = 1\right) = \frac{\pi_{ir} - \frac{1}{2M}}{1 - \frac{1}{2M}} = \frac{2M\pi_{ir} - 1}{2M - 1} \ . \qquad \text{(S3)}$$

Similarly, $\mathsf{P}\left(\Lambda_{imr}^{(h)}=1 \,\middle|\, \Lambda_{im'r}^{(h')}=0, \Lambda_{jm'r}^{(h')}=1\right)$ is the global ancestry proportion on the same shrunken set of alleles, but group $r$ is known not to have been shrunk, so

$$\mathsf{P}\left(\Lambda_{imr}^{(h)}=1 \,\middle|\, \Lambda_{im'r}^{(h')}=0, \Lambda_{jm'r}^{(h')}=1\right)=\frac{\pi_{ir}}{1-\frac{1}{2M}}=\frac{2M\pi_{ir}}{2M-1} \ . \tag{S4}$$

Insert (S3) and (S4) into (S2), which we insert into (S1) to end up at

$$\begin{aligned}
\mathsf{Cov}\left(\widetilde{\Lambda}_{imr}^{(h)}, \widetilde{\Lambda}_{jm'r}^{(h')}\right) &= \left(\frac{2M\pi_{ir}-1}{2M-1}\times\frac{\theta_{ij}^{(r)}}{\pi_{jr}}+\frac{2M\pi_{ir}}{2M-1}\times\frac{\pi_{jr}-\theta_{ij}^{(r)}}{\pi_{jr}}\right)\pi_{jr}-\pi_{ir}\pi_{jr} \\
&= \frac{\left(2M\pi_{ir}-1\right)\theta_{ij}^{(r)}+2M\pi_{ir}\left(\pi_{jr}-\theta_{ij}^{(r)}\right)}{2M-1}-\pi_{ir}\pi_{jr} \\
&= \frac{2M\pi_{ir}\pi_{jr}-\theta_{ij}^{(r)}}{2M-1}-\pi_{ir}\pi_{jr}=\frac{\pi_{ir}\pi_{jr}-\theta_{ij}^{(r)}}{2M-1} \\
&= -\frac{\Delta_{ij}^{(r)}}{2M-1}, \quad \forall i,\ j,\ m,\ m'\neq m,\ r,\ h,\ h' \ .
\end{aligned}$$

**Between-locus, between-group**

As for $\mathsf{Cov}\left(\widetilde{\Lambda}_{imr}^{(h)}, \widetilde{\Lambda}_{jm'r'}^{(h')}\right)$ for $m \neq m'$ and $r \neq r'$, we find

$$\mathsf{Cov}\left(\widetilde{\Lambda}_{imr}^{(h)}, \widetilde{\Lambda}_{jm'r'}^{(h')}\right)=\mathsf{P}\left(\Lambda_{imr}^{(h)}=1 \,\middle|\, \Lambda_{jm'r'}^{(h')}=1\right)\pi_{jr'}-\pi_{ir}\pi_{jr'} \ , \tag{S5}$$

similarly to the approach when $r=r'$. When considering $\mathsf{P}\left(\Lambda_{imr}^{(h)}=1 \,\middle|\, \Lambda_{jm'r'}^{(h')}\right)$, we this time instead condition on $\Lambda_{im'r'}^{(h')}$ in the law of total expectation, so that

$$\begin{aligned}
&\mathsf{P}\left(\Lambda_{imr}^{(h)}=1 \,\middle|\, \Lambda_{jm'r'}^{(h')}=1\right) \\
=&\mathsf{P}\left(\Lambda_{imr}^{(h)}=1 \,\middle|\, \Lambda_{im'r'}^{(h')}=1, \Lambda_{jm'r'}^{(h')}=1\right)\frac{\theta_{ij}^{(r')}}{\pi_{jr'}} \\
+&\mathsf{P}\left(\Lambda_{imr}^{(h)}=1 \,\middle|\, \Lambda_{im'r'}^{(h')}=0, \Lambda_{jm'r'}^{(h')}=1\right)\frac{\pi_{jr'}-\theta_{ij}^{(r')}}{\pi_{jr'}} \ .
\end{aligned} \tag{S6}$$

Again, $\mathsf{P}\left(\Lambda_{imr}^{(h)}=1 \,\middle|\, \Lambda_{im'r'}^{(h')}=1, \Lambda_{jm'r'}^{(h')}=1\right)$ will be the global ancestry proportion of $i$ in group $r$, on a shrunken set of alleles, where 1 out of the $2M$ alleles are already known to be in another group $r'$. Thus,

$$\mathsf{P}\left(\Lambda_{imr}^{(h)}=1 \,\middle|\, \Lambda_{im'r'}^{(h')}=1, \Lambda_{jm'r'}^{(h')}=1\right)=\frac{\pi_{ir}}{1-\frac{1}{2M}} \ . \tag{S7}$$

When it comes to $\mathsf{P}\left(\Lambda^{(h)}_{imr}=1 \,\middle|\, \Lambda^{(h')}_{im'r'}=0, \Lambda^{(h')}_{jm'r'}=1\right)$, all we know is that $i$'s loci allele $h'$ at $m'$ is *not* in group $r'$. Thus, it can either be in group $r$, or in neither group $r$ or $r'$. We weight the results from each of these cases by their respective probability, so

$$\begin{aligned}&\mathsf{P}\left(\Lambda^{(h)}_{imr}=1 \,\middle|\, \Lambda^{(h')}_{im'r'}=0, \Lambda^{(h')}_{jm'r'}=1\right)\\ &=\frac{\theta^{(rr')}_{ij}}{\pi_{jr'}-\theta^{(r')}_{ij}}\times\frac{\pi_{ir}-\frac{1}{2M}}{1-\frac{1}{2M}}+\frac{\pi_{jr'}-\theta^{(r')}_{ij}-\theta^{(rr')}_{ij}}{\pi_{jr'}-\theta^{(r')}_{ij}}\times\frac{\pi_{ir}}{1-\frac{1}{2M}}\end{aligned} \tag{S8}$$

We insert (S7) and (S8) into (S6), which we insert into (S5) to end up at

$$\begin{aligned}&\mathsf{Cov}\left(\widetilde{\Lambda}^{(h)}_{imr}, \widetilde{\Lambda}^{(h')}_{jm'r'}\right)\\ =\;&\left[\left(\frac{\theta^{(rr')}_{ij}}{\pi_{jr'}-\theta^{(r')}_{ij}}\times\frac{\pi_{ir}-\frac{1}{2M}}{1-\frac{1}{2M}}+\frac{\pi_{jr'}-\theta^{(r')}_{ij}-\theta^{(rr')}_{ij}}{\pi_{jr'}-\theta^{(r')}_{ij}}\times\frac{\pi_{ir}}{1-\frac{1}{2M}}\right)\times\frac{\pi_{jr'}-\theta^{(r')}_{ij}}{\pi_{jr'}}\right.\\ &\left.+\frac{\pi_{ir}}{1-\frac{1}{2M}}\times\frac{\theta^{(r')}_{ij}}{\pi_{jr'}}\right]\pi_{jr'}-\pi_{ir}\pi_{jr'}\\ =\;&\left[\frac{\theta^{(rr')}_{ij}\left(\pi_{ir}-\frac{1}{2M}\right)+\left(\pi_{jr'}-\theta^{(r')}_{ij}-\theta^{(rr')}_{ij}\right)\pi_{ir}+\pi_{ir}\theta^{(r')}_{ij}}{1-\frac{1}{2M}}\right]-\pi_{ir}\pi_{jr'}\\ =\;&\left[\frac{\pi_{ir}\pi_{jr'}-\theta^{(rr')}_{ij}\frac{1}{2M}}{1-\frac{1}{2M}}\right]-\pi_{ir}\pi_{jr'}=\frac{2M\pi_{ir}\pi_{jr'}-\theta^{(rr')}_{ij}-(2M-1)\,\pi_{ir}\pi_{jr'}}{2M-1}\\ =\;&\frac{\pi_{ir}\pi_{jr'}-\theta^{(rr')}_{ij}}{2M-1}=-\frac{\Delta^{(rr')}_{ij}}{2M-1},\quad \forall i,\ j,\ m,\ m'\neq m,\ r,\ r'\neq r,\ h,\ h'\ .\end{aligned}$$

**Within-locus, between allele**

The case $m=m'$, $h\neq h'$ has a similar derivation to the between locus case, both within and between groups. So

$$\mathsf{Cov}\left(\widetilde{\Lambda}^{(h)}_{imr}, \widetilde{\Lambda}^{(h')}_{jmr'}\right)=-\frac{\Delta^{(rr')}_{ij}}{2M-1},\quad \forall i,\ j,\ m,\ r,\ r',\ h,\ h'\neq h,\ .$$

#### Covariances between allele variant indicators and local ancestry indicators

We calculate the covariance $\mathsf{Cov}\left(\widetilde{\Lambda}^{(h)}_{imr}, \widetilde{W}^{(h')}_{jm'r'}\right)$, using the definition of the centered variables as follows.

$$\begin{aligned}&\mathsf{Cov}\left(\widetilde{\Lambda}^{(h)}_{imr}, \widetilde{W}^{(h')}_{jm'r'}\right)\\ =&\mathsf{Cov}\left(\Lambda^{(h)}_{imr}, \Lambda^{(h')}_{jm'r'}W^{(h')}_{jm'}\right)-\mathsf{Cov}\left(\Lambda^{(h)}_{imr}, \Lambda^{(h')}_{jm'r'}\right)p_{m'r'}\end{aligned}$$

$$=\mathsf{E}\left(\Lambda_{imr}^{(h)}\Lambda_{jm'r'}^{(h')}W_{jm'}^{(h')}\right)-\mathsf{E}\left(\Lambda_{imr}^{(h)}\right)\mathsf{E}\left(\Lambda_{jm'r'}^{(h')}W_{jm'}^{(h')}\right)-\mathsf{Cov}\left(\Lambda_{imr}^{(h)},\Lambda_{jm'r'}^{(h')}\right)\times p_{m'r'}$$
$$=\mathsf{P}\left(\Lambda_{imr}^{(h)}=1,\Lambda_{jm'r'}^{(h')}=1,W_{jm'}^{(h')}=1\right)-\pi_{ir}\pi_{jr'}p_{m'r'}-\mathsf{Cov}\left(\Lambda_{imr}^{(h)},\Lambda_{jm'r'}^{(h')}\right)\times p_{m'r'}\ .$$

Note that

$$\begin{aligned}
&\mathsf{P}\left(\Lambda_{imr}^{(h)}=1,\Lambda_{jm'r'}^{(h')}=1,W_{jm'}^{(h')}=1\right)\\
&=\mathsf{P}\left(W_{jm'}^{(h')}=1\,\middle|\,\Lambda_{imr}^{(h)}=1,\Lambda_{jm'r'}^{(h')}=1\right)\times\mathsf{P}\left(\Lambda_{imr}^{(h)}=1,\Lambda_{jm'r'}^{(h')}=1\right)\\
&=p_{m'r'}\times\mathsf{E}\left(\Lambda_{imr}^{(h)}\Lambda_{jm'r'}^{(h')}\right)\\
&=p_{m'r'}\times\left(\mathsf{Cov}\left(\Lambda_{imr}^{(h)},\Lambda_{jm'r'}^{(h')}\right)+\mathsf{E}\left(\Lambda_{imr}^{(h)}\right)\mathsf{E}\left(\Lambda_{jm'r'}^{(h')}\right)\right)\\
&=p_{m'r'}\times\left(\mathsf{Cov}\left(\Lambda_{imr}^{(h)},\Lambda_{jm'r'}^{(h')}\right)+\pi_{ir}\pi_{jr'}\right)\ .
\end{aligned}$$

Thus

$$\mathsf{Cov}\left(\widetilde{\Lambda}_{imr}^{(h)},\widetilde{W}_{jm'r'}^{(h')}\right)=0,\quad\forall i,\ j,\ m,\ m',\ r,\ r',\ h,\ h'\ .$$

#### Covariance between total genetic values

To recap, the only nonzero covariances are

- $\mathsf{Cov}\left(\widetilde{W}_{imr}^{(h)},\widetilde{W}_{jmr}^{(h')}\right)=\theta_{ij}^{(r)}\Gamma_{ij}^{(r)}p_{mr}\left(1-p_{mr}\right)$,
- $\mathsf{Cov}\left(\widetilde{\Lambda}_{imr}^{(h)},\widetilde{\Lambda}_{jmr'}^{(h')}\right)=\Delta_{ij}^{(rr')}$,
- $\mathsf{Cov}\left(\widetilde{\Lambda}_{imr}^{(h)},\widetilde{\Lambda}_{jm'r}^{(h')}\right)=-\frac{\Delta_{ij}^{(rr')}}{2M-1},\quad(m\neq m')$,
- $\mathsf{Cov}\left(\widetilde{\Lambda}_{imr}^{(h)},\widetilde{\Lambda}_{jmr}^{(h')}\right)=-\frac{\Delta_{ij}^{(rr')}}{2M-1},\quad(h\neq h')$,

which we will use to find the covariance between the total genetic values of two individuals

$$\begin{aligned}
&\mathsf{Cov}(U_i,U_j\mid\boldsymbol{\pi}_i,\boldsymbol{\pi}_j,\boldsymbol{\theta}_{ij},\boldsymbol{\Gamma}_{ij})\\
&=\mathsf{Cov}\left(\sum_{m=1}^{M}\sum_{r=1}^{R-1}\sum_{h=1}^{2}\frac{1}{2}\widetilde{\Lambda}_{imr}^{(h)}\left(\gamma_{mr}-\gamma_{mR}\right),\sum_{m=1}^{M}\sum_{r=1}^{R-1}\sum_{h=1}^{2}\frac{1}{2}\widetilde{\Lambda}_{jmr}^{(h)}\left(\gamma_{mr}-\gamma_{mR}\right)\right)\\
&+\mathsf{Cov}\left(\sum_{m=1}^{M}\sum_{r=1}^{R}\sum_{h=1}^{2}\frac{1}{2}\widetilde{W}_{imr}^{(h)}\left(\beta_{mr}^{\text{alt}}-\beta_{mr}^{\text{ref}}\right),\sum_{m=1}^{M}\sum_{r=1}^{R}\sum_{h=1}^{2}\frac{1}{2}\widetilde{W}_{jmr}^{(h)}\left(\beta_{mr}^{\text{alt}}-\beta_{mr}^{\text{ref}}\right)\right)\\
&=\sum_{m=1}^{M}\sum_{m'=1}^{M}\sum_{r=1}^{R-1}\sum_{r'=1}^{R-1}\frac{\mathsf{Cov}\left(\widetilde{\Lambda}_{imr}^{(1)}+\widetilde{\Lambda}_{imr}^{(2)},\widetilde{\Lambda}_{jm'r'}^{(1)}+\widetilde{\Lambda}_{jm'r'}^{(2)}\right)}{4}\left(\gamma_{mr}-\gamma_{mR}\right)\left(\gamma_{m'r'}-\gamma_{m'R}\right)
\end{aligned}$$

$$
\begin{aligned}
&+ \sum_{m=1}^{M}\sum_{m'=1}^{M}\sum_{r=1}^{R}\sum_{r'=1}^{R} \frac{\mathsf{Cov}\left(\widetilde{W}_{imr}^{(1)}+\widetilde{W}_{imr}^{(2)}, \widetilde{W}_{jm'r'}^{(1)}+\widetilde{W}_{jm'r'}^{(2)}\right)}{4} \left(\beta_{mr}^{\text{alt}}-\beta_{mr}^{\text{ref}}\right)\left(\beta_{m'r'}^{\text{alt}}-\beta_{m'r'}^{\text{ref}}\right) \\
&= \sum_{m=1}^{M}\sum_{r=1}^{R-1}\sum_{r'=1}^{R-1} \frac{1}{4}\left(2\times\Delta_{ij}^{(rr')} - 2\times\frac{\Delta_{ij}^{(rr')}}{2M-1}\right)\left(\gamma_{mr}-\gamma_{mR}\right)\left(\gamma_{mr'}-\gamma_{mR}\right) \\
&- \sum_{m=1}^{M}\sum_{m'\neq m}^{M}\sum_{r=1}^{R-1}\sum_{r'=1}^{R-1} \frac{4}{4}\frac{\Delta_{ij}^{(rr')}}{2M-1}\left(\gamma_{mr}-\gamma_{mR}\right)\left(\gamma_{m'r'}-\gamma_{m'R}\right) \\
&+ \sum_{m=1}^{M}\sum_{r=1}^{R} \frac{1}{4}\left(2\times\theta_{ij}^{(r)}\Gamma_{ij}^{(r)}p_{mr}\left(1-p_{mr}\right)+2\times 0\right)\left(\beta_{mr}^{\text{alt}}-\beta_{mr}^{\text{ref}}\right)^2 \ .
\end{aligned}
$$

We note that $\sum_{m=1}^{M}\sum_{m'\neq m}^{M} a_{mm'} = \sum_{m=1}^{M}\sum_{m'=1}^{M} a_{mm'} - \sum_{m=1}^{M} a_{mm}$, so

$$
\begin{aligned}
&\mathsf{Cov}(U_i, U_j \mid \boldsymbol{\pi}_i, \boldsymbol{\pi}_j,\ \boldsymbol{\theta}_{ij}, \boldsymbol{\Gamma}_{ij}) \\
&= \sum_{r=1}^{R-1}\sum_{r'=1}^{R-1}\Delta_{ij}^{(rr')} \quad \Bigg[\frac{1}{2}\left(1-\frac{1}{2M-1}\right)\sum_{m=1}^{M}\left(\gamma_{mr}-\gamma_{mR}\right)\left(\gamma_{mr'}-\gamma_{mR}\right) \\
&\qquad + \frac{1}{2M-1}\sum_{m=1}^{M}\left(\gamma_{mr}-\gamma_{mR}\right)\left(\gamma_{mr'}-\gamma_{mR}\right) \\
&\qquad - \frac{1}{2M-1}\sum_{m=1}^{M}\sum_{m'=1}^{M}\left(\gamma_{mr}-\gamma_{mR}\right)\left(\gamma_{m'r'}-\gamma_{m'R}\right)\Bigg] \\
&+ \sum_{m=1}^{M}\sum_{r=1}^{R}\frac{1}{2}\theta_{ij}^{(r)}\Gamma_{ij}^{(r)}p_{mr}\left(1-p_{mr}\right)\left(\beta_{mr}^{\text{alt}}-\beta_{mr}^{\text{ref}}\right)^2 \\
&= \sum_{r=1}^{R-1}\sum_{r'=1}^{R-1}\Delta_{ij}^{(rr')} \quad \Bigg[\frac{M}{2M-1}\sum_{m=1}^{M}\left(\gamma_{mr}-\gamma_{mR}\right)\left(\gamma_{mr'}-\gamma_{mR}\right) \\
&\qquad - \frac{1}{2M-1}\sum_{m=1}^{M}\sum_{m'=1}^{M}\left(\gamma_{mr}-\gamma_{mR}\right)\left(\gamma_{m'r'}-\gamma_{m'R}\right)\Bigg] \\
&+ \sum_{r=1}^{R}\theta_{ij}^{(r)}\Gamma_{ij}^{(r)}\sigma_{\mathrm{G}_r}^2 \ .
\end{aligned}
$$

Now note that $\sum_{m=1}^{M}\sum_{m'=1}^{M} a_m b_{m'} = \left(\sum_{m=1}^{M} a_m\right)\left(\sum_{m'=1}^{M} b_{m'}\right)$, so that

$$
\begin{aligned}
&\mathsf{Cov}(U_i, U_j \mid \boldsymbol{\pi}_i, \boldsymbol{\pi}_j, \boldsymbol{\theta}_{ij}, \boldsymbol{\Gamma}_{ij}) \\
&= \sum_{r=1}^{R-1}\sum_{r'=1}^{R-1}\Delta_{ij}^{(rr')} \quad \Bigg[\frac{M}{2M-1}\sum_{m=1}^{M}\left(\gamma_{mr}-\gamma_{mR}\right)\left(\gamma_{mr'}-\gamma_{mR}\right) \\
&\qquad - \frac{1}{2M-1}\left(\gamma_{r}-\gamma_{R}\right)\left(\gamma_{r'}-\gamma_{R}\right)\Bigg] \\
&+ \sum_{r=1}^{R}\theta_{ij}^{(r)}\Gamma_{ij}^{(r)}\sigma_{\mathrm{G}_r}^2 \ .
\end{aligned}
$$

Note that the contents of the above brackets can be rewritten as segregation variance terms because

$$
\begin{aligned}
&\sum_{m=1}^{M} \left(\gamma_{mr} - \gamma_{mR}\right)\left(\gamma_{mr'} - \gamma_{mR}\right) \\
&= \sum_{m=1}^{M} \left[\gamma_{mr}\gamma_{mr'} - \gamma_{mr}\gamma_{mR} - \gamma_{mr'}\gamma_{mR} + \gamma_{mR}^2\right] \\
&= \sum_{m=1}^{M} \left[-\frac{1}{2}\left(\gamma_{mr} - \gamma_{mr'}\right)^2 + \frac{1}{2}\gamma_{mr}^2 + \frac{1}{2}\gamma_{mr'}^2 - \gamma_{mr}\gamma_{mR} - \gamma_{mr'}\gamma_{mR} + \gamma_{mR}^2\right] \\
&= \frac{1}{2}\sum_{m=1}^{M} \left[\left(\gamma_{mr} - \gamma_{mR}\right)^2 + \left(\gamma_{mr'} - \gamma_{mR}\right)^2 - \left(\gamma_{mr} - \gamma_{mr'}\right)^2\right] \ ,
\end{aligned}
$$

and similarly

$$
\begin{aligned}
&\left(\gamma_{r} - \gamma_{R}\right)\left(\gamma_{r'} - \gamma_{R}\right) \\
&= \frac{1}{2}\left[\left(\gamma_{r} - \gamma_{R}\right)^2 + \left(\gamma_{r'} - \gamma_{R}\right)^2 - \left(\gamma_{r} - \gamma_{r'}\right)^2\right] \ ,
\end{aligned}
$$

and thus

$$
\begin{aligned}
&\mathsf{Cov}(U_i, U_j \mid \boldsymbol{\pi}_i, \boldsymbol{\pi}_j, \boldsymbol{\theta}_{ij}, \boldsymbol{\Gamma}_{ij}) \\
&= \frac{1}{2}\sum_{r=1}^{R-1}\sum_{r'=1}^{R-1} \Delta_{ij}^{(rr')} \Bigg[\frac{M}{2M-1}\sum_{m=1}^{M}\left(\gamma_{mr} - \gamma_{mR}\right)^2 - \frac{1}{2M-1}\left(\gamma_{r} - \gamma_{R}\right)^2 \\
&\qquad + \frac{M}{2M-1}\sum_{m=1}^{M}\left(\gamma_{mr'} - \gamma_{mR}\right)^2 - \frac{1}{2M-1}\left(\gamma_{r'} - \gamma_{R}\right)^2 \\
&\qquad - \left(\frac{M}{2M-1}\sum_{m=1}^{M}\left(\gamma_{mr} - \gamma_{mr'}\right)^2 - \frac{1}{2M-1}\left(\gamma_{r} - \gamma_{r'}\right)^2\right)\Bigg] \\
&+ \sum_{r=1}^{R} \theta_{ij}^{(r)} \Gamma_{ij}^{(r)} \sigma_{\mathrm{G}_r}^2 \\
&= \frac{1}{2}\sum_{r=1}^{R-1}\sum_{r'=1}^{R-1} \Delta_{ij}^{(rr')} \left[\sigma_{\mathrm{S}_{rR}}^2 + \sigma_{\mathrm{S}_{r'R}}^2 - \sigma_{\mathrm{S}_{rr'}}^2\right] + \sum_{r=1}^{R} \theta_{ij}^{(r)} \Gamma_{ij}^{(r)} \sigma_{\mathrm{G}_r}^2 \ .
\end{aligned}
$$

Now let $\mathcal{R}$ be the set $\{1, \ldots, R\}$. We note that per definition $\sigma_{\mathrm{S}_{rr}}^2 = 0$ and $\sigma_{\mathrm{S}_{rr'}}^2 = \sigma_{\mathrm{S}_{r'r}}^2$, so we can simplify to

$$
\begin{aligned}
&\mathsf{Cov}(U_i, U_j \mid \boldsymbol{\pi}_i, \boldsymbol{\pi}_j, \boldsymbol{\theta}_{ij}, \boldsymbol{\Gamma}_{ij}) \\
&= \sum_{r=1}^{R-1} \Delta_{ij}^{(r)} \sigma_{\mathrm{S}_{rR}}^2 + \frac{1}{2}\sum_{r=1}^{R-2}\sum_{r'=r+1}^{R-1} \left(\Delta_{ij}^{(rr')} + \Delta_{ij}^{(r'r)}\right)\left[\sigma_{\mathrm{S}_{rR}}^2 + \sigma_{\mathrm{S}_{r'R}}^2 - \sigma_{\mathrm{S}_{rr'}}^2\right]
\end{aligned}
$$

$$
\begin{aligned}
&+\sum_{r=1}^{R} \theta_{ij}^{(r)} \Gamma_{ij}^{(r)} \sigma_{\mathrm{G}_r}^2 \\
&= \sum_{r=1}^{R-1} \left( \Delta_{ij}^{(r)} + \frac{1}{2} \sum_{r' \neq r}^{R-1} \left( \Delta_{ij}^{(rr')} + \Delta_{ij}^{(r'r)} \right) \right) \sigma_{\mathrm{S}_{rR}}^2 - \frac{1}{2} \sum_{r=1}^{R-2} \sum_{r'=r+1}^{R-1} \left( \Delta_{ij}^{(rr')} + \Delta_{ij}^{(r'r)} \right) \sigma_{\mathrm{S}_{rr'}}^2 \\
&+ \sum_{r=1}^{R} \theta_{ij}^{(r)} \Gamma_{ij}^{(r)} \sigma_{\mathrm{G}_r}^2 \ .
\end{aligned}
$$

Since $\sum_{r=1} \Lambda_{imr}^{(h)} = 1$, we can find the identity

$$
\Delta_{ij}^{(r)} = \sum_{r', r'' \in \mathcal{R} \setminus \{r\}} \Delta_{ij}^{(r'r'')} \ , \tag{S9}
$$

which we can use to rewrite the coefficient in the sum over $\sigma_{\mathrm{S}_{rR}}^2$ (with a given $r$) as follows:

$$
\begin{aligned}
\Delta_{ij}^{(r)} + \frac{1}{2} \sum_{r' \neq r}^{R-1} \left( \Delta_{ij}^{(rr')} + \Delta_{ij}^{(r'r)} \right) &= \frac{1}{2} \left( 2\Delta_{ij}^{(r)} + \sum_{r' \neq r}^{R-1} \left( \Delta_{ij}^{(rr')} + \Delta_{ij}^{(r'r)} \right) \right) \\
&= \frac{1}{2} \left( \Delta_{ij}^{(r)} + \sum_{r', r'' \in \mathcal{R} \setminus \{R\}} \Delta_{ij}^{(r'r'')} - \sum_{r', r'' \in \mathcal{R} \setminus \{r, R\}} \Delta_{ij}^{(r'r'')} \right) \\
&\overset{eq. (S9)}{=} \frac{1}{2} \left( \Delta_{ij}^{(r)} + \Delta_{ij}^{(R)} - \sum_{r', r'' \in \mathcal{R} \setminus \{r, R\}} \Delta_{ij}^{(r'r'')} \right) \ .
\end{aligned}
$$

By again using eq. (S9) and the similar identities

$$
\Delta_{ij}^{(r)} = - \sum_{r' \neq r}^{R} \Delta_{ij}^{(rr')} \ , \tag{S10}
$$

$$
\Delta_{ij}^{(r)} = - \sum_{r' \neq r}^{R} \Delta_{ij}^{(r'r)} \ , \tag{S11}
$$

we can also rewrite the other coefficient (with a given $r$ and $r'$) as follows:

$$
\begin{aligned}
-\frac{1}{2} \left( \Delta_{ij}^{(rr')} + \Delta_{ij}^{(r'r)} \right) &= \frac{1}{2} \left( -\Delta_{ij}^{(rr')} - \sum_{r'' \neq r}^{R} \Delta_{ij}^{(r''r)} + \sum_{r'' \in \mathcal{R} \setminus \{r, r'\}} \Delta_{ij}^{(r''r)} \right) \\
&\overset{eq. (S11)}{=} \frac{1}{2} \left( -\Delta_{ij}^{(rr')} + \Delta_{ij}^{(r)} + \sum_{r'' \in \mathcal{R} \setminus \{r, r'\}} \Delta_{ij}^{(r''r)} \right) \\
&= \frac{1}{2} \left( - \sum_{r'' \neq r} \Delta_{ij}^{(rr'')} + \sum_{r'' \in \mathcal{R} \setminus \{r, r'\}} \Delta_{ij}^{(rr'')} + \Delta_{ij}^{(r)} + \sum_{r'' \in \mathcal{R} \setminus \{r, r'\}} \Delta_{ij}^{(r''r)} \right)
\end{aligned}
$$

$$= \frac{1}{2}\left( - \sum_{r'' \neq r} \Delta_{ij}^{(rr'')} + \sum_{r'',r^* \in \mathcal{R}\setminus\{r'\}} \Delta_{ij}^{(r''r^*)} - \sum_{r'',r^* \in \mathcal{R}\setminus\{r,r'\}} \Delta_{ij}^{(r''r^*)} \right)$$

$$\overset{eq.(\mathrm{S10}),\ eq.(\mathrm{S9})}{=} \frac{1}{2}\left( \Delta_{ij}^{(r)} + \Delta_{ij}^{(r')} - \sum_{r'',r^* \in \mathcal{R}\setminus\{r,r'\}} \Delta_{ij}^{(r''r^*)} \right) .$$

Inserting these rewritten coefficients, we can write

$$\begin{aligned}
&\mathsf{Cov}(U_i, U_j \mid \boldsymbol{\pi}_i, \boldsymbol{\pi}_j, \boldsymbol{\theta}_{ij}, \boldsymbol{\Gamma}_{ij}) \\
&= \frac{1}{2}\sum_{r=1}^{R-1} \left( \Delta_{ij}^{(r)} + \Delta_{ij}^{(R)} - \sum_{r',r'' \in \mathcal{R}\setminus\{r,R\}} \Delta_{ij}^{(r'r'')} \right) \sigma^2_{\mathrm{S}_{rR}} \\
&+ \frac{1}{2}\sum_{r=1}^{R-2}\sum_{r'=r+1}^{R-1} \left( \Delta_{ij}^{(r)} + \Delta_{ij}^{(r')} - \sum_{r'',r^* \in \mathcal{R}\setminus\{r,r'\}} \Delta_{ij}^{(r''r^*)} \right) \sigma^2_{\mathrm{S}_{rr'}} \\
&+ \sum_{r=1}^{R} \theta_{ij}^{(r)} \Gamma_{ij}^{(r)} \sigma^2_{\mathrm{G}_r} \\
&= \frac{1}{2}\sum_{r=1}^{R-1}\sum_{r'=r+1}^{R} \left( \Delta_{ij}^{(r)} + \Delta_{ij}^{(r')} - \sum_{r'',r^* \in \mathcal{R}\setminus\{r,r'\}} \Delta_{ij}^{(r''r^*)} \right) \sigma^2_{\mathrm{S}_{rr'}} \\
&+ \sum_{r=1}^{R} \theta_{ij}^{(r)} \Gamma_{ij}^{(r)} \sigma^2_{\mathrm{G}_r} \ .
\end{aligned}$$

Note that this formulation is useful in the case $R = 3$, as it then only involves $\Delta_{ij}^{(1)}, \Delta_{ij}^{(2)}$ and $\Delta_{ij}^{(3)}$, which are easier to compute than when we have $\Delta_{ij}^{(rr')}$ with $r \neq r'$. We can further rewrite the covariance structure:

$$\begin{aligned}
&\mathsf{Cov}(U_i, U_j \mid \boldsymbol{\pi}_i, \boldsymbol{\pi}_j, \boldsymbol{\theta}_{ij}, \boldsymbol{\Gamma}_{ij}) \\
&= \frac{1}{2}\sum_{r=1}^{R-1}\sum_{r'=r+1}^{R} \left( \Delta_{ij}^{(r)} + \Delta_{ij}^{(r')} - \sum_{r'',r^* \in \mathcal{R}\setminus\{r,r'\}} \Delta_{ij}^{(r''r^*)} \right) \sigma^2_{\mathrm{S}_{rr'}} \\
&+ \sum_{r=1}^{R} \theta_{ij}^{(r)} \Gamma_{ij}^{(r)} \sigma^2_{\mathrm{G}_r} \\
&= \frac{1}{2}\sum_{r=1}^{R-1}\sum_{r'=r+1}^{R} \left( \Delta_{ij}^{(r)} + \Delta_{ij}^{(r')} + \sum_{r^* \in \mathcal{R}\setminus\{r,r'\}} \left( \Delta_{ij}^{(rr^*)} + \Delta_{ij}^{(r'r^*)} \right) \right) \sigma^2_{\mathrm{S}_{rr'}} \\
&+ \sum_{r=1}^{R} \theta_{ij}^{(r)} \Gamma_{ij}^{(r)} \sigma^2_{\mathrm{G}_r} \\
&= \frac{1}{2}\sum_{r=1}^{R-1}\sum_{r'=r+1}^{R} \left( \Delta_{ij}^{(r)} + \sum_{r^* \in \mathcal{R}\setminus\{r,r'\}} \Delta_{ij}^{(rr^*)} + \Delta_{ij}^{(r')} + \sum_{r^* \in \mathcal{R}\setminus\{r,r'\}} \Delta_{ij}^{(r'r^*)} \right) \sigma^2_{\mathrm{S}_{rr'}}
\end{aligned}$$

$$
\begin{aligned}
&+ \sum_{r=1}^{R} \theta_{ij}^{(r)} \Gamma_{ij}^{(r)} \sigma^2_{\mathrm{G}_r} \\
&= \frac{1}{2} \sum_{r=1}^{R-1} \sum_{r'=r+1}^{R} \left( \sum_{r^* \in \mathcal{R} \setminus \{r'\}} \Delta_{ij}^{(rr^*)} + \sum_{r^* \in \mathcal{R} \setminus \{r\}} \Delta_{ij}^{(r'r^*)} \right) \sigma^2_{\mathrm{S}_{rr'}} \\
&+ \sum_{r=1}^{R} \theta_{ij}^{(r)} \Gamma_{ij}^{(r)} \sigma^2_{\mathrm{G}_r} \\
&= \frac{1}{2} \sum_{r=1}^{R-1} \sum_{r'=r+1}^{R} \left( -\Delta_{ij}^{(rr')} - \Delta_{ij}^{(r'r)} \right) \sigma^2_{\mathrm{S}_{rr'}} \\
&+ \sum_{r=1}^{R} \theta_{ij}^{(r)} \Gamma_{ij}^{(r)} \sigma^2_{\mathrm{G}_r} \\
&= -\frac{1}{2} \sum_{r=1}^{R-1} \sum_{r'=r+1}^{R} \left( \Delta_{ij}^{(rr')} + \Delta_{ij}^{(r'r)} \right) \sigma^2_{\mathrm{S}_{rr'}} + \sum_{r=1}^{R} \theta_{ij}^{(r)} \Gamma_{ij}^{(r)} \sigma^2_{\mathrm{G}_r} \ .
\end{aligned}
$$

This is a generalization of the result for the covariance between genetic values of different individuals found by Rio *et al.* (2020), and can be used to define the genetic covariance structure in a mixed model (the entries in a covariance matrix).

#### Usage and estimators

With the covariance structure for total genetic values in hand, we can see how to include random effects in a mixed model to implement this model. To find the group-specific additive genetic variance we include random effects with distribution $\mathsf{N}\left(\mathbf{0}, \sigma^2_{\mathrm{G}_r} \cdot \mathbf{G}_r\right)$ for each group $r$, where $\mathbf{G}_r$ is a GRM whose $ij^{\text{th}}$ entry estimates $\theta_{ij}^{(r)} \Gamma_{ij}^{(r)}$. Similarly, to include segregation variances in the model we include random effects with distribution $\mathsf{N}\left(\mathbf{0}, \sigma^2_{\mathrm{S}_{rr'}} \cdot \mathbf{S}_{rr'}\right)$ for each combination of groups $r$ and $r' \neq r$, where $\mathbf{S}_{rr'}$ is a matrix whose $ij^{\text{th}}$ entry estimates $-\frac{1}{2}\left(\Delta_{ij}^{(rr')} + \Delta_{ij}^{(r'r)}\right)$. We suggest estimators for $\theta_{ij}^{(r)}$ and $\Gamma_{ij}^{(r)}$ in the main text, while $\Delta_{ij}^{(rr')}$ can be rewritten as

$$
\Delta_{ij}^{(rr')} = \mathsf{Cov}\left(\Lambda_{imr}^{(h)},\ \Lambda_{jmr'}^{(h)}\right) = \mathsf{E}\left(\Lambda_{imr}^{(h)} \Lambda_{jmr'}^{(h)}\right) - \mathsf{E}\left(\Lambda_{imr}^{(h)}\right) \mathsf{E}\left(\Lambda_{jmr'}^{(h)}\right) = \theta_{ij}^{(rr')} - \pi_{ir}\pi_{jr'}.
$$

We suggest a definition for $\pi_{ir}$ in equation (5) in the main text, while $\theta_{ij}^{(rr')}$ can be estimated by

$$
\widehat{\theta}_{ij}^{(rr')} = \frac{1}{4M} \sum_{m=1}^{M} \sum_{h=1}^{2} \sum_{h'=1}^{2} \lambda_{imr}^{(h)} \lambda_{jmr'}^{(h')} \ .
$$

Thus, we can estimate the matrix $\mathbf{S}_{rr'}$ as follows:

$$
\mathbf{S}_{rr'} = -\frac{1}{2}\left(\widehat{\Delta}_{ij}^{(rr')} + \widehat{\Delta}_{ij}^{(r'r)}\right), \quad \text{where } \widehat{\Delta}_{ij}^{(rr')} = \widehat{\theta}_{ij}^{(rr')} - \widehat{\pi}_{ir}\widehat{\pi}_{jr'}. \tag{S12}
$$

#### Equivalence between vanRaden's GRM and our GRM in the one-group case

Note that in the case $R = 1$ we have $\lambda_{im1}^{(1)} = \lambda_{im1}^{(2)} = \lambda_{jm1}^{(1)} = \lambda_{jm1}^{(2)} = 1$. Inserting this into $\widehat{\theta}_{ij}^{(1)}$ (equation (7) in the main text) trivially yields $\widehat{\theta}_{ij}^{(1)} = 1$. Insert the same into the numerator of $\widehat{\Gamma}_{ij}^{(r)}$ (equation (8) in the main text) and rewrite:

$$
\begin{aligned}
&\sum_{m=1}^{M}\sum_{h=1}^{2}\sum_{h'=1}^{2} \lambda_{im1}^{(h)}\left(w_{im}^{(h)} - \widehat{p}_{m1}\right)\lambda_{jm1}^{(h')}\left(w_{jm}^{(h')} - \widehat{p}_{m1}\right)\\
&= \sum_{m=1}^{M}\sum_{h=1}^{2}\sum_{h'=1}^{2} \left(w_{im}^{(h)} - \widehat{p}_{m1}\right)\left(w_{jm}^{(h')} - \widehat{p}_{m1}\right)\\
&= \sum_{m=1}^{M}\Big[\left(w_{im}^{(1)} - \widehat{p}_{m1}\right)\left(w_{jm}^{(1)} - \widehat{p}_{m1}\right) + \left(w_{im}^{(1)} - \widehat{p}_{m1}\right)\left(w_{jm}^{(2)} - \widehat{p}_{m1}\right)\\
&\qquad + \left(w_{im}^{(2)} - \widehat{p}_{m1}\right)\left(w_{jm}^{(1)} - \widehat{p}_{m1}\right) + \left(w_{im}^{(2)} - \widehat{p}_{m1}\right)\left(w_{jm}^{(2)} - \widehat{p}_{m1}\right)\Big]\\
&= \sum_{m=1}^{M}\left(\left(w_{im}^{(1)} - \widehat{p}_{m1}\right) + \left(w_{im}^{(2)} - \widehat{p}_{m1}\right)\right)\left(\left(w_{jm}^{(1)} - \widehat{p}_{m1}\right) + \left(w_{jm}^{(2)} - \widehat{p}_{m1}\right)\right)\\
&= \sum_{m=1}^{M}\left(w_{im}^{(1)} + w_{im}^{(2)} - 2\widehat{p}_{m1}\right)\left(w_{jm}^{(1)} + w_{jm}^{(2)} - 2\widehat{p}_{m1}\right)\\
&= \sum_{m=1}^{M}\left(v_{im} - 2\widehat{p}_{m1}\right)\left(v_{jm} - 2\widehat{p}_{m1}\right)\\
&= \sum_{m=1}^{M}\left(v_{im} - 2\widehat{p}_{m}\right)\left(v_{jm} - 2\widehat{p}_{m}\right) \ ,
\end{aligned}
$$

since we have $\widehat{p}_{m1} = \widehat{p}_m$ when $R = 1$. As for the denominator of $\widehat{\Gamma}_{ij}^{(r)}$ (equation (8) in the main text), merely note that $\frac{1}{2}\left(\sum_{h=1}^{2}\sum_{h'=1}^{2}\lambda_{im1}^{(h)}\right) = \frac{1}{2}\left(\sum_{h=1}^{2}\sum_{h'=1}^{2} 1\right) = 2$. Thus,

$$
\begin{aligned}
(G_1)_{ij} = \widehat{\theta}_{ij}^{(1)}\cdot\widehat{\Gamma}_{ij}^{(1)} = \widehat{\Gamma}_{ij}^{(1)} &= \frac{\sum_{m=1}^{M}\sum_{h=1}^{2}\sum_{h'=1}^{2}\lambda_{im1}^{(h)}\left(w_{im}^{(h)} - \widehat{p}_{m1}\right)\lambda_{jm1}^{(h')}\left(w_{jm}^{(h')} - \widehat{p}_{m1}\right)}{\frac{1}{2}\sum_{m=1}^{M}\sum_{h=1}^{2}\sum_{h'=1}^{2}\lambda_{im1}^{(h)}\lambda_{jm1}^{(h')}\widehat{p}_{m1}\left(1 - \widehat{p}_{m1}\right)}\\
&= \frac{\sum_{m=1}^{M}\left(v_{im} - 2\widehat{p}_m\right)\left(v_{jm} - 2\widehat{p}_m\right)}{2\sum_{m=1}^{M}\widehat{p}_m\left(1 - \widehat{p}_m\right)} = G_{ij} \ .
\end{aligned}
$$

#### S2: Tutorial

Here we demonstrate how to generate group-specific GRMs starting from genotype data on a `.ped` and `.map` format, or a `.vcf` data format, and to fit Bayesian genetic group animal models using said GRMs.

##### Before you begin

Here we assume 3 genetic groups to be present in the study system. If your system has some other number of groups, and you can follow the code segments herein, it should not be hard to adapt the R code to a different number of groups. It should be noted that the parallelization in the R scripts is not implemented for Windows, and is therefore considerably slower on that OS. We recommend using Linux, or a Linux subsystem for Windows. You would further benefit from using a computational cluster, as some computations can be cumbersome depending on the size of your data set and the number of genetic groups. Furthermore, having plenty of free disc space is recommended, as the computations use some large temporary files. We also assume that the determination of which individuals are to be considered admixed or purebred in some genetic group has already been made by the user, as discussed in the main text.

##### Convert to .vcf format

We assume the genotype data is available on the `PLINK 1.9` (Chang *et al.*, 2015) genomic data file format `.ped` with an accompanying `.map` file. A `.ped` file contains counts of the alternate allele at each SNP for every individual, while a `.map` file contains additional information such as the chromosome each SNP is located on. More information on these files formats can be found at `https://www.cog-genomics.org/plink/1.9/formats`. The software we use to phase the genotype data, `Beagle`, takes `.vcf` files as input, so we use `PLINK` to convert the `.ped` and `.map` data to the `.vcf` format. Installation instructions for `PLINK` can be found at `https://www.cog-genomics.org/plink/1.9/`. Entering the `PLINK` command

```
plink --ped genotypes.ped --map genotypes.map --chr-set 32 --recode vcf-iid
    --out vcf_genotypes
```

to a command prompt outputs a file `vcf_genotypes.vcf` containing the genotype data on the `.vcf` format. The `chr-set` parameter refers to which chromosome set should be used, and might differ in your genotype data.

#### Gametic phasing

We used Beagle (Browning *et al.*, 2018) to impute missing genotypes and phase the genotype data so that we have data for single allele. We do this separately for each purebred population and for the admixed population. In each case we excluded the genotyped individuals ("samples") not belonging to our desired subpopulation, and other individuals one might want to filter out, by listing these individuals in .txt files. We similarly filtered the SNPs by call rates and alternate frequency, and left out SNPs not assigned to a specific chromosome. The following Beagle commands were used to phased/impute the data. See https://faculty.washington.edu/browning/beagle/beagle.html for download and further instructions on Beagle. We phase/impute the genotype data in the inner genetic group with the following (command line) command:

```
java -jar beagle.28Jun21.220.jar gt=vcf_genotypes.vcf
    excludemarkers=markers_to_exclude.txt
    excludesamples=NOT_inner_inds.txt out=inner_phased
```

And similarly for outer:

```
java -jar beagle.28Jun21.220.jar gt=vcf_genotypes.vcf
    excludemarkers=markers_to_exclude.txt
    excludesamples=NOT_outer_inds.txt out=outer_phased
```

other:

```
java -jar beagle.28Jun21.220.jar gt=vcf_genotypes.vcf
    excludemarkers=markers_to_exclude.txt
    excludesamples=NOT_other_inds.txt out=other_phased
```

and the admixed individuals:

```
java -jar beagle.28Jun21.220.jar gt=vcf_genotypes.vcf
    excludemarkers=markers_to_exclude.txt
    excludesamples=NOT_admixed_inds.txt out=admixed_phased
```

The output is given as compressed variant call format files .vcf.gz.

#### Local ancestry inference

We now have phased genomic data available on the .vcf format, separately for each of the three reference populations and the admixed population. We can run local ancestry inference on the admixed population using, for example, the command-line version of `Loter` (Dias-Alves *et al.*, 2018). Other local ancestry inference tools are available and might fit your needs better (Padhukasahasram, 2014; Geza *et al.*, 2019; Schubert *et al.*, 2020; Utsunomiya *et al.*, 2020). For installation instructions, see `https://github.com/bcm-uga/Loter`. We ran the local ancestry inference with the following command:

```
loter_cli -f 'vcf'
    -r inner_phased.vcf.gz outer_phased.vcf.gz other_phased.vcf.gz
    -a admixed_phased.vcf.gz -n 8 -o loter_out.txt -v
```

The use of the `-f` option makes `Loter` accept the phased genotype data on the `.vcf` format, while the `-r` option lists the reference populations, the `-a` option lists the admixed population, the `-n` option lists the number of cores to be used, `-o` names the output file and `-v` tells `Loter` to use the verbose option, that is, it outputs more information while running. Depending on your computational setup and the size of your data set, this might take several hours to days. The local ancestry output is written to `loter_out.txt` where entries 0, 1 and 2 indicate allele group ancestry in `inner`, `outer` and `other`, respectively (corresponding to the order they are listed in the above command). This file can take up a lot of storage depending on the size of your data set (it required 1.86GB in our data set with 3032 individuals and 181354 SNPs). The rows are organized by individual and chromosome copy, so that row 1 contains individual 1 and copy $h = 1$, row 2 contains individual 1 copy $h = 2$, row 3 contains individual 2 copy $h = 1$, row 4 contains individual 2 copy $h = 2$, *etc.* The columns are organized by locus, so that column $m$ contains alleles on locus $m$.

#### File-backed matrices

We perform the remaining analysis in R. Note that R accumulates memory usage in over longer sessions. We therefore regularly save partial results, so that the R session can be reloaded in the case it needs to be restarted to flush memory. To work with the very large genomic data sets directly in R we use the package `BGData` (Grueneberg & de los Campos, 2019). `BGData` objects have a slot for phenotype data and a slot for genotype data, the latter being where the large genomic data sets are stored in file backed matrices. The package has a function `readRAW()` to read data from dosage files, but since we will be reading data on the `.vcf` format, we create the utility function `initFileBackedMatrix()` which lets us create custom file backed matrices. The function `initFileBackedMatrix()` utilizes the function `ffNodeInitializer()`, which we have taken from the source code of `BGData` – it is not necessary to understand the details of `ffNodeInitializer()`. Importantly, we can specify the data type contained in the matrix, which will determine how much disk space is taken up.

```
library(Matrix)
library(BGData)
library(ff)
library(cgwtools)
# Utility function to initialize file backed system
ffNodeInitializer = function(nodeIndex, nrow, ncol, vmode, folderOut, ...) {
    filename <- paste0("geno_", nodeIndex, ".bin")
    node <- ff(dim = c(nrow, ncol), vmode = vmode,
               filename = paste0(folderOut, "/", filename), ...)
    # Change ff path to a relative one
    physical(node)[["filename"]] <- filename
    return(node)
}
# Function that initializes a file-backed matrix of a specific size, and data
# type (outputType). Data types: "boolean" = TRUE/FALSE without NA,
# "logical" = TRUE/FALSE (with NA), "byte" (integers up to 2^7), "double", etc.
initFileBackedMatrix = function(nrows, ncols, folderOut, outputType) {
    dir.create(folderOut)
    chunkSize = min(nrows, floor(.Machine[["integer.max"]] / ncols / 1.2))
    nNodes = ceiling(nrows / chunkSize)
    matrix = LinkedMatrix(nrow = nrows, ncol = ncols, nNodes = nNodes,
                          linkedBy = "rows", vmode = outputType,
                          folderOut = folderOut, dimorder = 2:1,
                          nodeInitializer = ffNodeInitializer)
    return(matrix)
}
```

#### Allele variant matrix

We load information such as the individual ID in each of the subpopulations (`inner`, `outer`, `other` and admixed individuals), the population sizes, SNP names, *etc.* into R. Note that the `getPop()` function might need to skip a different number of lines than 8, depending on the number of lines of metadata the `.vcf` file contains.

```
library(R.utils)
library(data.table)
# Function that fetches the ring numbers identifying individuals in each
# subpopulation from the .vcf files with phased genotype data
getPop = function(group) {
    groupFile = file(paste0(group, "_phased.vcf.gz"), open = "r")
    groupLine = scan(groupFile, nlines = 1, skip = 8, what = character(),
                     quiet = TRUE)
    close(groupFile)
    return(groupLine[-(1:9)])
}
# Population is grouped by inner, outer, other, admixed
innerInds = getPop("inner")
outerInds = getPop("outer")
otherInds = getPop("other")
admixedInds = getPop("admixed")
pop = c(innerInds, outerInds, otherInds, admixedInds) # full population
# Number of individuals in each (sub)population
numInnerInds = length(innerInds)
numOuterInds = length(outerInds)
numOtherInds = length(otherInds)
numAdmixedInds = length(admixedInds)
N = length(pop)
# Fetch the names of each SNP (from smallest group to load quicker)
SNPs = fread(file = "other_phased.vcf.gz", select = 3)$ID
numSNPs = length(SNPs) # Nnumber of SNPs
save.image(file = "setup.RData")
```

Now the imputed/phased allele data located in the four `.vcf.gz` files is read and are merged into a single file-backed matrix `W` on the `BGData` format. The entries of this matrix will contain indicators $w_{im}^{(h)}$ for the alternate allele, as defined in the main text, where $w_{im}^{(h)} = 1$ if the alternate allele is present at individual $i$'s $m^{\text{th}}$ locus on chromosome copy $h$ (and $w_{im}^{(h)} = 0$ otherwise). We let the first $N(= 3032)$ rows of `W` correspond to each individual's $h = 1$ allele, and the next $N$ rows correspond to each individual's $h = 2$ allele. Thus, the rows are sorted hierarchically by chromosome copy $h$ and individual $i$. Column $m$ corresponds to locus $m$. In other words, the entries in the first row of `W` are $w_{11}^{(1)}$, $w_{12}^{(1)}$, $w_{13}^{(1)}$, etc, while the entries in the $(N+1)^{\text{th}}$ row of `W` are $w_{11}^{(2)}$, $w_{12}^{(2)}$, $w_{13}^{(2)}$, *etc*. This grouping is useful for when we will compare all combinations of $h = 1$ and $h = 2$. The individuals are ordered so that all purebred `inner` individuals come first, followed by all purebred `outer` individuals, all purebred `other` individuals and finally all admixed individuals. The four `.vcf` files contain the genotype data on the format $x|y$, where $x$ is an indicator of the $h = 1$ allele and $y$ is an indicator of the $h = 2$ allele. Thus, $x = 1$ if the alternate allele is present on the $h = 1$ chromosome copy and 0 otherwise, and $y = 1$ if the alternate allele is present on the $h = 2$ chromosome copy and 0 otherwise. We use regular expressions to extract this information.

```
# Create file-backed matrix to hold alternate allele indicators w_{im}^{(h)}.
# First N rows correspond to chromosome copy 1 and next N correspond to chromosome copy 2
# When initializing a BGData object, the `pheno` slot holds phenotypes, and
# the `geno` slot holds genotypes.
W = BGData(pheno = data.frame(ind = rep(pop, 2), h = rep(c(1, 2), each = N)),
           geno = initFileBackedMatrix(nrows = 2 * N, ncols = numSNPs,
                                       outputType = "boolean", folderOut = "W"))
# Function to fill matrix with allele variant data from the .vcf format
fillW = function(group, firstInd, lastInd) {
    # Load group data into groupVCF
    groupVCF = fread(file = paste0(group, "_phased.vcf.gz"), drop = 1:9)
    # Check if alleles have alternate allele present using regular expressions
    h1 = lapply(groupVCF, function(x) grepl("^1", x)) # Loci with w=1 on h=1
    h2 = lapply(groupVCF, function(x) grepl("1$", x)) # Loci with w=1 on h=2
    rm(groupVCF)
    # Save alternate allele indicators in BGData object
    for (ind in firstInd:lastInd) {
```

```
        geno(W)[ind, ] = h1[[ind + 1 - firstInd]]
        geno(W)[ind + N, ] = h2[[ind + 1 - firstInd]]
    }
    rm(h1, h2)
}
# Run function on each subpopulation:
# Inner
fillW("inner", 1, numInnerInds)
# Outer
fillW("outer", numInnerInds + 1, numInnerInds + numOuterInds)
# Other
fillW("other", numInnerInds + numOuterInds + 1, N - numAdmixedInds)
# Admixed
fillW("admixed", numInnerInds + numOuterInds + numOtherInds + 1, N)
# Give the BGData object reasonable row and column names in each slot
rownames(map(W)) = colnames(geno(W)) = SNPs
rownames(pheno(W)) = apply(pheno(W), 1, paste, collapse = "_")
# Save the BGData object so we don't have to rerun this to access W
save(W, file = "W/W.RData")
```

#### Local ancestry matrix

We convert the local ancestry data output from Loter in the file `loter_out.txt` to a local ancestry matrix `A`, again a `BGData` object. The entries in `A` correspond to the local ancestry indicators $\lambda_{imr}^{(h)}$ from the main text, and equal 1 when individual $i$'s allele at locus $m$, chromosome copy $h$, is descended from genetic group $r$ (and equal 0 otherwise). Rows are sorted hierarchically by genetic group $r$, chromosome copy $h$ and individual $i$. Column $m$ corresponds to locus $m$. Thus, the first row has entries $\lambda_{111}^{(1)}$, $\lambda_{121}^{(1)}$, $\lambda_{131}^{(1)}$, *etc.*, the second row has entries $\lambda_{211}^{(1)}$, $\lambda_{221}^{(1)}$, $\lambda_{231}^{(1)}$, *etc.*, the $(N+1)^{\text{th}}$ row has entries $\lambda_{111}^{(2)}$, $\lambda_{121}^{(2)}$, $\lambda_{131}^{(2)}$, *etc.*, the $(N+2)^{\text{th}}$ row has entries $\lambda_{211}^{(2)}$, $\lambda_{221}^{(2)}$, $\lambda_{231}^{(2)}$, *etc.* and the $(2N+1)^{\text{th}}$ row has entries $\lambda_{112}^{(1)}$, $\lambda_{122}^{(1)}$, $\lambda_{132}^{(1)}$, *etc.*

```
# Create a BGData object for the local ancestry matrix.
# Row order: inner h=1, inner h=2, outer h=1, outer h=2, other h=1, other h=2
A = BGData(geno = initFileBackedMatrix(3 * 2 * N, numSNPs, outputType="boolean",
```

```
                                        folderOut = "A"),
            pheno = data.frame(ind = rep(pop, 3 * 2),
                                   h = rep(rep(c(1, 2), each = N), 3)))
# Give reasonable row and column names
dimnames(map(A))= dimnames(map(W))
dimnames(geno(A))[[2]] = dimnames(geno(W))[[2]]
dimnames(pheno(A)) =
    list(paste0(rep(c("Inner_", "Outer_", "Other_"), each = N * 2),
                dimnames(pheno(W))[[1]]),
         dimnames(pheno(W))[[2]])
# Loop through the "inner" purebreds and set local ancestry indicator to
# 1 (TRUE) for inner group, and to 0 (FALSE) for "outer" and "other", for
# all alleles (both chromosomes) in these individuals.
for (ind in 1:numInnerInds) {
    geno(A)[ind, ] = geno(A)[ind + N, ] = rep(TRUE, numSNPs)
    geno(A)[ind + (2 * N), ] =
        geno(A)[ind + (3 * N), ] =
        geno(A)[ind + (4 * N), ] =
        geno(A)[ind + (5 * N), ] =
        rep(FALSE, numSNPs)
}
# Loop through the "outer" purebreds and set local ancestry indicator to
# 1 (TRUE) for inner group, and to 0 (FALSE) for "inner" and "other", for
# all alleles (both chromosomes) in these individuals.
for (ind in (numInnerInds + 1):(numInnerInds + numOuterInds)) {
  geno(A)[ind + (2 * N), ] = geno(A)[ind + (3 * N), ] = rep(TRUE, numSNPs)
  geno(A)[ind , ] =
    geno(A)[ind + N, ] =
    geno(A)[ind + (4 * N), ] =
    geno(A)[ind + (5 * N), ] =
    rep(FALSE, numSNPs)
}
# Loop through the "other" purebreds and set local ancestry indicator to
```

```
# 1 (TRUE) for inner group, and to 0 (FALSE) for "inner" and "outer", for
# all alleles (both chromosomes) in these individuals.
for (ind in (numInnerInds + numOuterInds + 1):(N - numAdmixedInds)) {
  geno(A)[ind + (4 * N), ] = geno(A)[ind + (5 * N), ] = rep(TRUE, numSNPs)
  geno(A)[ind , ] =
    geno(A)[ind + N, ] =
    geno(A)[ind + (2 * N), ] =
    geno(A)[ind + (3 * N), ] =
    rep(FALSE, numSNPs)
}
# For the admixed individuals we use the loter results to determine values
# of the local ancestry indicators. In the loter output text file
# 0 indicates "inner", 1 indicates "outer", 2 indicates "other"
admixedFile = file("loter_out.txt", open = "r")
for (ind in (N - numAdmixedInds + 1):N) {
    # chromosome 1:
    line1 = scan(admixedFile, nlines = 1, what = integer(), quiet = TRUE)
    geno(A)[ind, ] = (line1 == 0)
    geno(A)[ind + (2 * N), ] = (line1 == 1)
    geno(A)[ind + (4 * N), ] = (line1 == 2)
    # chromosome 2:
    line2 = scan(admixedFile, nlines = 1, what = integer(), quiet = TRUE)
    geno(A)[ind + N, ] = (line2 == 0)
    geno(A)[ind + (3 * N), ] = (line2 == 1)
    geno(A)[ind + (5 * N), ] = (line2 == 2)
}
close(admixedFile)
# Save converted data in local ancestry matrix with 0/1 entries
save(A, file = "A/A.RData")
```

#### Group-specific allele frequencies

Before moving on we compute some auxillary information regarding how the parallel implementation of BGData, such as the number of cores to be used and how the tasks shoueld be split up in parallel.

```
library(parallel)
# The parallel implementation in BGData does not work on Windows, so if ran on
# Windows, use only one core (slow), otherwise use desired amount. Here we use
# all cores with detectCores() function. If you are using the computer you are
# running this on for something else, consider using a lower number of cores.
numCores = ifelse(Sys.info()[['sysname']] == "Windows", 1, detectCores())
# Find the optimal number of "chunks" each core should compute (=passes) and how
#large the chunks should be (=chunksize).
passes = round(numSNPs / 5000 / numCores)
passes = ifelse(passes == 0, 1, passes)
colChunkSize = ceiling(numSNPs / (passes * numCores))
rowChunkSize = ceiling(N * 6 / (passes * numCores))
save(numCores, colChunkSize, rowChunkSize, file = "parallel_aux_info.RData")
```

We compute group-specific allele frequency vectors $\boldsymbol{p}_r$ for each group $r \in \{1,2,3\}$ with entries $p_{1r}, p_{2r}, p_{3r}$, *etc.* We use the estimator defined in the main text, namely

$$\widehat{p}_{mr} = \frac{\sum_{i=1}^{N}\sum_{h=1}^{2}\lambda_{imr}^{(h)} w_{im}^{(h)}}{\sum_{i=1}^{N}\sum_{h=1}^{2}\lambda_{imr}^{(h)}} .$$

We compute the numerator and denominator vectors separately, and divide entry-wise to find $\widehat{p}_{mr}$. As a step of computing the numerator, we create the file-backed matrices `WAInner`, `WAOuter`, and `WAOther`, which contain the entry-wise product of `W` and the entries of `A` relating to `inner`, `outer` and `other`, respectively.

```
# We create a temporary matrix WA to efficiently compute the numerator of
# the estimator for the allele frequencies p. Each matrix holds the product of
# W and A for a given group.
WAInner = BGData(pheno = pheno(W),
                 geno = initFileBackedMatrix(2 * N, numSNPs,
                                             outputType = "boolean",
                                             folderOut = "WAInner"))
```

```
WAOuter = BGData(pheno = pheno(W),
                 geno = initFileBackedMatrix(2 * N, numSNPs,
                                             outputType = "boolean",
                                             folderOut = "WAOuter"))
WAOther = BGData(pheno = pheno(W),
                 geno = initFileBackedMatrix(2 * N, numSNPs,
                                             outputType = "boolean",
                                             folderOut = "WAOther"))
# Fill WA matrices, implemented in parallel
WA_func = function(row) {
    geno(WAInner)[row, ] = geno(W)[row, ] & geno(A)[row, ]
    geno(WAOuter)[row, ] = geno(W)[row, ] & geno(A)[row + (2 * N), ]
    geno(WAOther)[row, ] = geno(W)[row, ] & geno(A)[row + (4 * N), ]
    return(NULL)
}
trash = mclapply(1:(2 * N), WA_func, mc.cores = numCores)
rm(trash)
save(WAInner, file = "WAInner/WAInner.RData")
save(WAOuter, file = "WAOuter/WAOuter.RData")
save(WAOther, file = "WAOther/WAOther.RData")
# Compute vector of numerator of estimator of p for each group
pNumerInner = chunkedApply(geno(WAInner), 2, sum, nCores = numCores, v = TRUE,
                           chunkSize = colChunkSize)
pNumerOuter = chunkedApply(geno(WAOuter), 2, sum, nCores = numCores, v = TRUE,
                           chunkSize = colChunkSize)
pNumerOther = chunkedApply(geno(WAOther), 2, sum, nCores = numCores, v = TRUE,
                           chunkSize = colChunkSize)
# Compute vector of denominator of estimator of p for each group
pDenomInner = chunkedApply(geno(A), 2, sum, nCores = numCores, v = TRUE,
                           i = 1:(2 * N), chunkSize = colChunkSize)
pDenomOuter = chunkedApply(geno(A), 2, sum, nCores = numCores, v = TRUE,
                           i = (2 * N + 1):(4 * N), chunkSize = colChunkSize)
pDenomOther = chunkedApply(geno(A), 2, sum, nCores = numCores, v = TRUE,
```

```r
                              i = (4 * N + 1):(6 * N), chunkSize = colChunkSize)
# Group-specific allele frequency vector for each group
pInner = pNumerInner / pDenomInner
pOuter = pNumerOuter / pDenomOuter
pOther = pNumerOther / pDenomOther
rm(pNumerInner, pNumerOuter, pNumerOther, pDenomInner, pDenomOuter, pDenomOther)
save(pInner, pOuter, pOther, file = "alleleFreqs.RData")
```

#### Global ancestry proportions

We estimate the global ancestry proportions using the estimator

$$\widehat{\pi}_{ir} = \frac{1}{2M} \sum_{m=1}^{M} \sum_{h=1}^{2} \lambda_{imr}^{(h)} \ .$$

```r
# Mean of each row in A
piTemp = chunkedApply(geno(A), MARGIN = 1, mean, nCores = numCores, v = TRUE,
                      chunkSize = rowChunkSize)
# Take the mean of piTemp entries that refer to same locus and same group
piFull = unname(tapply(X = piTemp, FUN = mean,
                       INDEX = c(rep(1:(N), 2),
                                 rep((N + 1):(N * 2), 2),
                                 rep((N * 2 + 1):(N * 3), 2))))
rm(piTemp)
piInner = piFull[1:N]
piOuter = piFull[(N + 1):(N * 2)]
piOther = piFull[(N * 2 + 1):(N * 3)]
rm(piFull)
names(piInner) = names(piOuter) = names(piOther) = pop
# Save global ancestries
save(piInner, piOuter, piOther, file = "pi.RData")
```

##### Ancestry overlaps

We compute the matrix $\widehat{\boldsymbol{\theta}}^{(r)}$ which has $ij^{\text{th}}$ entry

$$\widehat{\theta}_{ij}^{(r)} = \frac{1}{4M} \sum_{m=1}^{M} \sum_{h=1}^{2} \sum_{h'=1}^{2} \lambda_{imr}^{(h)} \lambda_{jmr}^{(h')}$$

for a given group $r$. This involves summing four matrix products, one for each permutation of $h \in \{1, 2\}$ and $h' \in \{1, 2\}$. We use the `BGData` function `getG()` to parallelize these matrix products.

```
# Function that finds the theta matrix for a given group
find_theta = function(group) {
    groupNum = switch(group, Inner = 0, Outer = 2, Other = 4)
    # Comparison of h=1 in individual i and h=1 in individual j
    theta_11 = getG(geno(A), center = FALSE, scale = FALSE, scaleG = FALSE,
                    nCores = numCores, verbose = TRUE, chunkSize = colChunkSize,
                    i = (1 + N * groupNum):(N * (groupNum + 1)))
    # Comparison of h=2 in individual i and h=2 in individual j
    theta_22 = getG(geno(A), center = FALSE, scale = FALSE, scaleG = FALSE,
                    nCores = numCores, verbose = TRUE, chunkSize = colChunkSize,
                    i = (1 + N * (groupNum + 1)):(N * (groupNum + 2)))
    # Cross-comparison: h=1 in individual i and h=2 in individual j
    theta_12 = getG(geno(A), center = FALSE, scale = FALSE, scaleG = FALSE,
                    nCores = numCores, verbose = TRUE, chunkSize = colChunkSize,
                    i = (1 + N * (groupNum)):(N * (groupNum + 1)),
                    i2 = (1 + N * (groupNum + 1)):(N * (groupNum + 2)))
    # The second cross comparison is simply the transpose of the first.
    theta_21 = t(theta_12)
    # Sum and average over all four matrix products to get theta
    theta = (theta_11 + theta_12 + theta_21 + theta_22) / (4 * numSNPs)
    dimnames(theta) = list(pop, pop)
    return(theta)
}
# Find theta matrix for inner group
thetaInner = find_theta("Inner")
```

```
# Save inner theta matrix
save(thetaInner, file = "theta.RData")
# Find theta matrix for outer group
thetaOuter = find_theta("Outer")
# Save outer theta matrix - resave() adds the matrix to an existing .RData object
resave(thetaOuter, file = "theta.RData")
# Find theta matrix for other group
thetaOther = find_theta("Other")
# Save other theta matrix
resave(thetaOther, file = "theta.RData")
```

#### Construction of GRMs

We compute the matrix $\widehat{\mathbf{\Gamma}}^{(r)}$ with entries

$$\widehat{\Gamma}_{ij}^{(r)} = \frac{\sum_{m=1}^{M}\sum_{h=1}^{2}\sum_{h'=1}^{2}\lambda_{imr}^{(h)}\left(w_{im}^{(h)} - \widehat{p}_{mr}\right)\lambda_{jmr}^{(h')}\left(w_{jm}^{(h')} - \widehat{p}_{mr}\right)}{\frac{1}{2}\sum_{m=1}^{M}\sum_{h=1}^{2}\sum_{h'=1}^{2}\lambda_{imr}^{(h)}\lambda_{jmr}^{(h')}\widehat{p}_{mr}\left(1 - \widehat{p}_{mr}\right)},$$

by calculating the numerator and denominator of the estimator separately and then divide entry-wise. Both the numerator and denominator involves a sum of four matrix products. The computations for the matrix making up the numerator of $\widehat{\mathbf{\Gamma}}^{(r)}$ rely on a large temporary file-backed matrix which we denote $\mathbf{V}_r$. This matrix will require a lot of storage (in this case 8.42GB) due to its data type (double precision floats), so the user must have sufficient memory storage before use.

```
findGammaNumerator = function(group) {
    groupNum = switch(group, Inner = 0, Outer = 2, Other = 4)
    p = get(paste0("p", group))
    # Create temporary V matrix
    V = BGData(pheno = pheno(A)[(1 + groupNum * N):(((2 + groupNum) * N)), ],
               geno = initFileBackedMatrix(2 * N, numSNPs,
                                           folderOut = paste0("V", group),
                                           outputType = "double"))
    # Function to fill a given row of V
    V_func = function(row) {
        geno(V)[row, ] = geno(get(paste0("WA", group)))[row, ] -
```

```r
                         geno(A)[row + (N * groupNum), ] * p
        return(NULL)
    }
    # Fill the V matrix
    trash = mclapply(1:(2 * N), V_func, mc.cores = numCores)
    rm(trash)
    # Compute four matrix products: all permutations of h=1 and h=2
    gammaNumer11 = getG(geno(V), i = 1:N,
                        center = FALSE, scale = FALSE, scaleG = FALSE,
                        nCores = numCores, v = TRUE, chunkSize = colChunkSize)
    gammaNumer22 = getG(geno(V), i = (N + 1):(2 * N),
                        center = FALSE, scale = FALSE, scaleG = FALSE,
                        nCores = numCores, v = TRUE, chunkSize = colChunkSize)
    gammaNumer12 = getG(geno(V), i = 1:N, i2 = (N + 1):(2 * N),
                        center = FALSE, scale = FALSE, scaleG = FALSE,
                        nCores = numCores, v = TRUE, chunkSize = colChunkSize)
    gammaNumer21 = t(gammaNumer12)
    # Delete the V matrix (there is no further need for it)
    unlink(paste0("V", group, "/geno_1.bin"), recursive = TRUE)
    unlink(paste0("V", group), recursive = TRUE)
    # Sum the four matrix products
    gammaNumer = gammaNumer11 + gammaNumer12 + gammaNumer21 + gammaNumer22
    dimnames(gammaNumer) = list(pop, pop)
    return(gammaNumer)
}
# Find numerator matrix of gamma in inner group
gammaNumerInner = findGammaNumerator("Inner")
save(gammaNumerInner, file = "gammaNumerator.RData")
# Find numerator matrix of gamma in outer group
gammaNumerOuter = findGammaNumerator("Outer")
resave(gammaNumerOuter, file = "gammaNumerator.RData")
# Find numerator matrix of gamma in other group
gammaNumerOther = findGammaNumerator("Other")
```

```r
resave(gammaNumerOther, file = "gammaNumerator.RData")
```

Similarly, the denominator matrix of $\widehat{\mathbf{\Gamma}}^{(r)}$ is computed as a sum of four matrix products.

```r
findGammaDenominator = function(group){
    groupNum = switch(group, Inner = 0, Outer = 2, Other = 4)
    p = get(paste0("p", group))
    pScaling = 1 / sqrt(p * (1 - p))
    gammaDen11 = getG(geno(A), i = (1 + N * groupNum):((groupNum + 1) * N),
                      center = FALSE, scale = pScaling, scaleG = FALSE,
                      nCores = numCores, verbose = TRUE, chunkSize = colChunkSize)
    gammaDen22 = getG(geno(A), i = ((groupNum + 1) * N + 1):((groupNum + 2) * N),
                      center = FALSE, scale = pScaling, scaleG = FALSE,
                      nCores = numCores, verbose = TRUE, chunkSize = colChunkSize)
    gammaDen12 = getG(geno(A), i = (1 + N * groupNum):((groupNum + 1) * N),
                      i2 = ((groupNum + 1) * N + 1):((groupNum + 2) * N),
                      center = FALSE, scale = pScaling, scaleG = FALSE,
                      nCores = numCores, verbose = TRUE, chunkSize = colChunkSize)
    gammaDen21 = t(gammaDen12)
    # Sum the four matrix products and don't forget the 1/2 from the definition
    gammaDen = (gammaDen11 + gammaDen12 + gammaDen21 + gammaDen22) / 2
    # Set zeros to very small values, to avoid dividing by zero later
    gammaDen = ifelse(gammaDen == 0, 1e-12, gammaDen)
}
# Find denominator matrix of gamma in inner group
gammaDenInner = findGammaDenominator("Inner")
save(gammaDenInner, file = "gammaDenominator.RData")
# Find denominator matrix of gamma in outer group
gammaDenOuter = findGammaDenominator("Outer")
resave(gammaDenOuter, file = "gammaDenominator.RData")
# Find denominator matrix of gamma in other group
gammaDenOther = findGammaDenominator("Other")
resave(gammaDenOther, file = "gammaDenominator.RData")
```

We finally find the group-specific GRMs $\mathbf{G}_r$ for a given group $r$ by putting all the pieces together:

divide the numerator matrix entry-wise by the denominator matrix, and also multiply entry-wise by $\widehat{\boldsymbol{\theta}}^{(r)}$.

```
# Find final GRMs by entry-wise division of numerator and denominator of gamma,
# and entry-wise multiplication with theta for each group.
innerGRM = gammaNumerInner / gammaDenInner * thetaInner
outerGRM = gammaNumerOuter / gammaDenOuter * thetaOuter
otherGRM = gammaNumerOther / gammaDenOther * thetaOther
dimnames(innerGRM) = dimnames(outerGRM) = dimnames(otherGRM) = list(pop, pop)
save(innerGRM, outerGRM, otherGRM, file = "GRMs.RData")
```

#### INLA model

We use the group-specific GRMs to fit genomic genetic group animal models using the package `R-INLA` (Rue *et al.*, 2009). Operating in a Bayesian framework, `INLA` produces posterior distributions for our model parameter. In terms of priors, we used $\mathsf{N}\left(\mathbf{0}, 10^3\mathbf{I}\right)$ normal distributions for the fixed effects, while all variances were given penalized complexity priors PC$(1, 0.05)$ (Simpson *et al.*, 2017). The version of the `R-INLA` released March 2020 was used, as we find this version to be the most stable. We fit three different models here, with the responses being the phenotypes wing length, body mass and tarsus length, respectively. The modeling choices are described more thoroughly in the main text.

```
library(fs)
library(INLA)
# Load phenotype data in the data frame `phenotypes`. The data frame contains
# wing length, body mass, tarsus length, a ring number identifying each
# sparrow (`ringnr`) and a corresponding numerical ID.
load(file = "phenotypes.RData")
# We add global ancestries to the phenotype data frame.
phenotypes$innerPi = piInner[match(phenotypes$ringnr, colnames(innerGRM))]
phenotypes$outerPi = piOuter[match(phenotypes$ringnr, colnames(outerGRM))]
phenotypes$otherPi = piOther[match(phenotypes$ringnr, colnames(otherGRM))]
# We shrink the relatedness matrices to only retain relatednesses for
# individuals for which we have phenotype data. Individuals that were not
# phenotyped are no longer useful.
```

```r
innerGRM = innerGRM[colnames(innerGRM) %in% phenotypes$ringnr,
                    colnames(innerGRM) %in% phenotypes$ringnr]
outerGRM = outerGRM[colnames(outerGRM) %in% phenotypes$ringnr,
                    colnames(outerGRM) %in% phenotypes$ringnr]
otherGRM = otherGRM[colnames(otherGRM) %in% phenotypes$ringnr,
                    colnames(otherGRM) %in% phenotypes$ringnr]
N = dim(innerGRM)[1]
# Numeric IDs for use in INLA (multiple copies required for technical reasons)
# ID1 will be used to fit an identity effect, ID2 to fit inner-specific VA,
# ID3 to fit outer-specific VA and ID4 to fit other-specific VA.
phenotypes$ID4 = phenotypes$ID3 = phenotypes$ID2 = phenotypes$ID1 = phenotypes$ID
# Rename dimnames to match the numeric IDs
dimnames(innerGRM)[[1]] = dimnames(innerGRM)[[2]] =
    phenotypes[match(colnames(innerGRM), phenotypes$ringnr), "ID"]
dimnames(outerGRM)[[1]] = dimnames(outerGRM)[[2]] =
    phenotypes[match(colnames(outerGRM), phenotypes$ringnr), "ID"]
dimnames(otherGRM)[[1]] = dimnames(otherGRM)[[2]] =
    phenotypes[match(colnames(otherGRM), phenotypes$ringnr), "ID"]
# Set group-specific IDs to NA if an individual does not belong to a group.
# This is done for INLA-implementation reasons.
phenotypes$ID2 = ifelse(phenotypes$innerPi > 0, phenotypes$ID2, NA)
phenotypes$ID3 = ifelse(phenotypes$outerPi > 0, phenotypes$ID3, NA)
phenotypes$ID4 = ifelse(phenotypes$otherPi > 0, phenotypes$ID4, NA)
# Ensure GRMs are positive-definite by adding a tiny value to the diagonal
innerGRM = innerGRM + diag(1e-12, N, N)
outerGRM = outerGRM + diag(1e-12, N, N)
otherGRM = otherGRM + diag(1e-12, N, N)
```

Note that the tiny value added to the diagonal might need to be larger depending on your data. An alternative to this method is to use the function `make.positive.definite()` from the package `corpcor` package, which implements a technique (Higham, 1988) for finding the closest positive definite matrix.

```r
# Check if matrices are positive definite
```

```r
stopifnot(all(eigen(innerGRM, only.values = TRUE)$values > 0))
stopifnot(all(eigen(outerGRM, only.values = TRUE)$values > 0))
stopifnot(all(eigen(otherGRM, only.values = TRUE)$values > 0))
# We now define the INLA formula. First, fixed effects to be used in the model:
effects = c("sex", "FGRM", "month", "age", "outerPi", "otherPi")
# Random effects to be used in the model are hatch year, island of observation,
# ID (named `ID1`), and each group-specific genetic value (`ID2, `ID3`, `ID4`).
# The former three are implemented with IID models, and the latter three with
# generic0 models. See INLA documentation for more information on how to fit
# mixed models with other random effect distributions.
# Here we also specify the prior distributions for the random effect variances.
randomEffects =
    c("f(hatchYear, model = \"iid\",
         hyper = list(prec = list(initial = 0, prior = \"pc.prec\",
                                  param = c(1, 0.05))))",
      "f(islandCurrent, model = \"iid\",
         hyper = list(prec = list(initial = 0, prior = \"pc.prec\",
                                  param = c(1, 0.05))))",
      "f(ID1, model = \"iid\",
         hyper = list(prec = list(initial = 0, prior = \"pc.prec\",
                                  param = c(1, 0.05))))",
      "f(ID2, values = as.numeric(colnames(innerCmatrix)), model = \"generic0\",
         Cmatrix = innerCmatrix, constr = TRUE,
         hyper = list(prec = list(initial = 0, prior = \"pc.prec\",
                                  param = c(1, 0.05))))",
      "f(ID3, values = as.numeric(colnames(outerCmatrix)), model = \"generic0\",
         Cmatrix = outerCmatrix, constr = TRUE,
         hyper = list(prec = list(initial = 0, prior = \"pc.prec\",
                                  param = c(1, 0.05))))",
      "f(ID4, values = as.numeric(colnames(otherCmatrix)), model = \"generic0\",
         Cmatrix = otherCmatrix, constr = TRUE,
         hyper = list(prec = list(initial = 0, prior = \"pc.prec\",
                                  param = c(1, 0.05))))")
```

```
# Add effects together
effects = c(effects, randomEffects)
# Make INLA formula for different phenotypic responses.
wingFormula = reformulate(effects, response = "wing")
massFormula = reformulate(effects, response = "mass")
tarsusFormula = reformulate(effects, response = "tarsus")
# Find inverse of the GRMs
innerCmatrix = solve(innerGRM)
outerCmatrix = solve(outerGRM)
otherCmatrix = solve(otherGRM)
# Function to transform precisions to variances in INLA-generated marginals
precToVar = function(marginal){
  sigmaMarg = inla.tmarginal(function(x) 1 / x, marginal)
  summaryStats = inla.zmarginal(sigmaMarg, silent = TRUE)[c(1, 5, 3, 7)]
  summaryStats = round(as.numeric(summaryStats), digits = 2)
  names(summaryStats) = c("mean", "mode", "2.5%", "97.5%")
  return(summaryStats)
}
# Function to add random effect variances to INLA model objects
addVariances = function(model){
  variances = do.call("rbind", lapply(model$marginals.hyperpar, precToVar))
  rownames(variances) = gsub("Precision", "Variance", rownames(variances))
  model$variances = variances
  return(model)
}
# Run INLA to fit each model. We rerun each model twice to increase stability.
# Also define the prior distribution of the residual effect variance here.
# Run INLA on wing length
wingModel1 =
  inla(formula = wingFormula, family = "gaussian", data = phenotypes,
       verbose = TRUE,
       control.family = list(hyper = list(prec = list(initial = 0,
                                                      prior = "pc.prec",
```

```
                                                       param = c(1, 0.05)))))
save(wingModel1, file = "wingModel.RData")
wingModel2 = inla.rerun(wingModel1)
resave(wingModel2, file = "wingModel.RData")
wingModel = inla.rerun(wingModel2)
wingModel = addVariances(wingModel)
resave(wingModel, file = "wingModel.RData")

# Run INLA on body mass
massModel1 =
  inla(formula = massFormula, family = "gaussian", data = phenotypes,
       verbose = TRUE,
       control.family = list(hyper = list(prec = list(initial = 0,
                                                       prior = "pc.prec",
                                                       param = c(1, 0.05)))))
save(massModel1, file = "massModel.RData")
massModel2 = inla.rerun(massModel1)
resave(massModel2, file = "massModel.RData")
massModel = inla.rerun(massModel2)
massModel = addVariances(massModel)
resave(massModel, file = "massModel.RData")

# Run INLA on tarsus length
tarsusModel1 =
  inla(formula = tarsusFormula, family = "gaussian", data = phenotypes,
       verbose = TRUE,
       control.family = list(hyper = list(prec = list(initial = 0,
                                                       prior = "pc.prec",
                                                       param = c(1, 0.05)))))
save(tarsusModel1, file = "tarsusModel.RData")
tarsusModel2 = inla.rerun(tarsusModel1)
resave(tarsusModel2, file = "tarsusModel.RData")
tarsusModel = inla.rerun(tarsusModel2)
```

```
tarsusModel = addVariances(tarsusModel)

resave(tarsusModel, file = "tarsusModel.RData")
```

#### S3: The house sparrow study metapopulation

The extended genomic genetic groups animal model was applied to a metapopulation of house sparrows living on islands in the Helgeland region in Northern Norway, which has been subject to individual-based long-term study since 1993 (see eg. Sæther *et al.*, 1999; Jensen *et al.*, 2008, 2013; Baalsrud *et al.*, 2014). The available data include phenotypic records of 1932 sparrows measured between 1993 and 2016, from eight islands that are known to have inter-island dispersal (Ranke *et al.*, 2021; Saatoglu *et al.*, 2021). Due to repeated measurements (every bird is recognizable through being banded with a metal leg ring with a unique number), we have 4625 records in total. Measured phenotypes include, among others, wing length (to the nearest millimeter), body mass (to the nearest 0.1 gram) and tarsus length (to the nearest 0.01 millimeter), and these phenotypes will be the traits of interest in our models. Some data is missing for each of the phenotypic traits, with wing length missing for 131 records, body mass missing for 250 and tarsus length missing for 126.

Large-scale genotyping of house sparrows from the study system has been performed by taking blood samples from each individual, extracting DNA from the blood and genotyping the samples on a custom Axiom array with probes for 200 000 SNPs, as described in Lundregan *et al.* (2018). After quality control, filtering for minor allele frequency ($> 0.01$), SNP call rate ($> 0.9$), individual call rate ($> 0.95$) and only retaining SNPs assigned to a chromosome in the house sparrow reference genome (Elgvin *et al.*, 2017), 181 354 genotyped SNPs (with some missing genotypes) in 3032 individuals (including the 1932 phenotyped individuals) remained.

Subpopulations of sparrows located on the eight different islands vary in environmental conditions such as habitat, buffering against bad weather, and population density (Pärn *et al.*, 2012; Baalsrud *et al.*, 2014; Muff *et al.*, 2019; Niskanen *et al.*, 2020; Araya-Ajoy *et al.*, 2021). The populations on five islands closer to the mainland reside mostly in colonies on dairy farms where the birds can hide inside farm buildings during bad weather and enjoy more stable access to food but sparrow densities are higher compared to populations on the three islands further out to sea, where birds mostly live scattered in local people's gardens and are outside all year round. To account for possible genetic differences between these subpopulations originating from different island groups, we partitioned the study population into genetic groups, where each group is associated with a set of islands. The former group of islands we label as the `inner` genetic group (encoded as 1), and the latter group as the `outer` genetic group (encoded as 2). Sparrows from other islands in the study system (that were not systematically SNP-genotyped) are also present in the data set due to dispersal, and we place these sparrows in a final genetic group `other` (encoded as 3).

The pedigree used in the pedigree-based genetic groups animal model (Muff *et al.*, 2019) was constructed from the genomic data using the R package SEQUOIA (Huisman, 2017). The pedigree-based global ancestries were derived using the R package nadiv (Wolak, 2012), as described in Niskanen *et al.* (2020).

To perform the local ancestry inference, we had to assign some individuals as purebred in the genetic groups of interest, since methods for local ancestry inference in admixed populations generally rely on reference panels of representative (*i.e.*, purebred) individuals from each group. In most local ancestry inference methods these reference panels are used as templates for what the genomes of individuals from that group generally look like (Geza *et al.*, 2019), and they are used to assign regions of each admixed individual's genome as descended from a specific group. We assigned any individual that has both parents missing in the pedigree as a purebred in one of the groups for the genomic model. This approach ensures that the starting condition of the genomic genetic groups model is as similar as possible to the corresponding pedigree-based genetic groups model (Muff *et al.*, 2019) and hence allows a valid comparison between pedigree-based and genomic genetic group models. Which genetic group purebred individuals were assigned to depended on information about their natal island as determined either from ecological data or genetic assignment (Kuismin *et al.*, 2020; Saatoglu *et al.*, 2021). For $< 5\%$ of the genotyped purebred individuals the natal island was unknown. For these individuals, the earliest island of observation was used, as this is their most likely natal island (Saatoglu *et al.*, 2021). Conversely, individuals with at least one known parent in the pedigree were considered admixed in the genomic model.

### S4: Additional results

#### S4.1: Homogeneous-variance models

We fit genetic group models with homogeneous additive genetic variances (Wolak & Reid, 2017). Thus, the genetic groups share a single $V_{\text{A}}$ and differ only in their genetic value means. Apart from the homogeneous $V_{\text{A}}$, they are otherwise identical to the models fitted in the main text. Here we use three different genetic relatedness matrices to this end: the standard pedigree-based relatedness matrix $\mathbf{A}$, the widely used GRM by VanRaden (2008) defined in Equation (1) in the main text and finally a one-group ($R = 1$) version of our GRM (defined in the main text) which uses single allele data instead of genotype data. The entries of the latter GRM are thus defined as

$$G_{ij} = \frac{\sum_{m=1}^{M} \sum_{h=1}^{2} \sum_{h'=1}^{2} \left( w_{im}^{(h)} - \widehat{p}_m \right) \left( w_{jm}^{(h')} - \widehat{p}_m \right)}{2 \sum_{m=1}^{M} \widehat{p}_m \left(1 - \widehat{p}_m\right)} \ . \tag{S13}$$

A further difference between the models using Equation (1) and Equation (S13) is that in the latter model the missing genotype data was imputed. The purpose of using this final GRM is to show that the move from genotypes to single alleles (gametic phasing) and the imputation of missing genotypes has little impact on the model results.

The estimated posterior distributions for $V_{\mathrm{A}}$ in each of these different models are shown in Fig. S1. Since the results from the two genomic models almost completely overlap for all three phenotypes, it seems that using single alleles rather than genotypes (the result of gametic phasing) has little impact, nor does the imputation of missing genotypes in this data set. We also note that the differences between the genomic models and the pedigree-based models are similar to the differences seen in the genetic group-models with heterogeneous additive genetic variances (in the main text). This is implies that the discrepancies between genomic and pedigree-based results do not stem from the design of the genomic genetic groups model (moving from considering genotypes to single alleles), the gametic phasing or the imputation of missing genotypes. Rather, the differences stem from inherent differences in the genomic and pedigree-based approaches.

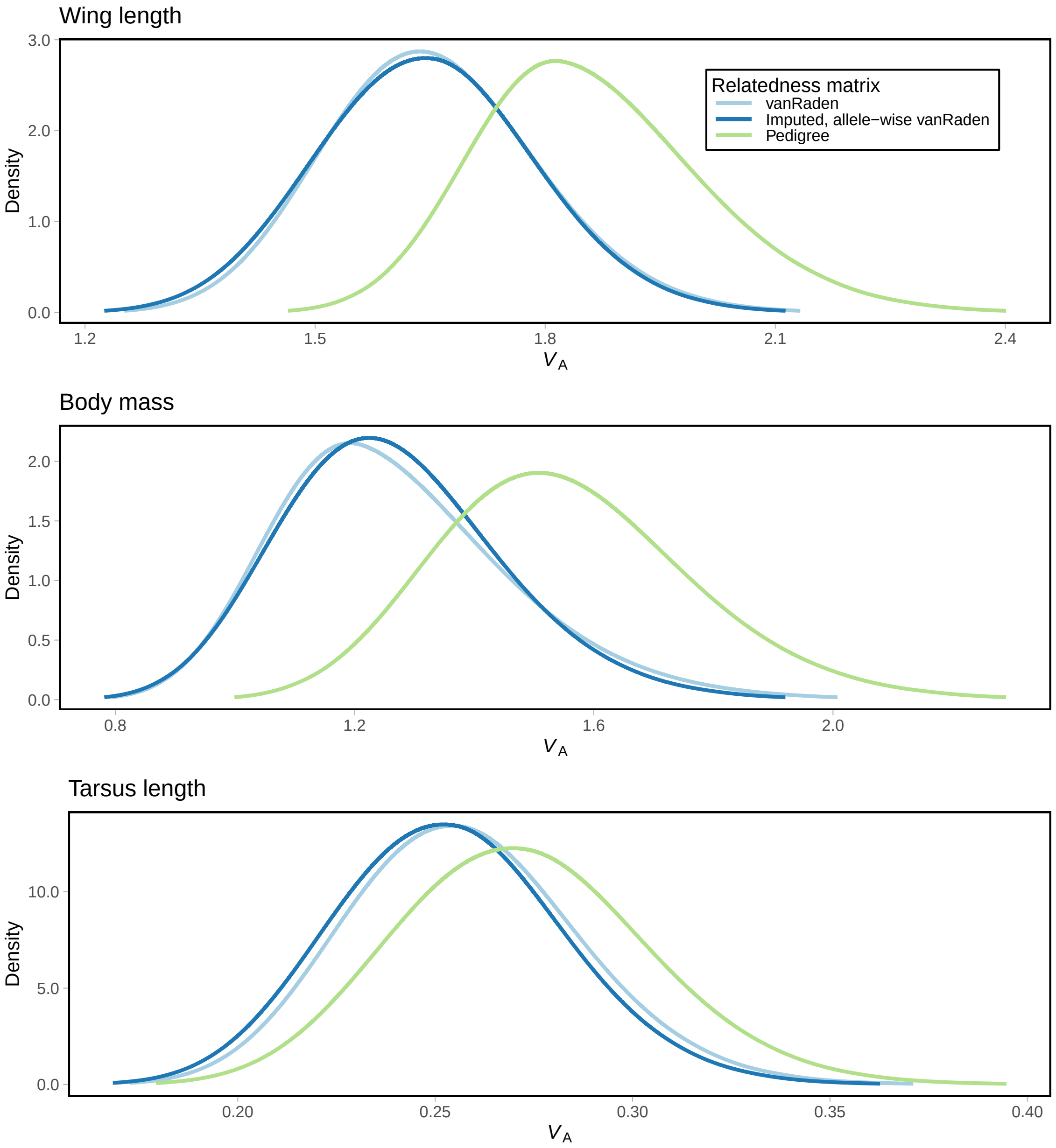

Figure S1: The estimated posterior distribution of additive genetic variances in the models for wing length (top), body mass (middle) and tarsus length (bottom). The pedigree-based model posteriors are shown in lime, while the genomic model posteriors are blue, with light blue indicating the standard genotype-based GRM and dark blue indicating the single-allele GRM with imputed genotypes.

#### S4.2: Two-group model including segregation variance

In addition to additive genetic variance, an further source of genetic variance is induced for admixed individuals, namely the segregation variance, which is caused by group differences in the level of linkage disequilibrium (LD) and allele effects (Slatkin & Lande, 1994). Such variances can be large when considering admixture in breeding scenarios (see e.g. Rio *et al.*, 2020), or when the number of loci impacting the phenotypes is very low. Since segregation variances occur between all combinations of groups, $R(R-1)/2$ segregation variances must be estimated in the presence of $R$ genetic groups. Thus, models that include segregation variances (such as Lo *et al.*, 1993; Cantet & Fernando, 1995; García-Cortés & Toro, 2006) rapidly grow computationally cumbersome as $R$ increases, and require much statistical power to fit. As we mention in the main text, segregation variances will be small given that the infinitesimal model holds.

Here, we fit models on our data set which include segregation variances to test the assumption that segregation variance is negligible. However, including segregation variances in the model with three genetic groups would add three random effect terms, and put too much strain on the available data. Furthermore, the segregation covariance matrices $\mathbf{S}_{rr'}$ were not positive-definite in the three-group model, and were thus not proper covariance matrices. We thus fit model with only two genetic groups, inducing only one segregation term $g_i^{(12)}$. This was accomplished by folding the `other` group into the `inner` group, by letting `other` purebreds be considered `inner` purebreds instead. Let $\boldsymbol{g}^{(12)} \sim \mathsf{N}\left(\mathbf{0}, \sigma^2_{\mathrm{S}_{12}}\mathbf{S}_{12}\right)$, where $(\mathbf{S}_{12})_{ij} = -\frac{1}{2}\left(\widehat{\Delta}^{(12)}_{ij} + \widehat{\Delta}^{(21)}_{ij}\right)$ as found in equation (S12), Supporting Information S1. Using equations (S10) and (S11) in Supporting Information S1, $-\frac{1}{2}\left(\Delta^{(12)}_{ij} + \Delta^{(21)}_{ij}\right)$ can be rewritten $\Delta^{(1)}_{ij}$, the same result found by Rio *et al.* (2020). We thus estimate the $ij^{\mathrm{th}}$ entry of the segregation covariance matrix by $\widehat{\Delta}^{(1)}_{ij} = \widehat{\theta}^{(1)}_{ij} - \widehat{\pi}_{i1}\widehat{\pi}_{j1}$. Segregation variances were given penalized complexity priors PC(0.1, 0.01). We also fit the same two-group model, but with no segregation term, to see the impact of its inclusion on the other model parameters. Aside from the combination of the `inner` and `other` groups and the inclusion of segregation variance, these models are identical to the models in the main text.

The results from the models are found in Tables S1 and S2, and the model posteriors of the additive genetic variances are shown in Fig. S2. The segregation variances are very small for all phenotypes. Furthermore, and their inclusion have little impact on the remaining model parameters. The largest difference is seen for the wing length in the `inner` group, but here we see that in fact the posterior distribution is shifted to slightly larger values in the model *with* segregation variance and the HPD CIs are almost identical – so it is not so that the segregation terms have “absorbed” any variance from other variance components. Regardless, given the uncertainties in the posteriors, this difference is negligible. Thus, we surmise that considering segregation to be negligible is reasonable, at least in the case of this data.

Table S1: Posterior statistics for the fixed effects of the 2-group genetic group animal models, as derived from models with or without segregation variance, for the three investigated phenotypic traits. For each parameter, the posterior mean is reported in the first row, and the 95% HPD CI in the second row. The parameters denote the effects of being female compared to male ($\beta_{\text{sex}}$), inbreeding ($\beta_{F_{\text{GRM}}}$), month ($\beta_{\text{month}}$) and age ($\beta_{\text{age}}$). The genetic group effect $\gamma_2$ of `outer` denotes the effect of being purebred in this group relative to `inner`.

| | Wing length | | Body mass | | Tarsus length | |
|---|---|---|---|---|---|---|
| Parameter | W/ seg. | W/o seg. | W/ seg. | W/o seg. | W/ seg. | W/o seg. |
| $\beta_{\text{sex}}$ | -2.76 | -2.76 | 0.48 | 0.48 | -0.08 | -0.08 |
| | (-2.89, -2.63) | (-2.89, -2.63) | (0.30, 0.65) | (0.30, 0.65) | (-0.14, -0.01) | (-0.14, -0.01) |
| $\beta_{F_{\text{GRM}}}$ | -1.70 | -1.69 | -0.84 | -0.85 | -0.69 | -0.70 |
| | (-3.23, -0.17) | (-3.22, -0.16) | (-2.92, 1.23) | (-2.93, 1.23) | (-1.51, 0.12) | (-1.51, 0.11) |
| $\beta_{\text{month}}$ | -0.19 | -0.19 | -0.30 | -0.30 | 0.03 | 0.03 |
| | (-0.23, -0.15) | (-0.23, -0.15) | (-0.36, -0.24) | (-0.36, -0.24) | (0.02, 0.04) | (0.02, 0.04) |
| $\beta_{\text{age}}$ | 0.46 | 0.46 | 0.09 | 0.09 | -0.00 | -0.00 |
| | (0.42, 0.49) | (0.42, 0.50) | (0.03, 0.15) | (0.03, 0.15) | (-0.01, 0.00) | (-0.01, 0.00) |
| $\gamma_2$ | -0.30 | -0.31 | -0.78 | -0.79 | 0.08 | 0.08 |
| | (-0.87, 0.27) | (-0.87, 0.26) | (-1.47, -0.11) | (-1.47, -0.11) | (-0.09, 0.26) | (-0.09, 0.26) |

Table S2: Posterior statistics for the random effect variances of the 2-group genetic group animal models, as derived from models with or without segregation variance, for the three investigated phenotypic traits. For each parameter, the posterior mode and mean (formatted mode;mean) are reported in the first row, and the 95% HPD CI in the second row. The parameters denote variance explained by year of measurement ($\sigma^2_{\text{year}}$), island of measurement ($\sigma^2_{\text{island}}$), permanent environmental effects ($\sigma^2_{\text{ID}}$), group-specific $V_{\text{A}}$ ($\sigma^2_{\text{G}_r}$, $r = 1$ for `inner`, $r = 2$ for `outer`), segregation variance between `inner` and `outer` ($\sigma^2_{\text{S}_{12}}$) and residual environmental effects ($\sigma^2_{\varepsilon}$).

| | Wing length | | Body mass | | Tarsus length | |
|---|---|---|---|---|---|---|
| Parameter | W/ seg. | W/o seg. | W/ seg. | W/o seg. | W/ seg. | W/o seg. |
| $\sigma^2_{\text{year}}$ | 0.05;0.06 | 0.05;0.06 | 0.04;0.04 | 0.04;0.05 | 0.01;0.01 | 0.01;0.01 |
| | (0.02, 0.16) | (0.02, 0.14) | (0.01, 0.10) | (0.01, 0.14) | (0.00, 0.04) | (0.00, 0.03) |
| $\sigma^2_{\text{island}}$ | 0.08;0.10 | 0.08;0.10 | 0.12;0.15 | 0.11;0.15 | 0.00;0.00 | 0.00;0.01 |
| | (0.02, 0.31) | (0.02, 0.29) | (0.03, 0.43) | (0.02, 0.51) | (0.00, 0.02) | (0.00, 0.03) |
| $\sigma^2_{\text{ID}}$ | 0.44;0.44 | 0.46;0.46 | 1.14;1.15 | 1.12;1.13 | 0.36;0.36 | 0.36;0.36 |
| | (0.32, 0.61) | (0.31, 0.62) | (0.87, 1.48) | (0.86, 1.49) | (0.32, 0.40) | (0.32, 0.40) |
| $\sigma^2_{\text{G}_1}$ | 1.58;1.59 | 1.54;1.56 | 1.11;1.12 | 1.11;1.12 | 0.29;0.29 | 0.29;0.29 |
| | (1.32, 1.92) | (1.28, 1.95) | (0.79, 1.55) | (0.78, 1.54) | (0.23, 0.36) | (0.23, 0.36) |
| $\sigma^2_{\text{G}_2}$ | 2.07;2.10 | 2.10;2.15 | 2.37;2.43 | 2.40;2.45 | 0.02;0.03 | 0.03;0.04 |
| | (1.34, 3.03) | (1.42, 3.15) | (1.40, 3.82) | (1.39, 3.81) | (0.00, 0.12) | (0.00, 0.13) |
| $\sigma^2_{\text{S}_{12}}$ | 0.05;0.07 | | 0.01;0.02 | | 0.01;0.03 | |
| | (0.01, 0.20) | | (0.00, 0.07) | | (0.00, 0.15) | |
| $\sigma^2_{\varepsilon}$ | 0.98;0.98 | 0.98;0.98 | 2.89;2.89 | 2.89;2.89 | 0.02;0.02 | 0.02;0.02 |
| | (0.93, 1.04) | (0.93, 1.03) | (2.73, 3.05) | (2.73, 3.06) | (0.02, 0.02) | (0.02, 0.02) |

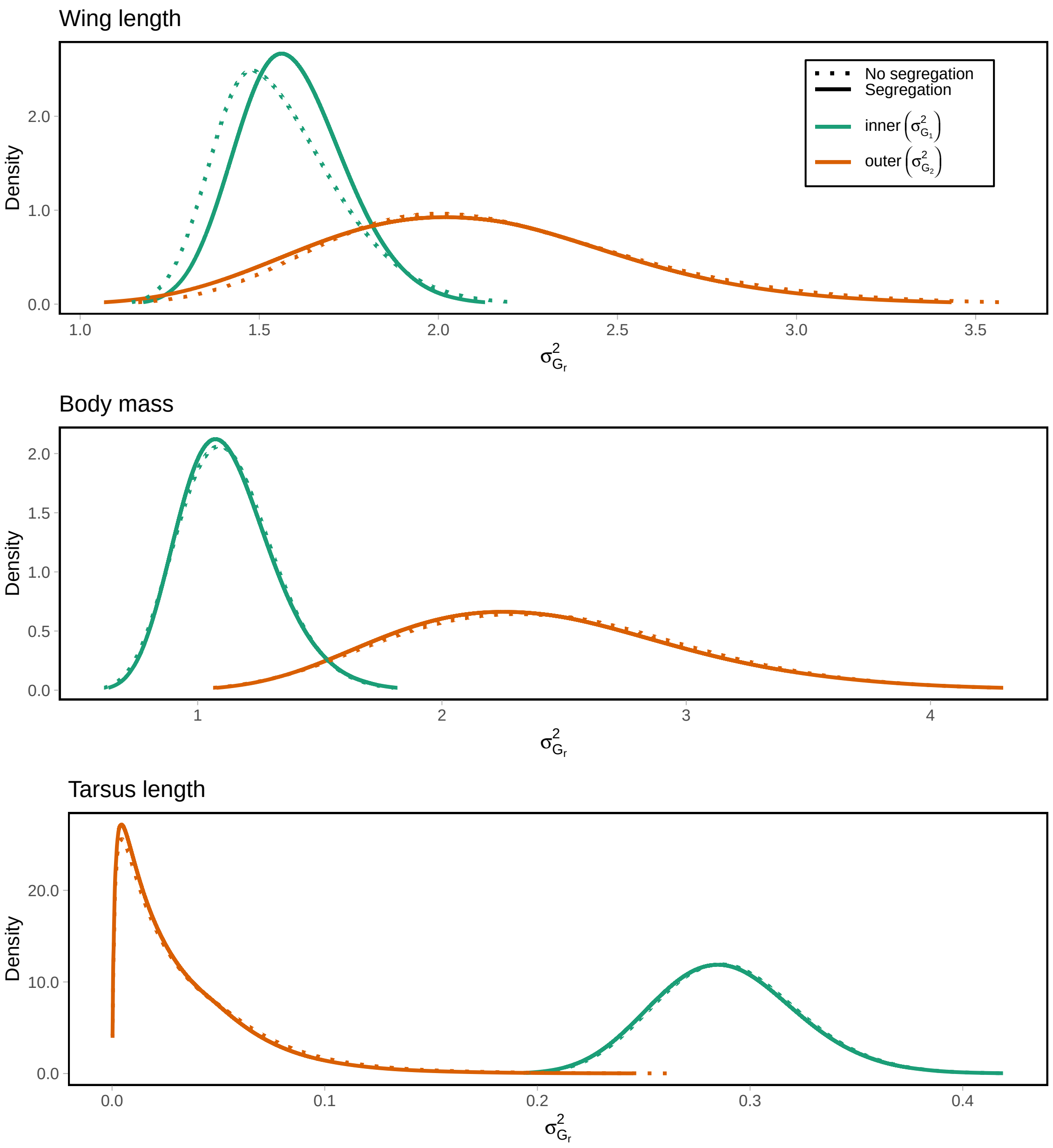

Figure S2: The posterior distribution of group-specific additive genetic variances and segregation variance in the models for wing length (top), body mass (middle) and tarsus length (bottom). Posterior variances for the different genetic groups are shown in different colors. Posteriors from the models containing segregation variance have solid lines, whereas the no-segregation model posteriors are shown with dotted lines.

#### S4.3: Group-differences in allele frequencies

We present here the differences in allele frequencies between all combinations of genetic groups (Fig. S3).

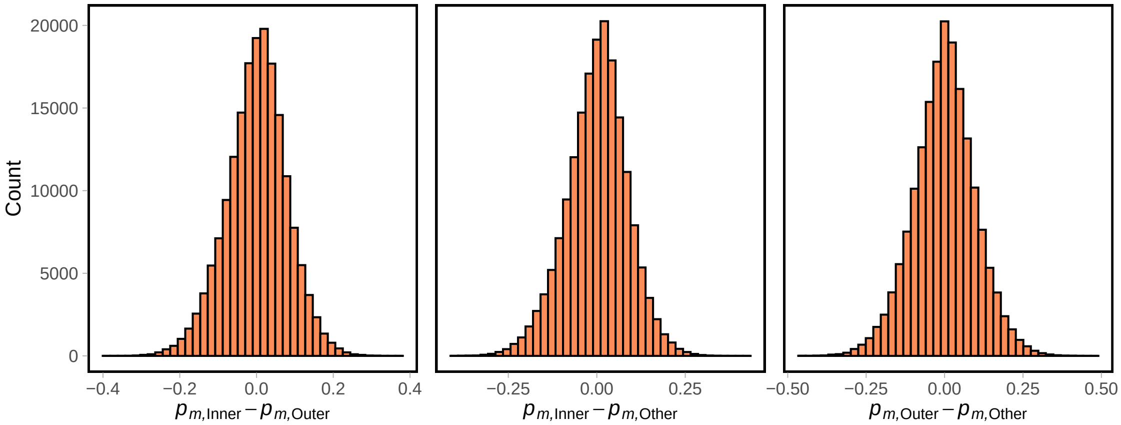

Figure S3: Distribution of group-differences in allele frequencies of the alternate allele between each pair of genetic groups.
